# Identification of PTGR2 inhibitors as a new therapy for diabetes and obesity

**DOI:** 10.1101/2024.12.17.629058

**Authors:** Yi-Cheng Chang, Meng-Lun Hsieh, Hsiao-Lin Lee, Siow-Wey Hee, Chi-Fon Chang, Hsin-Yung Yen, Yi-An Chen, Yet-Ran Chen, Ya-Wen Chou, Fu-An Li, Yi-Yu Ke, Shih-Yi Chen, Ming-Shiu Hung, Alfur Fu-Hsin Hung, Jing-Yong Huang, Chu-Hsuan Chiu, Shih-Yao Lin, Sheue-Fang Shih, Chih-Neng Hsu, Juey-Jen Hwang, Teng-Kuang Yeh, Ting-Jen Rachel Cheng, Karen Chia-Wen Liao, Daniel Laio, Shu-Wha Lin, Tzu-Yu Chen, Chun-Mei Hu, Ulla Vogel, Daniel Saar, Birthe B. Kragelund, Lun Kelvin Tsou, Yu-Hua Tseng, Lee-Ming Chuang

## Abstract

Peroxisome proliferator-activated receptor γ (PPARγ) is a master transcriptional regulator of systemic insulin sensitivity and energy balance. The anti-diabetic drug thiazolidinediones (TZDs) are potent synthetic PPARγ ligands with undesirable side effects, including obesity, fluid retention, and osteoporosis. 15-keto prostaglandin E2 (15-keto-PGE2) is an endogenous PPARγ ligand metabolized by prostaglandin reductase 2 (PTGR2). Here, we confirmed that 15-keto-PGE2 binds and activates PPARγ via covalent binding. In patients with type 2 diabetes and obese mice, serum 15-keto-PGE2 levels were decreased. Administration of 15-keto-PGE2 improves glucose homeostasis and prevented diet-induced obesity in mice. Either genetic inhibition of PTGR2 or PTGR2 inhibitor BPRPT0245 protected mice from diet-induced obesity, insulin resistance, and hepatic steatosis without fluid retention and osteoporosis.In conclusion, inhibition of PTGR2 is a new therapeutic approach to treat diabetes and obesity through increasing endogenous PPARγ ligands without side effects of synthetic PPARγ ligands TZDs.

## Introduction

Insulin resistance is the pathological feature of type 2 diabetes mellitus with obesity as the main contributor. Peroxisome proliferator-activated receptor γ (PPARγ) is a master transcription regulator of insulin sensitivity, lipid metabolism, thermogenesis and inflammation (Ahmadian *et al*., 2013). Thiazolidinediones (TZDs) are synthetic full PPARγ agonists and potent insulin sensitizers clinically used for treating type 2 diabetes. However, TZD use has several undesirable side effects including weight gain, osteoporosis, and fluid retention, which limit their clinical application (Soccio *et al*, 2014). Recent clinical trials of sodium-glucose cotransporter 2 (SGLT2) inhibitors and glucagon-like peptide 1 (GLP-1) analogs have demonstrated that their cardiovascular benefits derive mainly from weight loss (Wiviott *et al*., 2019; Marso *et al*., 2016). In addition, among all current anti-diabetic drugs, only PPARγ agonists and metformin are insulin-sensitizer. Therefore, novel approaches to activate PPARγ without side effects of TZDs are eagerly awaited (DePaoli *et al*., 2014; Choi *et al*., 2011; Bruning *et al*., 2007; Waki *et al*., 2007).

We and other groups previously demonstrated that the polyunsaturated fatty acid 15-keto-prostaglandin E2 (15-keto-PGE2) is a natural endogenous PPARγ ligand derived from PGE2 and has minimal binding affinity for prostanoid receptors (Chou *et al*., 2007; Waku *et al*., 2009; Shiraki *et al*., 2005; Harmon *et al*., 2010). 15-keto-PGE2 is further catalyzed by prostaglandin reductase 2 (PTGR2) to become inactive metabolite 13,14-dihydro-15-keto-PGE2 (Chou *et al*., 2007). In this study, we sought to inhibit PTGR2 to increase endogenous PPARγ ligands for treating diabetes instead of using synthetic PPARγ ligand TZDs.

We demonstrated that 15-keto-PGE2 levels are reduced in obese/insulin-resistant mice and human subjects with type 2 diabetes. Direct administration of 15-keto-PGE2 improved glucose homeostasis and prevented diet-induced obesity without fluid retention. Either genetic or pharmacological inhibition of PTGR2 prevented diet-induced obesity, improved insulin sensitivity, glucose tolerance, and ameliorated hepatic steatosis without fluid retention or osteoporosis.

## Results

### 15-keto-PGE2 increased insulin-stimulated glucose uptake and activated PPARγ through covalent binding to the cysteine residue of PPARγ

Previous studies have shown that the polyunsaturated fatty acid 15-keto-PGE2 is a natural endogenous PPARγ ligand (Choi *et al*, 2011; Waku *et al*, 2009; Harmon *et al*, 2010). 15-keto- PGE2 is further catalyzed by PTGR2 to become inactive metabolite 13,14-dihydro-15-keto-PGE2 (Fig 1A). To validate this, we incubated 15-keto-PGE2 with recombinant human PTGR2 protein and found that 99.83 % of 15-keto-PGE2 is rapidly converted 13,14-dihydro-15-keto-PGE2 (Appendix Fig S1). Using the Gal4-PPARγ/UAS-LUC reporter assay system, we discovered that 15-keto-PGE2 enhanced the transactivation activity of PPARγ in a dose-dependent manner (Fig 1B). Expression of *Glut4* (Fig 1C) and other PPARγ-downstream genes involved in insulin signaling (*Irs2, Sorbs1*) (Fig 1D and E), lipid metabolism (*Cd36, Acs*) (Fig 1F and G), and adipogenesis (*Cebpa, Adipq*) (Fig 1H and I) were significantly increased in 15-keto-PGE2-treated 3T3-L1 adipocytes, supporting that 15-keto-PGE2 enhanced the transactivation activity of PPARγ. Furthermore, we found that 15-keto-PGE2 increased insulin-stimulated glucose uptake in induced 3T3-L1 adipocytes dose-dependently, while 13,14-dihydro-15-keto-PGE2 showed little effect (Fig 1J). Of note, the maximal dose of exogenously added 15-keto-PGE2 (10-20 μM) in 3T3-L1 adipocytes resulted in an approximately ∼1.25 to 1.78-fold increase of the physiological intracellular 15-keto-PGE2 levels, suggesting that near-physiological intracellular 15-keto-PGE2 levels are sufficient to increase insulin-stimulated glucose uptake (Fig EV1).

**Figure 1.**
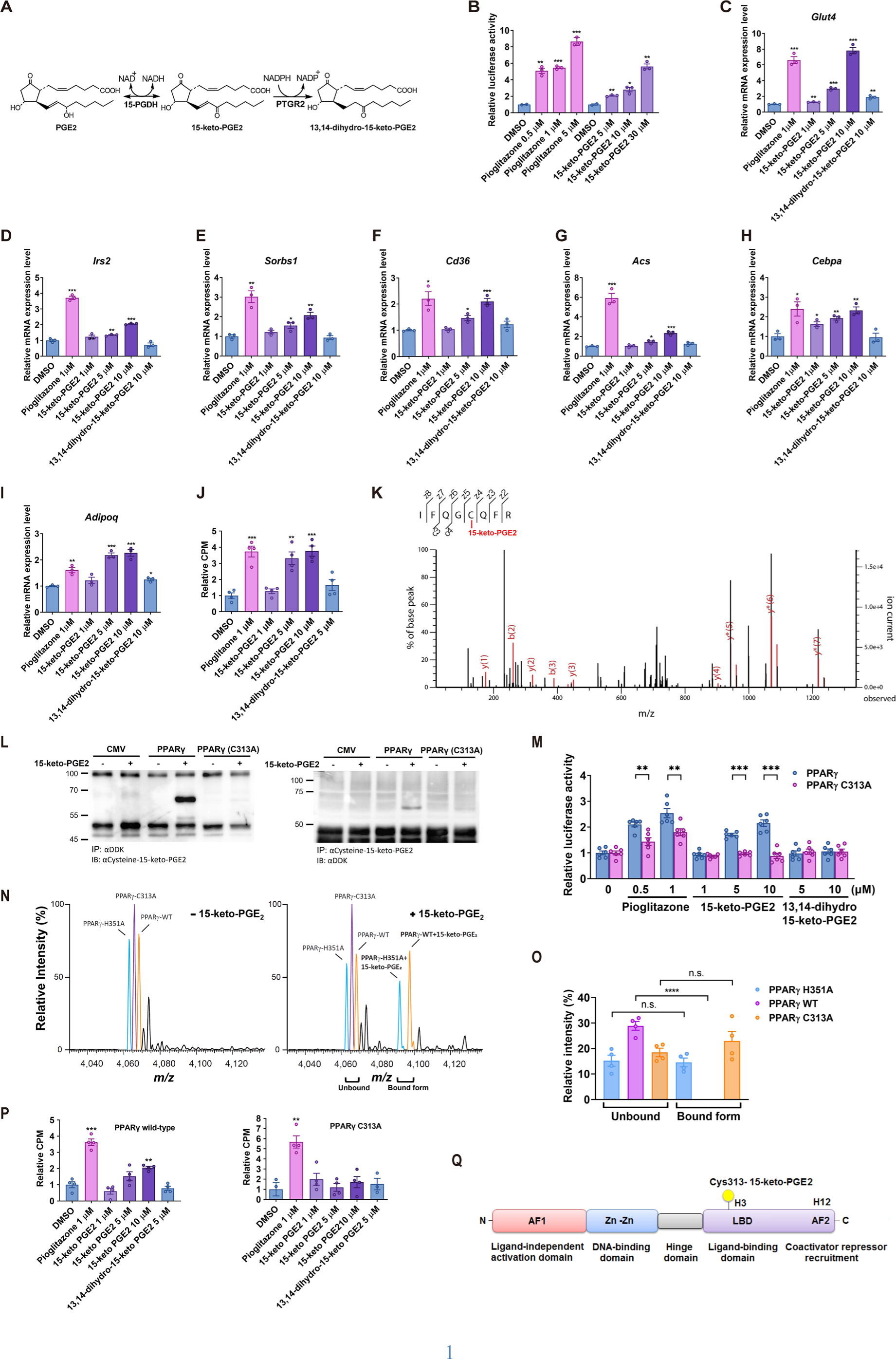
15-keto-PGE2 activates murine PPARγ through binding to cysteine 313 residue. (A) Metabolism of 15-keto-PGE2. (B) Activation of murine PPARγ (mPPARγ) measured by Gal-PPARγ/UAS-LUC reporter assay in HEK293T cells (n=3 per group). Cells were transfected with Gal4-PPARγ, UAS-LUC, and TK-Rluc (Renilla luciferase), and treated with pioglitazone or 15-keto-PGE2. RT-qPCR of (C) *Glut4* and other PPARγ-downstream genes including (D) *Irs2*, (E) *Sorbs1*, (F) *Cd36,* (G) *Acs*, (H) *Cepba*, and (I) *Adipoq* in induced 3T3-L1 adipocytes treated with 15-keto- PGE2 (n=3 per group). (J) Effect of 15-keto-PGE2 on insulin-stimulated glucose uptake in induced 3T3-L1 adipocytes (n=4 per group). (K) HEK293T cells transfected by mPPARγ and treated with 15-keto-PGE2. Covalent binding of 15-keto-PGE2 to mPPARγ detected by liquid-chromatography tandem mass spectrometry (LC-MS/MS). (L) Reciprocal co-immunoprecipitation of mPPARγ and cysteine-15-keto-PGE2. Myc-DDK- mPPARγ and Myc-DDK-mPPARγ C313A were expressed in HEK293T cells, and immunoprecipitation (IP) conducted using either anti-DDK (anti-Flag) or anti-15-keto-PGE2- cysteine-BSA antibody, followed by immunoblotting with anti-15-keto-PGE2-cysteine-BSA and anti-DDK antibody. (M) PPRE reporter activity after addition of 15-keto-PGE2 to HEK293T cells transfected with wild-type and C313A mutant mPPARγ. (N) Native mass spectrometry spectrum showed the binding of 15-keto-PGE2 to wild-type and mPPARγ mutants (C313A and H351A). The spectrum of unbound free-form proteins was shown in the left panel. The spectrum of bound form after the addition of 15-keto-PGE2 was shown in the right panel. (n=4:4) and (O) histogram. (P) 15-keto-PGE2 enhanced insulin-stimulated glucose uptake in PPARγ-null 3T3-L1 clones (#1296) rescued with wild-type mPPARγ but not in those rescued with mutant mPPARγ (C313A) (n=4 per group). (Q) Diagram showing the motifs of mPPARγ and 15-keto-PGE2 binding site. Data information: All data are presented as mean and standard error (S.E.M.). Statistical significance was calculated by one-way analyses of variance (ANOVA) with Tukey’s post hoc test in (B-J, O, P) and two-sample independent *t*-test in (M). \**p* < 0.05, ** *p* < 0.01, *** *p* < 0.001, **** *p* < 0.0001. n.s. means no statistical difference. Source data are available online for this figure.

To explore how 15-keto-PGE2 activated PPARγ, murine PPARγ (mPPARγ) was overexpressed and then incubated with 15-keto-PGE2. Cell lysates were then separated by SDS-PAGE and stained with Coomassie blue. The protein band around 60 kDa was excised from the gel, followed by digested in gel. The peptides were further analyzed using LC-MS/MS. The results showed that 15-keto-PGE2 was covalently coupled to mPPARγ at the Cys313 residue (Fig 1K). To confirm the data obtained from LC-MS/MS, we performed reciprocal immunoprecipitations to demonstrate the interaction between 15-keto-PGE2 and mPPARγ. We generated a monoclonal antibody against 15-keto-PGE2 conjugated to the cysteine residues of bovine serum albumin (BSA). HEK293T cells were transfected with Myc-DDK-tagged mPPARγ and treated with 15-keto-PGE2. Cell lysates were first immunoprecipitated with anti-FLAG antibody and then immunoblotted with monoclonal anti-15-keto-PGE2-cysteine-BSA antibody. A single band of PPARγ (57 kDa) was detected in samples expressing wild-type mPPARγ but not in samples expressing mPPARγ with C313A mutation (Fig 1L, left panel). Reciprocally, when protein lysates were immunoprecipitated with anti-15-keto-PGE2-cysteine-BSA antibody and then immunoblotted with anti-FLAG antibody, a band of approximately 57 kDa was detected only in cells overexpressing wild-type mPPARγ but not mPPARγ with C313A mutation (Fig 1L, right panel). We then expressed wild-type mPPARγ or mPPARγ with the C313A mutation in HEK 293T cells transfected with mPPARγ transactivation reporter (PPRE). The addition of 15-keto-PGE2 dose-dependently increases the reporter activity of mPPARγ, which was abolished in cells expressing mutant mPPARγ (C313A), suggesting that 15-keto-PGE2 activates mPPARγ through binding the Cys313 residue (Fig 1M).

We further validate the interaction between 15-keto-PGE2 and mPPARγ in mouse fat tissue using reciprocal co-immunoprecipitation. Lysates from the epididymal white adipose tissues of *Ptgr2* wild-type (*Ptgr*2^+/+^) and *Ptgr2* knockout (*Ptgr*2^-/-^) mice were immunoprecipitated with a mouse anti-15-keto-PGE2-cysteine-BSA antibody and then immunoblotted with a rat anti-PPARγ antibody. A single band corresponding to PPARγ (57 kDa) was detected (upper panel, Appendix Fig S2), confirming the covalent binding of mPPARγ to 15-keto-PGE2. However, when protein lysates were reciprocally immunoprecipitated with the anti-PPARγ antibody and immunoblotted with the anti-15-keto-PGE2-cysteine-BSA antibody, the mPPARγ band was obscured by the IgG heavy chain, which is enriched in tissue (lower panel, Appendix Fig S2).

To directly analyze the effect of Cys313 mutation on the binding of 15-keto-PGE2 to mPPARγ, we applied near-atomic high-resolution native mass spectrometry and performed ligand competition assay using wild-type mPPARγ and two mPPARγ mutants, including C313A and H351A, was applied to directly analyze the impact of these mutations on 15-keto-PGE2-binding to mPPARγ. The His351 residue adjacent to the Cys313 residue is important for the binding of pioglitazone to mPPARγ (Chhonker *et al*, 2021; Yamada *et al*, 2015). The mixture of wild-type mPPARγ and two mutants was well resolved (Fig 1N, left panel and Appendix Fig S3) and the binding assay with 15-keto-PGE2 was performed simultaneously. The resulting native mass spectrum showed almost complete disruption of 15-keto-PGE2-binding by C313A mutation whereas H351A had only minor impact on the binding of 15-keto-PGE2 of mPPARγ (Fig 1N, right panel, and Fig 1O). These data confirmed the critical role of the Cys313 residue for the binding for 15-keto-PGE2-binding to mPPARγ.

To further examine the role of mPPARγ Cys313 in 15-keto-PGE2-induced glucose uptake of cultured adipocytes, insulin-stimulated glucose uptake assay was conducted in mPPARγ-null 3T3- L1 cells expressing either wild-type or C313A mutant mPPARγ. mPPARγ-null 3T3-L1 cells were generated using CRISPR/Cas9 genome editing technique. Two mPPARγ-null 3T3-L1 clones were tested: #1296 was generated using sgRNA#1, and #2328 was generated using sgRNA#2 (Fig EV2A). Wild-type and C313A mutant mPPARγ were introduced to PPARγ-null 3T3-L1 clones, which were then induced to mature adipocytes (Fig EV2A, 2B). We found that 15-keto-PGE2 enhanced insulin-stimulated glucose uptake in clone # 1296 when rescued with wild-type mPPARγ (Fig 1P) but not when rescued with mutant mPPARγ (C313A) (Fig 1P) Similar findings were also observed in another mPPARγ-null 3T3-L1 clone #2328 reconstituted with wild-type or C313A mutant mPPARγ (Fig EV2C). Taken together, these results revealed that the insulin-sensitizing effect 15-keto-PGE2 is mediated through mPPARγ via binding to Cys313 (Fig 1Q).

### 15-keto-PGE2 levels were decreased in patients with type 2 diabetes and insulin-resistant/obese mice

In humans, the serum level of 15-keto-PGE2 was significantly reduced by ∼63 % in 24 individuals with type 2 diabetes compared with 24 age- and sex-matched non-diabetic controls (Fig 2A). In 50 non-diabetic humans, serum level of 15-keto-PGE2 was inversely correlated with the Homeostasis Model Assessment of Insulin resistance (HOMA-IR) index (*r*=-0.37, *P*=0.007) (Fig 2B), fasting glucose (*r*=-0.31, *P*= 0.02) (Fig 2C), and fasting insulin (*r*=-0.33, *P*= 0.02) (Appendix Fig S4). These findings showed the inverse association of endogenous PPARγ ligand 15-keto- PGE2 levels with insulin sensitivity and glucose homeostasis in humans. Consistently, serum levels of 15-keto-PGE2 were markedly reduced by ∼56 % in high-fat high-sucrose diet (HFHSD)- induced obese mice compared with chow-fed lean mice (Fig 2D). Similar decrease of 15-keto- PGE2 content in inguinal fat and perigonadal fat were similarly reduced (∼53% and ∼56%, respectively) in diet-induced obese mice compared with controls (Fig 2E and F).

**Figure 2.**
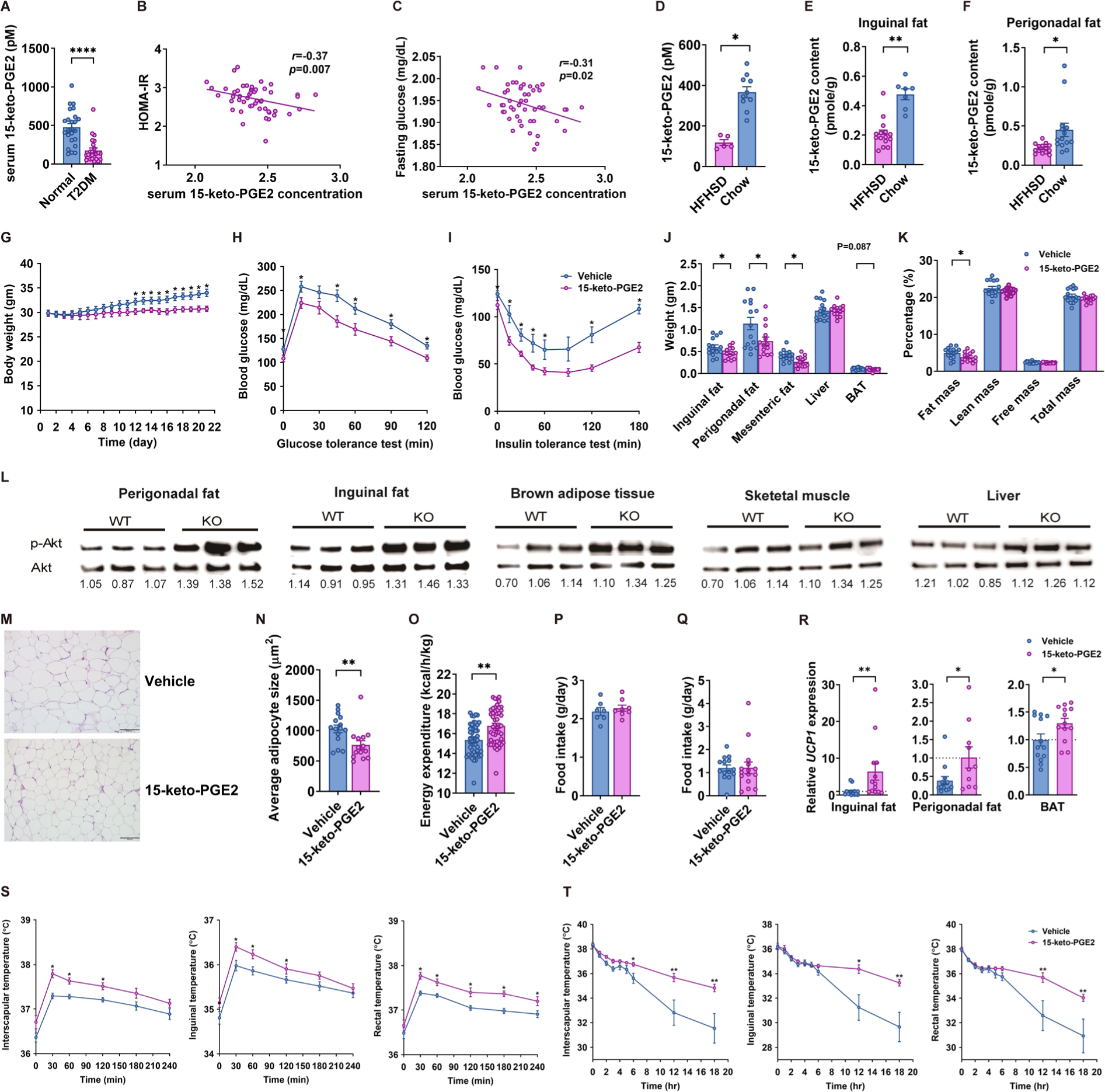
15-keto-PGE2 prevents diet-induced obesity and improves glucose homeostasis in mice. (A) Serum 15-keto-PGE2 concentration of non-diabetic human subjects and patients with type 2 diabetic patients(n=24:24). Correlation of serum 15-keto-PGE2 with (B) homeostasis model assessment of insulin resistance (HOMA-IR) and (C) fasting glucose with serum 15-keto-PGE2 levels in 50 non-diabetic human subjects. (D) Serum 15-keto-PGE2 concentration (n=5:10) and (E) 15-keto-PGE_2_ content in inguinal fat (n=15:7), and (F) perigonadal fat (n=14:14) of C57BL6/J mice on high-fat high-sucrose diet (HFHSD) or chow. (G) Body weight(n=15:15), (H) glucose levels during intraperitoneal glucose tolerance test (ipGTT) (n=15:15), and (I) insulin tolerance test (ITT) (n=15:15) of HFHSD-fed C57BL6/J mice treated with 15-keto-PGE2 or vehicles. (J) Weights of perigonadal fat, inguinal fat, mesenteric fat, skeletal muscle, and brown adipose tissue (BAT) of HFHSD-fed C57BL6/J mice treated with 15-keto-PGE2 or vehicles (n=15:15). (K) Body composition of HFHSD-fed C57BL6/J mice treated with 15-keto-PGE2 or vehicles (n=15:15) (L) Phospho-Akt levels in perigonadal fat, inguinal fat, muscle, and brown adipose tissue after intraperitoneal insulin injection. (M) H&E stain of perigonadal fat and (N) average adipocyte size (n=15:15). (O) Energy expenditure measured by indirect calorimetry (n=15:15), (P) food intake (n=15:15), and (Q) physical activity (n=15:15) of HFHSD-fed C57BL6/J mice treated with 15-keto-PGE2 or vehicles. (R) Relative *Ucp1* expression in inguinal fat, perigonadal fat, and brown adipose tissue of HFHSD-fed C57BL6/J mice treated with 15-keto-PGE2 or vehicles (n=13-14 per group). (S) Body surface temperatures of interscapular area, inguinal area, and rectal temperatures of 15- keto-PGE2-treated mice and vehicle control mice after HFHSD feeding (n=15:15). (T) Body surface temperatures of interscapular area, inguinal area, and rectal temperatures of 15- keto-PGE2-treated mice and vehicle control mice during cold test (n=15:15). Data information: All data are presented as mean and standard error (S.E.M.). Statistical significance was calculated by two-sample independent *t*-test in (A, D-K, N-T) and Spearman’s correlation analyses in (B, C). \**p* < 0.05, ** *p* < 0.01, **** *p* < 0.0001. Source data are available online for this figure.

### 15-keto-PGE2 treatment protected against diet-induced obesity and improved insulin resistance without fluid retention

In view of the low 15-keto-PGE2 levels observed in obese mice, we sought to examine whether 15-keto-PGE2 could rescue obesity and insulin resistance in mice. 15-keto-PGE2 was administered to HFHSD-fed obese C57BL6/J mice for 3 weeks. Results showed that 15-keto-PGE2 protected against diet-induced obesity (Fig 2G) and markedly improved both glucose tolerance (Fig 2H) and insulin sensitivity (Fig 2I) in HFHSD-fed mice. White fat mass was also reduced in mice treated with 15-keto-PGE2 (Fig 2J) and body composition analysis showed reduced fat mass without fluid retention (Fig 2K). Immunoblots showed increased phospho-Akt in perigonadal fat, inguinal fat, brown adipose tissue, and liver of 15-keto-PGE2-treated mice (Fig 2L). Moreover, the adipocyte size was smaller (Fig 2M and N) in 15-keto-PGE2-treated mice but the number of adipocytes showed no difference (Appendix Fig S5). Higher energy expenditure (Fig 2O) was observed in mice treated with 15-keto-PGE2 compared with vehicle at the age of 8 weeks when there was no difference in body weight between the two groups. Food intake (Fig 2P) and physical activity (Fig 2Q) were similar between mice treated with 15-keto-PGE2 and vehicles. Mice receiving 15-keto- PGE2 had increased expression of *Ucp1* in inguinal, perigonadal, and brown adipose tissues (Fig 2R). Similarly, mice receiving 15-keto-PGE2 exhibited higher interscapular, inguinal, and rectal temperatures after HFHSD feeding (Fig 2S). Furthermore, mice receiving 15-keto-PGE2 exhibited higher interscapular, inguinal, and rectal temperatures than vehicle controls during prolonged cold tests (Fig 2T), indicating that 15-keto-PGE2 increases diet- and cold-induced thermogenesis.

### *Ptgr2* knockout mice with increased 15-keto-PGE2 exhibited less diet-induced weight gain and improved insulin sensitivity

As mentioned above, 15-keto-PGE2 can be irreversibly metabolized by PTGR2 into the inactive metabolite 13,14-dihydro-15-keto-PGE2, we generated *Ptgr2*^-/-^ mice, which lack the inactivating enzyme of 15-keto-PGE2 to investigate the physiological role of 15-keto-PGE2, on systemic glucose homeostasis and energy balance. As expected, relative serum 15-keto-PGE2 concentration (∼ 2.40-fold increase) and 15-keto-PGE2 content in perigonadal fat (∼1.75-fold increase) are higher in *Ptgr2^-/-^*mice than *Ptgr2*^+/+^ controls (Fig EV3A and B).

When fed with regular chow, there was no difference in body weight, fasting glucose, glucose tolerance, and insulin sensitivity between knockout mice and controls (Appendix Fig S6). However, when fed with HFHSD, *Ptgr2*^-/-^ mice exhibited significantly less body weight gain (Fig 3A) and lower fasting glucose levels compared with *Ptgr2^+/+^* littermates (Fig 3B). *Ptgr2*^-/-^ mice were more glucose-tolerant (Fig 3C) and more insulin-sensitive (Fig 3D) than controls. ^18^F-FDG-positron emission tomography (PET) showed higher ^18^F-FDG uptake following insulin stimulation in the perigonadal fat and inguinal fat of *Ptgr2*^-/-^ mice compared with controls (Fig 3E and F). Consistently, immunoblots showed significantly increased phospho-Akt levels in perigonadal fat, inguinal fat, and liver, but not in skeletal muscle of *Ptgr2*^-/-^ mice after insulin injection (Fig 3G). There was also a trend of increased phospho-Akt levels in the brown adipose tissue (BAT) of *Ptgr2*^-/-^ mice compared with *Ptgr2^+/+^* controls (Fig 3G). These data suggest that adipose tissues and liver are the primary sites of increased insulin-stimulated glucose uptake.

**Figure 3.**
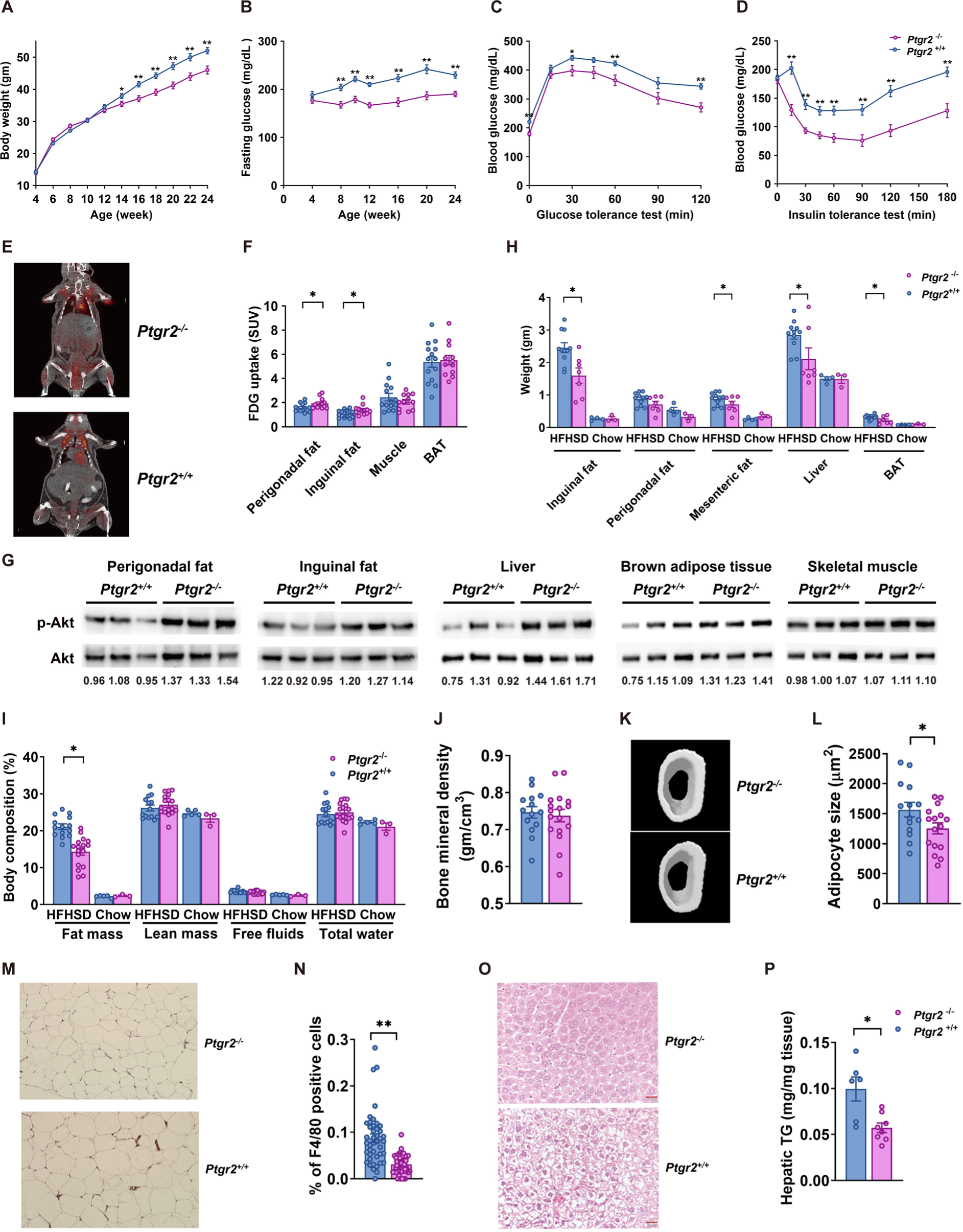
*Ptgr2* knockout mice were protected from diet-induced obesity, insulin resistance, glucose intolerance, and fatty liver without fluid retention and reduced bone density. (A) Body weight (n=20:21), (B) fasting glucose (n=20:21), and (C) glycemic level during intraperitoneal glucose tolerance test (ipGTT) (n=20:21) and (D) insulin tolerance test (ITT) (n=20:21) of *Ptgr2* ^-/-^ and *Ptgr2* ^+/+^ mice on high-fat high-sucrose diet (HFHSD). (E) ^18^F-FDG PET scan of *Ptgr2* ^-/-^ and *Ptgr2* ^+/+^ mice after injection of glucose and insulin. (F) ^18^F-FDG uptake in perigonadal fat, inguinal fat, skeletal muscle, and brown adipose tissue (BAT) after injection of glucose and insulin (n=14:12). (G) Phospho-Akt levels after intraperitoneal insulin injection in perigonadal fat, inguinal fat, skeletal muscle, and BAT. (H) Tissue weights of perigonadal fat, inguinal fat, mesenteric fat, liver, BAT (n=11:7). (I) Body composition (n=14:17), (J) bone mineral density of femur (14:17), (K) micro computed tomography (CT) image of femur of C57BL6/J mice (n=14:17), and (L) average adipocyte size. (M) F4/80 immunohistochemical stain of perigonadal fat and (N) percentage of F4/80-positive cells in perigonadal fat (n=43:56). (O) H&E stain of liver and (P) hepatic triglyceride contents of (n=6:8) of *Ptgr2* ^-/-^ mice and *Ptgr2* ^+/+^ on HFHSD. Data information: All data are presented as mean and standard error (S.E.M.). Statistical significance was calculated by two-sample independent *t*-test in (A-D, F, H, I, J, L, N, O, P). \**p* < 0.05, ** *p* < 0.01. Source data are available online for this figure.

The inguinal fat, mesenteric fat, brown fat, and liver of *Ptgr2*^-/-^ mice were smaller than those of controls under HFHSD (Fig 3H). Body composition analysis revealed significantly lower fat content in *Ptgr2*^-/-^ mice than in controls, while lean mass was not altered (Fig 3I). Importantly, the free fluid (water contained in urine), total water (water contained in urine, tissue, and blood) (Fig 3I) and the mineral density of the femur of *Ptgr2*^-/-^ mice were similar to those observed in controls (Fig 3J and K). The adipocyte size was smaller (Fig 3L), but the adipocyte number was not significantly reduced in the perigonadal fat of *Ptgr2*^-/-^ mice (Appendix Fig S7). Immunohistochemical staining showed fewer F4/80-positive infiltrating macrophages and crown-like structures in the perigonadal fat of *Ptgr2*^-/-^ mice compared with controls (Fig 3M and N). The Hematoxylin-eosin (H&E) stain of the liver showed a lesser degree of hepatic steatosis (Fig 3O) and the hepatic triglyceride content is lower in *Ptgr2*^-/-^ mice than in controls (Fig 3P). Fasting serum levels of total cholesterol, leptin, and insulin were also lower in *Ptgr2*^-/-^ mice (Appendix Fig S8).

### *Ptgr2* knockout mice exhibited increased thermogenesis

To explore the mechanism by which *Ptgr2*^-/-^ mice are leaner than *Ptgr2^+/+^* controls, we examined their energy balance. Indirect calorimetry performed at the age of 8 weeks, when there was not different between in body weight, showed significantly higher energy expenditure (Fig 4A) during the dark phase (active phase) in *Ptgr2*^-/-^ mice than in controls. There were no differences in food intake (Appendix Fig S9A), physical activity (Appendix Fig S9B), and fecal triglyceride content (Appendix Fig S9C) between *Ptgr2^-/-^* and *Ptgr2^+/+^* mice. Grossly, the inguinal fat, perigonadal fat, and brown adipose tissue of *Ptgr2*^-/-^ mice were browner in appearance (Appendix Fig S10). Chronic activation of PPARγ by ligands has been shown to induce *Ucp1* expression (Ohno et al, 2012; Kozak *et al*, 1994; Cassard-Doulcier *et al*, 1993). Immunoblots showed increased *Ucp1* protein expression in all fat depots of *Ptgr2*^-/-^ mice compared with controls (Fig 4B). RT-qPCR showed higher expression of genes involved in browning, including *Ucp1, Dio2,* and *Cidea* in perigonadal fat, inguinal fat, and brown fat of *Ptgr2*^-/-^ mice (Fig 4C-E).

**Figure 4.**
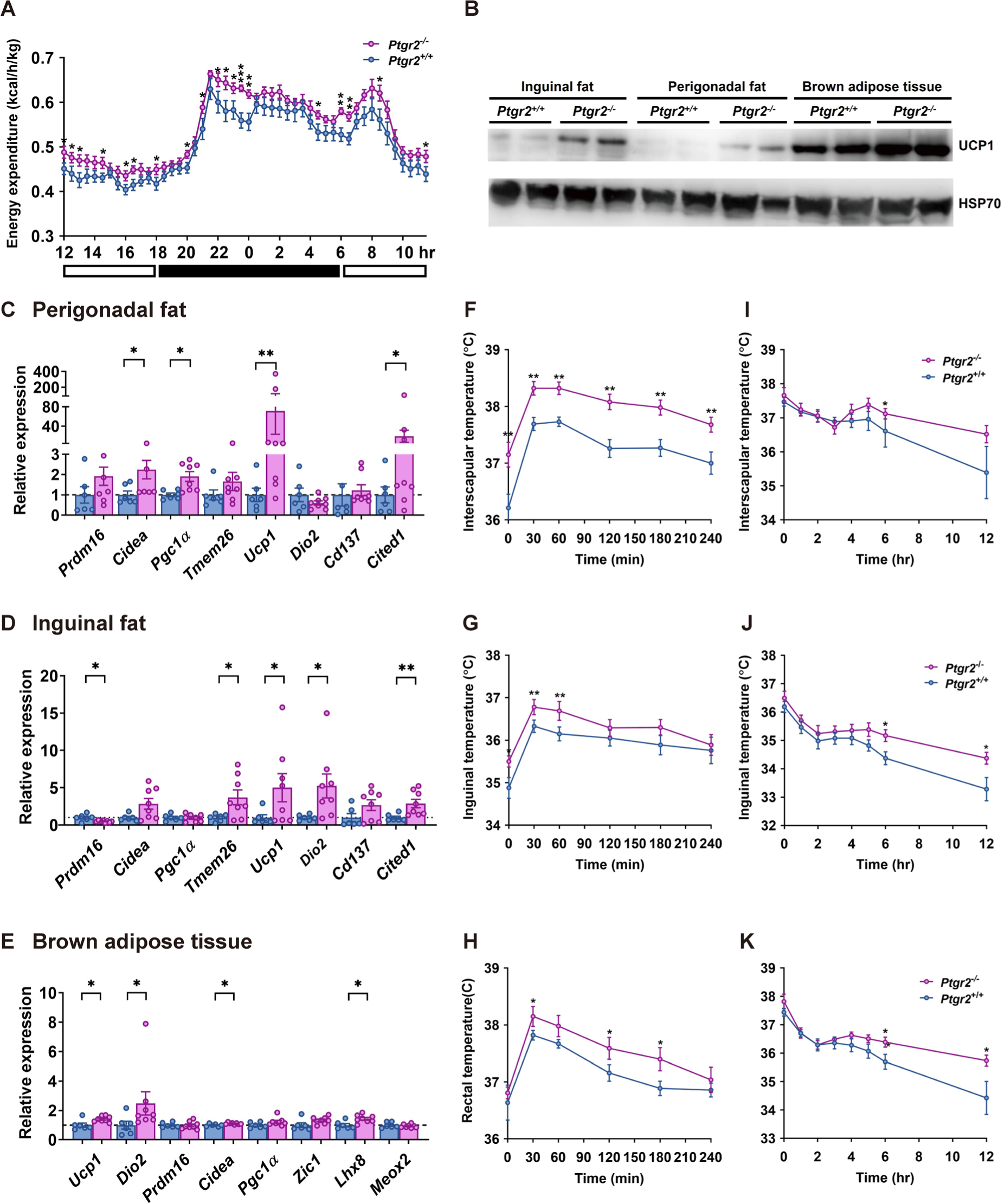
*Ptgr2* knockout mice displayed increased energy expenditure and thermogenesis. (A) Energy expenditure (the white bar indicates light-up time, the black bar indicates light-off time) (n=12:12) (B) Immunoblots showing *Ucp1* level in different fat pads. RT-qPCR of genes involved in browning of (C) perigonadal fat, (D) inguinal fat, and (E) brown adipose tissue (BAT) (n=6:8). For the diet-induced thermogenesis test, 24-week-old mice were fasted overnight. Body surface including (F) interscapular area, (G) inguinal area and (H) core rectal temperature at 0, 30, 60, 90, 120, 150, 180, and 240 min after HFHSD refeeding (n=14:16). Body surface temperatures in (I) interscapular area, (J) inguinal area, and (K) rectal temperature during cold tolerance test of *Ptgr2* ^-/-^ mice and *Ptgr2* ^+/+^ mice on HFHSD (n=14:16). Data information: All data are presented as mean and standard error (S.E.M.). Statistical significance was calculated by two-sample independent *t*-test in (A, C-K). \**p* < 0.05, ** *p* < 0.01, *** *p* < 0.001. Source data are available online for this figure.

Further evaluation of diet- and cold-induced thermogenesis revealed significantly increase in body surface temperatures of interscapular and inguinal regions, as well as in core rectal temperature after HFHSD feeding (Fig 4F-H) in *Ptgr2*^-/-^ mice compared with controls. Similarly, when exposed to a cold environment, *Ptgr2*^-/-^ mice displayed higher surface interscapular, surface inguinal, and core rectal temperatures (Fig 4I-K). These data showed enhanced thermogenesis in *Ptgr2*^-/-^ mice, probably due to the browning of fat tissues.

### A PTGR2 small-molecule inhibitor prevented diet-induced obesity and improved insulin sensitivity and glucose tolerance via activating PPARγ

Based on these findings from the *Ptgr2*^-/-^ mice, we next searched for druggable chemical inhibitors that can effectively suppress human PTGR2 (hPTGR2) enzymatic activity. Using a high- throughput screen of 12,500 chemicals, we identified 31 hits and generated 282 derivatives using a molecular hybridization strategy among the hits. The *in vitro* IC_50_ (the half-maximal inhibitory concentration), EC_50_ (the concentration for half-maximal effect), and the compound’s activity to restore 15-keto-PGE2-dependent PPARγ trans-activation in HEK293T cells expressing recombinant PTGR2 were profiled. Of the 282 newly synthesized inhibitors, a derivative of triazole-pyrimidione (BPRPT0245) (Fig 5A) exhibited an IC_50_ of 8.92 nM and an EC_50_ of 49.22 nM (Fig 5B and C). Importantly, BPRPT0245 not only increased intracellular 15-keto-PGE2 concentrations in a dose-dependent manner (Fig 5D) but also augmented insulin-stimulated glucose uptake of induced 3T3-L1 adipocytes (Fig 5E). Molecular docking showed that BPRPT0245 interfered with the interaction between 15-keto-PGE2 and NADPH within the catalytic site of hPTGR2 (Fig 5F and G).

**Figure 5.**
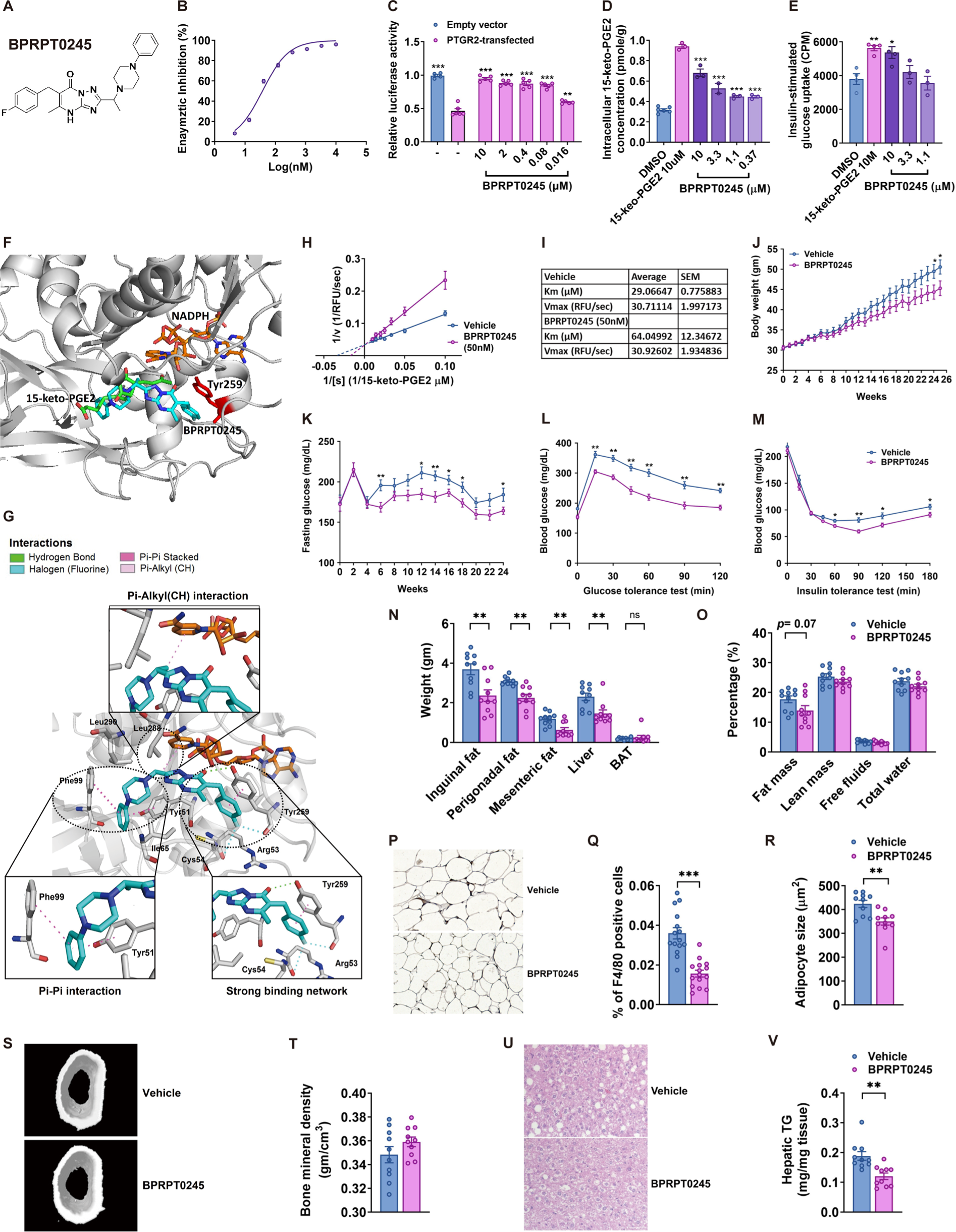
PTGR2 inhibitor BPRPT0245 protected mice from diet-induced obesity, insulin resistance, glucose intolerance, and fatty liver without fluid retention and reduced bone density. (A) Structure of BPRPT0245. (B) Half-maximal inhibitory concentration (IC_50_) of BPRPT0245. (C) Effect of BPRPT0245 on 15-keto-PGE2-dependent PPARγ transcriptional activity in PTGR2- transfected HEK293T cells (n=5 per group). (D) Intracellular 15-keto-PGE2 content of induced 3T3-L1 adipocytes treated with BPRPT0245 (n=3-6 per group) and (E) insulin-stimulated glucose uptake of induced 3T3-L1adipocytes treated with BPRPT0245 (n=3-4 per group). (F) Molecular docking of 15-keto-PGE2 (green), NADPH (orange brown), and BPRPT0245 (cyan) of PTGR2, Tyr259 (red) of PTGR2. (G) Molecular docking showing that PTGR2 inhibitor BPRPT0245 (cyan) interferes the interaction between 15-keto-PGE2 and NADPH through a strong biding network, Pi-Alkyl (CH) interaction, and Pi-Pi interaction with PTGR2 and NADPH. (H, I) Lineweaver–Burk plots showing the competitive inhibitory action of BPRPT0245 (50 nM) for the interaction between 15-keto-PGE2 and PTGR2 (n=5) values of Vmax and Km for 15-keto- PGE2 with or without the addition of BPRPT0245 (50 nM) (n=5). (I) Body weight (n=15:15), (K) fasting glucose (n=15:15), (L) glycemic level during intraperitoneal glucose tolerance test (ipGTT) (n=15:15), (M) insulin tolerance test (ITT) (n=15:15), (N) tissue weight of perigonadal fat, inguinal fat, mesenteric fat, liver, brown adipose tissue (BAT) (n=10:10), (O) body composition (n=10:10), (P) F4/80 immunohistochemical stain of gonadal fat, (Q) percentage of F4/80-positive in perigonadal fat (n=15:15), (R) adipocyte size (n=10:10), (S) representative micro computed tomography (CT) image of femur of C57BL6/J mice (n=10:10), (T) bone mineral density (n=10:10), (U) representative H&E stain of liver section, and (V) hepatic triglycerides content (n=10:10) of high-fat high-sucrose-fed C67BL6/J mice treated with BPRPR0245 (100mg/kg/day) and vehicles. Data information: All data are presented as mean and standard error (S.E.M.). Statistical significance was calculated by one-way analyses of variance (ANOVA) with Tukey’s post hoc test in (C-E) and two-sample independent *t*-test in (J-M, O, Q, R, T, V). Statistical significance was calculated by two-sample independent *t*-test in (A, C-K). \**p* < 0.05, ** *p* < 0.01, *** *p* < 0.001. Source data are available online for this figure.

Our previous crystallographic study demonstrated that NADPH binding to hPTGR2 was critical for subsequent binding of 15-keto-PGE2 (Wu *et al*, 2008). NAPDH bind mainly with Tyr259 residue of hPTGR2 through hydrogen bond (Wu *et al*, 2008), which was interfered with by BPRPT0245. BPRPT0245 forms a strong hydrogen bonding network with Tyr259, Arg53, and Cys54 of hPTGR2; Pi-Pi sandwiched with Tyr51 and Phe99 of hPTGR2; and CH-Pi interacted with the nicotinamide ring of NADPH, thus disrupting of the interaction among 15-keto-PGE2, hPTGR2, and NADPH (Fig 5F and G). Lineweaver–Burk plot further showed BPRPT0245 (50 nM) is a competitive hPTGR2 inhibitor (Fig 5H). The addition of BPRPT0245 (50 nM) changed the Km between 15-keto-PGE2 and hPTGR2 from 29.06±0.77 to 64.04±12.34 (μM) (P=0.02) but Vmax remained unchanged, consistent with a competitive inhibitor model (Fig 5I).

We further examined the effect of BPRPT0245 on obesity and insulin resistance in mice by oral administration of BPRPT0245 (100 mg/kg/day) in HFHSD-fed obese C57BL6/J mice. BPRPT0245 significantly prevented diet-induced obesity (Fig 5J), lowered fasting plasma glucose (Fig 5K), improved glucose tolerance (Fig 5L), and insulin sensitivity (Fig 5M) in mice. In addition, white fat mass and liver mass were significantly reduced by the BPRPT0245 treatment (Fig 5N). Body composition analysis revealed a marginally significantly reduced fat mass in BPRPT0245- treated mice (Fig 5O). However, BPRPT0245 did not cause fluid retention (Fig 5O) compared with vehicle. F4/80 immunohistochemical stain of the perigonadal fat showed fewer F4/80- positive macrophages in mice receiving BPRPT0245 than vehicles (Fig 5P, 5Q). The adipocyte size was reduced in mice receiving BPRPT0245 compared to vehicle (Fig 5R) but the number of adipocyte number was not changed (Appendix Fig S11). Bone mineral density analysis revealed no difference between mice receiving BPRPT0245 and vehicles for 26 weeks (Fig 5S, 5T). The extent of hepatic steatosis (Fig 5U) and hepatic triglycerides content were also significantly decreased (Fig 5V).

Pharmacokinetic studies including the half-life, clearance, steady-state volume of distribution, maximum serum concentration, and oral bioavailability after single intravenous injection and oral gavage of BPRPT0245 are shown in Appendix Table S1. There was a trend of increased 15-keto- PGE2 content in perigonadal fat (∼1.34-fold increase) and of significant increase in 15-keto- PGE2 content in inguinal fat (∼1.32-fold increase) of mice receiving BPRPT0245 compared with vehicle (Fig EV4). Pathological examination of bone marrow, brain, heart, lung, kidney, pancreas, brown fat, and perigonadal fat of mice showed no significant difference between mice receiving BPRPT0245 and vehicles at 26 weeks (Appendix Table S2). Moreover, there were no differences in serum alanine aminotransferase (ALT), total bilirubin, blood urea nitrogen (BUN), and creatinine levels (Appendix Fig S12).

## Discussion

This study, we confirmed that polyunsaturated fatty acid15-keto-PGE2 binds and activates PPARγ through covalent binding. 15-keto-PGE2 is markedly reduced in patients with type 2 diabetes or obese mice. Increasing 15-keto-PGE2 through either genetic disruption or pharmacological inhibition of its degrading enzyme PTGR2 or direct injection of 15-keto-PGE2 into mice improved insulin sensitivity and prevented diet-induced obesity. These beneficial effects were devoid of side effects of pioglitazone, including weight gain, fluid retention, and osteoporosis. Our results highlighted inhibition of PTGR2 as a new effective approach to prevent obesity, improve insulin sensitivity and insulin tolerance, and reduce hepatic steatosis without side effects of TZDs through increasing endogenous PPARγ ligands.

Consistently, a recent global mapping for lipid-interacting proteins identified PTGR2 as a lipid-interacting protein (Niphakis *et al*, 2015). In addition, using global mapping for small-molecule fragment-protein interaction, a fragment-derived ligand was identified to inhibit PTGR2 enzymatic activity. This ligand inhibited PTGR2 activity and increased 15-keto-PGE2-dependent PPARγ transcriptional activity dose-dependently *in vitro* (Parker *et al*, 2017), supporting the hypothesis that PTGR2 inhibition effectively activates PPARγ. In addition, using the Lipid-Protein Interaction Profiling (LiPIP) technique, which globally maps lipid-protein interaction and the effects of drugs on the interactions, a natural product KDT501, an extract from hops with anti- diabetic PPARγ-activating action in mice (Konda *et al*, 2014) was identified to inhibit the enzymatic activity of recombinant PTGR2 protein with an IC_50_ of 8.4 μM and the PTGR2 activity in cell lysate with an IC_50_ of 1.8 μM. Recently, a sufonyl-triazole compound HHS-0701 that interacts with the tyrosine sites of PTGR2 was also identified to inhibit PTGR2 enzymatic activity and increase intracellular 15-keto-PGE2 concentration dose-dependently (Toroitich *et al*, 2021)_._ Collectively, these data support PTGR2 inhibition as an effective way to increase 15-keto-PGE2 and to activate PPARγ.

PTGR3 is a homologous protein of PTGR2. 15-keto-PGE2 has been reported as a substrate of PTGR3 and knockdown of *Ptgr3* in adipocytes has been shown to activate PPARγ (Yu *et al*, 2013). We also found that *Ptgr3* knockout mice displayed markedly improve insulin sensitivity and glucose intolerance without side effects of TZDs (unpublished data). Taken together, these data support that increasing 15-keto-PGE2 via inhibition of its degrading enzymes as a feasible approach to activate PPARγ and to treat diabetes and obesity.

Several potential endogenous PPARγ ligands have been proposed, including oxidized polyunsaturated fatty acids, nitrated fatty acids, eicosanoids, and serotonin metabolites (Krey *et al*, 1997; Kliewer *et al*, 1997; Forman *et al*, 1995). For example, 9-HODE (9-hydroxyoctadecadienoic acid) and 13-HODE (13-hydroxyoctadecadienoic acid), the oxidized linoleic acid metabolites, were found to be abundant in atherosclerotic plaques to induce CD36 expression on macrophages to uptake lipid through activation of PPARγ (Nagy *et al*, 1998). 15-HETE (15- hydroxyeicosatetraenoic acid), a metabolite of arachidonic acid, was shown to promote adipocyte differentiation via PPARγ (Song *et al*, 2016). We Similarly, 15-deoxy-Δ12, 14-PGJ2 also induced adipocyte differentiation through PPARγ (Forman *et al*, 1995). Nitrated linoleic acid and oleic acid are potent PPARγ ligands that induce adipocyte differentiation (Schopfer *et al*, 2005). The metabolite of serotonin, 5-methyl-indole-acetate has been proposed as an endogenous PPARγ ligand that stimulates adipogenesis (Itoh *et al*, 2008).

Owing to the large binding pocket of PPARγ ligand binding domain (LBD) that can accommodate a variety of fatty acids, it is possible that PPARγ could sense and respond to different endogenous lipid ligands in response to different dietary or environmental exposure. To investigate whether the hPPARγ LBD can accommodate one or two 15-keto-PGE2 molecules, we compared the NMR spectra of the hPPARγ LBD bound to 15-keto-PGE2 at two different molar ratios (1:1 and 2:1). The similarity of the NMR spectra at both ratios indicates that the hPPARγ LBD accommodates only a single 15-keto-PGE2 molecule (Appendix Fig. S13).

However, the identification of endogenous PPARγ ligands remains difficult. Their binding affinity, binding site, specific mode of action, paracrine nature, and physiological concentrations of various proposed endogenous PPARγ ligands are yet to be defined. Previous structural analyses demonstrated that long-chain polyunsaturated fatty acid with an α, β-unsaturated moiety can form covalent bond with the Cys285 (or Cys313 in mice) residue of hPPARγ with a more stable thermal stability than others (Shiraki *et al*, 2005; Waku *et al*, 2009; Waku *et al*, 2010) and thus serve as more effective and physiologically relevant endogenous PPARγ ligands. Our study clearly confirmed that 15-keto-PGE2 which possess α, β-unsaturated moiety can form covalent binding with the Cys285 residue of hPPARγ LBD (or Cys313 residue of mPPARγ). Given the relatively low concentrations of various proposed endogenous PPARγ ligands, the high affinity of 15-keto- PGE2 to PPARγ through covalent binding suggests that 15-keto-PGE2 might be a physiologically functional PPARγ ligand.

The mechanism underlying the differential effects of 15-keto-PGE2 compared to other PPARγ modulators, such as non-covalent modulators (e.g. full agonists TZD and partial agonists MRL24 and SR1664) and covalent modulators (e.g. the transcriptionally neutral antagonist GW9662 and the transcriptionally repressive inverse agonist T0070907) remains to be elucidated.

The hPPARγ ligand-binding domain (LBD) consists of 13 α-helices (helices 1-12, and 2’), four small β-sheets (β1-4) (Nolte et al., 1998; Zoete et al., 2007), and link loops. The full hPPARγ agonists TZDs non-covalently bind to the activation function-2 (AF-2) surface comprising helix 3, helix 4/5, and helix 11 (the orthosteric binding pocket) and stabilizes the helix 12 mainly through hydrogen bond to Tyr473. This classic helix 12-dependent agonism of hPPARγ has been proposed to be associated with the side effects of TZDs (Bruning et al., 2007; Capelli et al., 2016; Hughes et al., 2012; Hughes et al., 2014; Miyamae. et al., 2021; Thangavel et al., 2017). In contrast, the synthetic partial agonists such as MRL24 and SR1664 non-covalently bind to an alternative binding pocket surrounded by helix 3, helix 5, β-sheets, helix 2-2’ link of m, and Ω loop. Despite their low PPARγ transactivation activity, these partial agonists lower insulin resistance *in vivo* without side effects of TZD, possibly by interfering CDK5 (cyclin-dependent kinase 5)-mediated Ser245 phosphorylation at the helix 2-2’ link of hPPARγ, recruiting different co-activators and co-repressors (Hughes, et al., 2014; Dias et al., 2020), or indirectly stabilizing helix 12 (Bruning et al., 2007; Choi et al., 2010; Choi et al., 2011; Chrisman, et al., 2018; Hughes, et al., 2014).

X-ray crystallography revealed that the transcriptionally neutral covalent PPARγ antagonist GW9662 forms a covalent bond with Cys285 (helix 3) and hydrogen bonds with Tyr327 (helix 5), His449 (helix 10), and Tyr473 (helix 12) of hPPARγ LBD. These interactions effectively block the binding of classical full agonists, such as rosiglitazone, to the orthosteric pocket (Appendix Fig S14A, B). However, GW9662 does not inhibit the binding of partial agonists, such as MRL20, to the alternative binding site (Hughes et al., 2014).

In contrast, the transcriptionally repressive covalent inverse agonist T0070907, which differs from GW9662 by only a single nitrogen atom, exhibits a similar X-ray co-crystal structure with GW9662 binding to the hPPARγ LBD (Appendix Fig S14C, D) (Hughes et al., 2014). The polar pyridyl group of T0070907 interacts with a water molecule, forming a hydrogen bond network that connects the Arg288 and Glu295 residues in helix 3, thereby altering the recruitment profiles of co-activators, such as TRAP220 (thyroid hormone receptor-associated protein 220), and co-repressors, such as NCoR1 (nuclear receptor co-repressor 1) (Brust et al., 2018). The recruitment of co-repressors like NCoR1 by T0070907 results in a distinct transcriptionally repressive hPPARγ LBD conformation. This conformation is stabilized by several polar π-stacking interactions among the pyridyl group of T0070907 and residues H323 in helix 5, H449 in helix 11, and Y477 in helix 12. Additionally, a network of hydrogen bonds and electrostatic interactions, including Gln286 in helix 3, Tyr327 in helix 5, and Met364 and Lys367 in helix 6, further supports this repressive conformation. These conformational changes flip the helix 12 into a pocket flanked by helix 3, helix 2’, and β-sheets, effectively blocking the binding of TZDs and repress PPARγ transcriptional activity (Appendix Fig S14C, D) (Chrisman et al., 2018; Shang et al., 2020; Irwin et al., 2022).

We confirmed that 15-keto-PGE2 forms a covalent bond with Cys285 at helix 3, similar to the bind mode of GW9662 and T0070907. In addition, our molecular docking showed that 15- keto-PGE2 forms a covalent bond with human Cys285 (helix 3) and hydrogen bonds with Tyr327 (helix 5), Arg288 (helix 3), and Ile326 (helix 5) with hPPARγ LBD. These binding restrains 15- keto-PGE2 within a binding pocket between helix 3, helix 5, and the β-sheets, which is distant from helix 12 (Appendix Fig S14 E, F).

Furthermore, our 2D ^1^H-^15^N-TROSY-HSQC NMR spectra that compare apo- and 15-keto- PGE2-bound hPPARγ LBDs showed missing peaks at helix 1 (His217), helix 2-2’ link (Ser245), β2-β4 (Gly344, Gly346), helix 7 (Phe368, Ala376) and helix 8-9 (Ser394), and chemical shift peaks at helix 1(Ser221, Ser225), helix 2 (Arg234), helix 2’ (Met252), helix 3 (Ala235), helix 3 (Ile303), helix 7 (Glu378, Asn375, Lys373), β1 (Val248), and β2-β4 (Gln345). Most of signal changes are close to the alternative binding site and distal to helix 12 and AF-2 surface (Figure EV5). Previous studies showed that the NMR spectra of GW9662-bound hPPARγ LBD had little difference to the spectra of apo-hPPARγ LBD (Brust et al., 2018; Ardenkjær-Skinnerup et al., 2024), and the T0070907-bound NMR peaks covered extensive residues located in the β-sheets, helix 3, helix 7, and a peak of Val322 on helix 5 within the AF-2 surface (Brust, et al.,2018).

Collectively, GW9662 blocks the binding of full agonists TZDs by interfering with their interaction with the AF-2 domain but not the alternative binding site. In contrast, T0070907 forms an extensive interaction network via the polar pyridyl ring and causes a helical turn of helix 12 that prevent TZD binding and recruit increased transcriptional co-repressors, thereby suppressing hPPARγ transcriptional activity. 15-keto-PGE2 covalently interacts with the helix 1, 2, 2-2’ link, β sheets, helix 3, 5, and 7-9, close to the alternative binding site and distal to helix 12.

Furthermore, posttranslational modifications, such as phosphorylation and acetylation, were reported to regulate PPARγ functions. CKD5-mediated Ser273 phosphorylation inhibited mPPARγ activation and was interfered by rosiglitazone (Choi et al., 2010, Mottin et al., 2015, Montanari et al., 2020). Inhibition of HDAC3 (histone deacetylase 3)-mediated deacetylations induced the transcriptional activity of PPARγ, and pioglitazone treatment increased the acetylation of PPARγ (Jiang et al., 2014; Ding et al., 2020). Here, we found that 15-keto-PGE2 dose-dependently reduced phosphorylation of Ser273 at the 2-2’ link in both cultured murine adipocytes and adipose tissue, which is close to the alternative binding site (Appendix Fig S15A, B), and there was no altered acetylation of the PPARγ LBD by 15-keto-PGE2 (Appendix Fig S16).

The major limitation of our study is the uncertainty of physiological concentration of 15-keto- PGE2 due to different detection techniques. A systemic screen of 94 lipids confirmed 15-keto- PGE2 as an endogenous PPARγ ligand *in vivo* with a physiological concentration approximating sub-micromolar range (∼0.25 μM) (Harmon *et al*, 2010; Harmon *et al*, 2011). Another chromatography study reported a tissue concentration of 15-keto-PGE2 at ∼4.3 μM (1500 pg/mg) in human colon (Chhonker *et al*, 2021). However, another systemic chromatographic survey of 15-keto-PGE2 in different human tissues reported only ∼0.04 μM in murine small intestines (13.81 pg/mg) (Yamada *et al*, 2015). In addition, it is also difficult to estimate the intracellular or spatial distribution of 15-keto-PGE2 due to the paracrine nature of eicosanoids.

Although we used relative high concentration (1-20 μM) of 15-keto-PGE2 in cell experiments, the maximal dose of added 15-keto-PGE2 (10-20 μM) only raised intracellular 15-keto-PGE2 by ∼1.25 to 1.78 folds, indicating that near-physiological concentrations can activate PPARγ (Fig EV1). Furthermore, either genetic inhibition or chemical inhibition increased 15-keto-PGE2 tissue concentrations by only ∼1.7 and ∼1.3 folds, respectively (Fig EV3 and EV4). Such modest increases in 15-keto-PGE2 content can clearly leads to obvious beneficial metabolic effects. Given the high affinity of 15-keto-PGE2 to PPARγ through covalent binding, it is likely that 15-keto- PGE2 is a physiologically functional PPARγ ligand even at a relatively low concentration. Lastly, we cannot rule out the possibility that metabolites involved in the metabolic flux of 15-keto-PGE2 influence the phenotype of *Ptgr2*-deficient mice. Therefore, we measured the levels of PGE2, PGF2α, PGD2, 15-deoxy-PGJ2, and 6-keto-PGF1α in the perigonadal fat and serum of *Ptgr2*^+/+^ and *Ptgr2*^−/−^ mice (Inazumi T et al., 2020; Forman et al., 1995) No significant differences were observed between the *Ptgr2^+/+^* and *Ptgr2^−/−^* mice, suggesting that their phenotypic differences are primarily mediated by 15-keto-PGE2 (Appendix Fig S17).

In conclusion, this study showed that genetic or pharmacological inhibition of PTGR2 prevented diet-induced obesity, reduced insulin resistance, and decreased hepatic steatosis via increasing endogenous PPARγ ligands without the side effects of synthetic ligands TZD.

## Materials and Methods

### Generation of *Ptgr2* knockout mice

The generation of *Ptgr2* knockout mice has been described previously (Chen *et al*, 2018). Briefly, the mouse *Ptgr2* gene comprises 10 exons with exon 3 containing the catalytic domain. Deletion of exon 3 is predicted to delete the catalytic domain and creates a frameshift mutation resulting in a stop codon in exon 4. The construct used for targeting the *Ptgr2* gene was designed to insert a loxP sequence together with an “FRT-flanked” pgk-neo cassette in intron 2 and a loxP sequence in intron 4. The liberalized targeting vector was electroporated into the 129/J embryonic stem cell line and selected by neomycin and ganciclovir. The selected clone was used for blastocyst injection. Immunoblots for Ptgr2 showed deletion of Ptgr2 in all tissues of knockout mice compared with wild-type controls (Appendix Fig S18).

### Animal models

All animal experiments were performed according to institutional ethical guidelines and were approved by the Institutional Animal Care and Use Committee (IACUC) of National Taiwan University Medical College (IACUC No.: 20140456). All mice were housed under standard conditions at 23°C and 12/12 hrs. light/dark (7 AM-7 PM) cycle in animal centers of the National Taiwan University Medical College, which is accredited by the Association for Assessment and Accreditation of Laboratory Animal Care International (AAALAC). Mice were provided shelters and sticks weekly. We used adequate anesthetic procedure to reduce pain, suffering and distress according to our animal center regulation. We reported expected or unexpected adverse events to the veterinarian of our animal center. The humane endpoints include 20% weight loss, Rough hair coat, hunched posture, lethargy or persistent, or any condition interfering with eating or drinking. Mice were fed on either a high-fat high-sucrose diet (HFHSD) (cat. no. D12331, Research Diets) or a regular chow diet (cat no. 5001, Lab Diet). BPRPT0245 was dissolved 3% dimethylacetamide and 10% cremophor in water and administered daily to HFHSD-fed obese C57BL6/J mice by oral gavage (100 mg/kg/day). Mice were pursued from the National Laboratory Animal Center, Taiwan.

### Cell culture

All cells were cultured at 37°C, 5% CO_2_ in a humidified incubator. HEK293T cells were maintained in DMEM (cat. no. SH30003.02, HyClone) supplemented with 10% fetal bovine serum (FBS) (cat. no. 04-001-1A, Biological Industries) and 1% antibiotic/antimycotic solution (cat. no. SV30079.01, HyClone). 3T3-L1 preadipocytes were maintained at ∼70% confluence in DMEM with 10% calf serum (cat. no. 16170078, Gibco) and 1% antibiotic/antimycotic solution. For adipocyte differentiation, confluent 3T3-L1 cells (defined as day 0) were exposed to an induction medium containing 10% FBS, 1 μM dexamethasone (cat. no. D4902, Sigma-Aldrich), and 1 μg/ml insulin (Humulin R, Eli Lilly) in DMEM with or without 0.5 mM isobutyl-methylxanthine (cat. no. sc-201188A, Santa Cruz Biotechnology) as indicated. After two days, the medium was replaced with DMEM containing 10% FBS and 1 μg/ml insulin and was replenished every two days until assay.

### Gal4-PPARγ/UAS-LUC reporter assay

Gal4-PPARγ/UAS-LUC reporter assay was conducted as previously described (Wu *et al*, 2008) with minor modifications. Briefly, HEK293T cells were seeded in 24-well plates at 1×10^5^ cells/well. After 24-hr growth, a DNA solution containing UASG reporter construct, GAL4-PPAR expression plasmid, and TK-Rluc (Renilla luciferase) reporter construct (internal control) was transfected using TurboFect™ transfection reagent (cat. no. R0532, Thermo Fisher). Cells were treated and harvested after another 24 and 48 hrs., respectively. Luciferase activity was measured using Luc-Pair™ Duo-Luciferase HS Assay Kit (cat. no. LF600, Promega) and normalized with the TK reporter signal.

### RNA extraction, cDNA synthesis, and real-time quantitative PCR (RT-qPCR)

A total of 1×10^5^ 3T3-L1 preadipocytes were seeded in 6-well plates and differentiated for six days. The 3T3-L1 adipocytes were then harvested in 1 ml of REzol C&T (cat. no. PT-KP200CT, Protech), and total RNA was extracted according to the manufacturer’s instructions with slight modifications. Briefly, 200 μl of chloroform were added to 1 ml of sample in REzol, and samples were vigorously mixed by shaking for 30 sec, followed by incubation at room temperature for 5 min. Then, samples were centrifuged at 12,000 g for 15 min at 4°C, and 400 μl of the upper aqueous phase were transferred to a new 1.5-ml tube. An equal volume of isopropanol was added. Samples were inverted several times for mixing and then centrifuged at 12,000 g for 10 min at 4°C. The RNA precipitate, which formed a pellet at the bottom of the 1.5-ml tube, was washed three times with 75% ethanol and air-dried for 15 min. Pellets were dissolved in RNase-free water. The concentration of RNA was measured using Nanodrop. cDNA was synthesized using a reverse transcription kit (cat. no. K1622, Thermo Scientific) using the oligo(dT)18 primers. RT-qPCR was performed in a 10-μl reaction with 50 ng cDNA and 0.2 μM primer using SYBR green reagent (cat. no. 11203ES08, YEASEN). Mouse peptidylprolyl isomerase A (Ppia) mRNA was used as the internal control. RT-qPCR reactions were performed using ABI 7900HT FAST (Applied Biosystems) and Sequence Detection Systems (SDS v2.3, Applied Biosystems). All qPCR reactions were run in duplicates. The primers used are listed in Appendix Table S3.

### Plasmid construction and site-directed mutagenesis

Murine PPARγ plasmid (pCMV6-Pparg) was purchased from OriGene (cat. no. MC201042, OriGene, USA). A cysteine-313 to alanine (C313A) substitution of mPPARγ construct was generated with a mutagenesis kit (cat. no. 210518, Agilent, USA) following the manufacturer’s protocol. Primers for site-directed mutagenesis was designed using a web-based program (http://www.genomics.agilent.com/primerDesignProgram.jsp) (Appendix Table S4).

### Preparation of cysteine-coupled 15-keto-PGE2 proteins

Five mg of ovomucoid (OVO, cat. no. T9253, Sigma-Aldrich) and bovine serum albumin (BSA, cat. no. A2153, Sigma-Aldrich) were dissolved in 1 mL of PBS containing 5% 2-mercaptoethanol (2-ME) (M6250, Sigma-Aldrich) and gently agitated at room temperature for 1 hr., respectively. The reduced proteins were buffer-exchanged with PBS using Amicon Ultra-15 Centrifugal Filter Units (cat. no. UFC901024, Millipore) to remove 2-ME. The proteins were then incubated with a 10-fold molar excess of 15-keto-PGE2 at room temperature for 1 hr. with gentle agitation. The cysteine-coupled proteins, 15-keto-PGE2-cysteine-OVO 15-keto-PGE2-cysteine-BSA were further exchanged with PBS buffer to remove free 15-keto-PGE2 and stored at 1 mg/mL

### Generation of monoclonal antibodies against cysteine-coupled 15-keto-PGE2

Ten Balb/c mice (National Laboratory Animal Center, Taiwan) were immunized with 15-keto- PGE2-cysteine-OVO (100 μg/mouse) emulsified with complete Freund’s adjuvant (cat. no. F5881, Sigma-Aldrich) via subcutaneous injection. After 4 weeks, mice were subcutaneously injected with 15-keto-PGE2-cysteine-OVO (100 μg/mouse) and emulsified with incomplete Freund’s adjuvants (cat. no. F5506, Sigma-Aldrich) three times at a two-week interval. Mice received 10 μg of 15-keto-PGE2-cysteine-OVO via tail vein three days before spleen harvest. Hybridomas were prepared by fusing splenic cells with the mouse myeloma cell line FO (cat. no. CRL-1646, ATCC) using PEG 1500 (cat. no. 10783641001, Roche). Hybridomas were selected in complete Dulbecco’s modified Eagle’s medium (DMEM) containing hypoxanthine-aminopterin-thymidine (cat. no. 11067030, Thermo Fisher) and UltraCruz Hybridoma Cloning Supplement (cat. no. sc-224479, Santa Cruz) for 12-14 days. Cell culture supernatants were screened with ELISA using 15-keto-PGE2-cysteine-BSA as antigens. Positive hybridomas were then cloned by limiting dilution. Antibody isotypes were determined with a Rapid ELISA Mouse mAb Isotyping Kit (cat. no. 37503, Thermo Fisher). Monoclonal antibodies were purified from hybridoma culture supernatants using Protein A Sepharose CL-4B (cat. no. 17078001, GE). The purification procedures were performed according to the manufacturer’s manual. Four anti-cysteine-coupled 15-keto-PGE2 hybridoma clones were prepared. Clones 1A6 and 17E7 had superior specificity and reacted only to cysteine-coupled 15-keto-PGE2 (Appendix Fig S19).

### Immunoprecipitation and proteomic analysis

HEK293T cells were transfected with murin PPARγ (mPPARγ) and mPPARγ C313A plasmids for immunoprecipitation, which are Myc-DDK-tagged (OriGene, USA). After 24 hrs, cells were treated with dimethyl sulfoxide (DMSO) or 30 μM 15-keto-PGE2 for an additional 24 hrs. and harvested with cell lysis buffer (50 mM Tris-HCl, pH 7.4, 1 mM EDTA, 150 mM NaCl, and 1% Triton X-100). According to the manufacturer’s protocol, proteins were immunoprecipitated using Anti-FLAG^®^ M2 Magnetic Beads (cat. no. F1804, Sigma-Aldrich). Subsequently, the eluted proteins were separated by SDS-PAGE and subjected to immunoblot analysis. Following separation by SDS-PAGE, proteins were stained with Coomassie blue and SYPRO-Ruby stain. The protein bands of interest were cut out for in-gel tryptic digestion followed by C18 Zip-Tip clean-up (Millipore, USA). For proteomic shotgun identifications, nanoLC-nanoESI-MS/MS analysis was performed on a nanoAcquity system (Waters, USA) connected to the LTQ Orbitrap Velos hybrid MS (Thermo Electron, USA) equipped with a PicoView nanospray interface (New Objective, USA). Samples were loaded onto a C18 BEH column (75 μm × 25 cm length, 130 Å, 1.7 μm particle size) (Waters, USA) and separated by a segmented gradient form in 60 min from 5% to 40% acetonitrile (with 0.1% formic acid) at a constant flow rate of 300 nL/min with a column temperature of 35°C. The mass spectrometer was operated in the data-dependent mode. Full scan MS spectra were acquired in the Orbitrap at a resolution of 60,000 (at m/z 400) with an automatic gain control target of 5’105. The 10 most abundant ions were isolated for high-energy collision dissociation, MS/MS fragmentation, and detection in the Orbitrap. For MS/MS measurements, a resolution of 7500, an isolation window of 2 m/z, and a target value of 50,000 ions, with maximum accumulation times of 100 ms, were used. Fragmentation was performed at 35% normalized collision energy and an activation time of 0.1 ms. Ions with single and unrecognized charge states were excluded. Protein identification and modification were analyzed using Mascot Daemon (Matrix Science Inc., Chicago).

### PPRE report assay

For PPRE reporter assay, HEK293T cells were seeded in 24-well plates at 1×10^5^ cells/well. After 24-hr growth, PPRE-LUC reporter vector, TK-LUC reporter and wild-type or mPPARγ C313A were transfected into cells was transfected via TurboFect™ transfection reagent (cat. no. R0532, Thermo Fisher). After additional 24 and 48 hrs, cells were treated and harvested. The luciferase activity was measured by Luc-Pair™ Duo-Luciferase HS Assay Kit (cat. no. LF600, Promega, USA) and normalized to the TK reporter signal.

### Analysis of ligand binding to PPARγ with native mass spectrometry

Wide-type mPPARγ ligand binding domain (LBD), mPPARγ LBD C313A mutant and mPPARγ LBD H351A mutant were mixed together in a 1:1:1 molar ratio and then incubated with 15-keto- PEG2 or pioglitazone in a 1:10 molar ratio for 5 min at room temperature, respectively. The protein mixture was buffer-exchanged into 500 mM ammonium acetate buffer pH 7.5 using 7K Zeba™ Spin Desalting Column (cat. no, 89877, Thermo) and immediately introduced into a modified Q-Exactive mass spectrometer (Thermo). The sets of optimized parameters were applied to analyze the ligand binding, including the higher-energy collisional dissociation (HCD) energy of 20V; spray voltage, 1.1 kV; capillary temperature, 200°C; S-lens RF level, 200; and resolution, 12500 at m/z 400. Spectra were acquired and processed manually using Thermo Xcalibur Qual Browser (version 4.4.16.14). The relative percentage of compound-bound forms was quantified by the UniDec software and the degree of effector coupling was calculated by normalizing the relative percentage of bound forms to the sum of the percentage of unbound and bound forms.

### Generation of PPARγ-null 3T3-L1 stable cell lines using CRISPR/Cas9 System

Gene editing was performed in 3T3-L1 preadipocytes using the clustered regularly interspaced short palindromic repeats (CRISPR)/Cas 9 system. Two single-guide RNAs (sgRNA) (Appendix Table S5) targeting exon 6 of mouse *Pparγ* were designed using a web-based design tool (http://crispr.mit.edu/). These oligonucleotides were cloned into the pSurrogate reporter and pAll- Cas9.Ppuro vector obtained from Academia Sinica, Taiwan. The pSurrogate reporter contains an EGFP and an out-of-frame mCherry downstream of the sgRNA; the pAll-Cas9.Ppuro vector expresses Cas9 nuclease and sgRNA. Once a double-strand breaking at the target site in pSurrogate reporter plasmid was created by Cas9, insertions and deletions (indels) could be introduced at the cleaved site. The indels cause frameshifts and result in the expression of the mCherry gene. 3T3- L1 preadipocytes were co-transfected with pSurrogate reporter and pAll-Cas9.Ppuro vectors at 80- 90% confluency using PolyJet™ transfection reagent (cat. no. SL100688, SignaGen Laboratories, USA). Two days after transfection, EGFP/mCherry double-positive cells were enriched by fluorescent-activated cell sorting. Single cells were isolated and expanded. The genomic DNA was amplified by PCR and sequenced to confirm the knockout cell lines. The primer sequences are shown in Appendix Table S5.

### Lentivirus production and transduction

The mPPARγ C313A DNA sequence was amplified from pCMV6-Pparg C313A and subcloned into the pLVX-IRES-Neo vector with XhoI/NotI restriction sites. The lentiviral expression plasmids, pMD.G, and pCMVΔR8.91 (RNAi Core, Academia Sinica, Taiwan), were co-transfected into HEK293T cells. The medium containing lentivirus was collected at 40 and 64 hrs. after transfection, centrifuged at 1,200 rpm for 5 min, and then filtered through a 0.45-μm filter. mPPARγ-null 3T3-L1 preadipocytes were infected with the lentivirus in the presence of 10 μg/ml polybrene (cat. no. sc-134220, Santa Cruz). Then, 48 hrs. after infection, cells selected with 400 μg/ml G418. The expression of PPARγ was verified by immunoblotting using an anti-PPARγ antibody (cat. no. sc-7273, Santa Cruz). HSP70 was probed with an anti-HSP70 antibody (cat. no. ab45133, Abcam) as a loading control.

### Insulin-stimulated glucose uptake assay by (^3^H)2-deoxyglucose

Differentiated 3T3-L1 cells were starved with serum-free DMEM containing 0.2% BSA (cat. no. A9647, Sigma-Aldrich). Cells were rinsed twice with KRH buffer (137 mM NaCl, 4.7 mM KCl, 1.85 mM CaCl_2_, 1.3 mM MgSO_4_, 50 mM HEPES and 0.1% BSA, pH 7.4), and treated with or without 1 μg/mg insulin in the presence of 20 μM cytochalasin B (cat. no.11328, Cayman Chemical), a GLUT inhibitor for measuring non-specific glucose uptake, for 30 min. Cells were then incubated with 0.5 μCi [^3^H]-2-deoxy-D-Glucose (cat. no. NET328A001MC, PerkinElmer) and 0.1 μM 2-deoxy-D-Glucose (2DG) (cat. no. 14325, Cayman Chemical) for 5 min. Uptake of 2DG was terminated by rapidly removing medium and washing with ice-cold PBS three times. Cells were lysed with 0.1% SDS, and the radioactivity was measured using a liquid scintillation counter. Insulin-mediated glucose uptake was calculated by subtracting insulin-treated glucose uptake with basal glucose uptake.

### Human subjects for measurement of serum 15-keto-PGE2 levels

Fifty non-diabetic male participants were recruited from a community-based screening for diabetes mellitus in Yunlin County in Taiwan. Twenty-four male patients with type 2 diabetes were recruited from the metabolic clinic of the Yunlin branch of National Taiwan University Hospital. Another 24 age- and body mass index (BMI)-matched male non-diabetic patients were recruited from a community-based screening program. The Institutional Review Board approved the study protocol of the National Taiwan University Hospital (serial number: 9561706032). Written informed consent was obtained from every participating subject. All procedures performed in this study involving human participants followed the WMA Declaration of Helsinki and the Department of Health and Human Services Belmont Report.

### 15-keto-PGE2 extraction from tissues, serum, and cultured cells

Liquid nitrogen-frozen tissue was grinded using pestle and mortar. Approximately 500 mg of tissue powders were immersed with 400 μL of the upper layer of acetonitrile (ACN) /hexane mixture supplemented with 0.5% formic acid, and then homogenized with ceramic beads by MagNA Lyser (Roche, USA). The homogenate was then added with 600 μL ACN containing 13,14 dihydro-15-keto-PGE2-d4 (cat no. 10007978, Cayman Chemical, Ann Arbor, MI) in 1:20,000 dilution and homogenized, followed by the addition of 600 μL ddH_2_O. The mixture was centrifuged at 12,000 g for 10 min at 0°C. The lower layer was transferred to a new tube. A C-18 solid-phase extraction (SPE) cartridge (Cat no. 400020, Cayman Chemical) was sequentially activated with 20 mL methanol and 20 mL ddH_2_O. The protein sample was loaded into activated SPE cartridge. Once the sample passed the cartridge, the cartridge was washed with 5 mL 15% methanol and then 5 mL ddH_2_O. Finally, the sample was eluted with 10 mL HPLC-degree methanol (Methanol Chromasolv LC-MS, Fluka) and aliquoted into new 2-mL tubes. The elute was air-dried using the SpeedVac system (Savant SPD1010, Thermo Scientific) and stored at - 80°C. The samples were reconstituted with 100 μL methanol before performing LC-MS/MS analysis.

600 μL of serum pooled from two individual mice with the same genotype was mixed with 1800 μL of ACN and 0.25% formic acid. The mixture was centrifuged at 12,000 g for 10 min at 0°C. The supernant was transferred into a new eppendrof and added 600 μL of ACN. The mixture was centrifuged at 12,000 g for 10 min at 0°C again. The supernant was air-dried using the SpeedVac system (Savant SPD1010, Thermo Scientific) and stored at −80°C. The samples were reconstituted with 60 μL methanol before performing LC-MS/MS analysis.

For 15-keto-PGE2 extraction from cultured cells (in 6-well culture plates), the medium was removed, and cells were scraped by adding 500 μL methanol containing 13,14 dihydro-15-keto- PGE2-d4 (1:20000 dilution), followed by the addition of 1 mL PBS buffer. The lysate was centrifuged at 300 g for 5 min to remove cellular debris. The supernatant was transferred to a new tube. The C18 SPE cartridge was sequentially activated with 2 mL methanol and 2 mL ddH_2_O. The lower-layer sample was transferred to the C18 SPE cartridge. Once the lysate passed the cartridge, the SPE cartridge was washed with 2 mL 15% methanol and 5 mL ddH_2_O. The sample was then eluted with 5 mL HPLC-degree methanol and aliquoted into new 2-mL tubes. The elute was air-dried using a SpeedVac system and stored at −80°C. The dried samples were reconstituted with 60 μL methanol before analysis.

### Glucose and insulin tolerance test

Glucose tolerance was evaluated by the oral (OGTT) and intraperitoneal glucose tolerance test (ipGTT) after a 6-hr fast. For the OGTT, glucose water (1 mg/g) was given by oral gavage, and tail blood glucose was measured with a glucometer (ACCU-CHEK Performa, Roche) at 0, 15, 30, 45, 60, 90, and 120 min. For the ipGTT, tail blood glucose was measured at 0, 15, 30, 45, 60, 90, and 120 min after intraperitoneal injection of glucose water (1 mg/g). For the insulin tolerance test (ITT), mice were fasted for 4 hrs. and then injected intraperitoneally with 1 U/kg of insulin (Humulin R, Eli Lilly). Tail blood glucose was measured at 0, 15, 30, 45, 60, 90, 120, and 180 min.

### Measurement of insulin signaling

To evaluate insulin signaling *in vivo*, mice were fasted overnight. Tissue was harvested 15 min after intraperitoneal insulin injection. The samples (perigonadal fat, inguinal fat, brown adipose tissue and liver) were extracted with RIPA buffer (50 mM Tris-HCl, pH 7.4, 2 mM ethylenediaminetetraacetic acid [EDTA]), 150 mM NaCl, 50 mM NaF, 1% Nonidet P-40, 1 mM phenylmethylsulfonyl fluoride [PMSF], 0.5% sodium deoxycholate, and 0.1% sodium dodecyl sulfate [SDS]) containing phosphatase inhibitor cocktail (cat. no. 04693132001, Roche) and homogenized. The homogenates were then centrifuged at 13000 rpm for 10 min at 4°C to remove debris. Samples were separated by SDS-polyacrylamide gel electrophoresis, transferred to polyvinylidene difluoride (PVDF) membrane, blocked with 3% BSA, and probed with anti-phospho-Akt antibody (cat. no. 4058, Cell Signaling) and anti-Akt antibody (cat. no. 9272, Cell Signaling), and then with HRP conjugated anti-rabbit IgG antibody (1:10000; cat. no. GTX26721, GeneTex).

### Hematoxylin-eosin (H&E) stain

Tissues were fixed in 4% paraformaldehyde and then embedded in paraffin before sectioning and staining. The stained sections were scanned and analyzed using a MIRAX viewer (http://www.zeiss.de/mirax) and Image J software (http://rsbwed.nih.gov/ij/).

### Energy expenditure, food intake and physical activity

Metabolic measurements (food and water intake, locomotor activity, VO_2_ consumption and VCO_2_ production) were obtained using the Promethion metabolic phenotyping system (Sable Systems). The mice were housed in the Promethion system with ad libitum access to food and water. Monitoring was performed for 5 days after mice have been acclimatized to the cages for 2 days.

### Cold-induced and diet-induced thermogenesis

For the cold tolerance test, 24-week-old mice with matched average body weight from the two groups were placed individually on HFHSD in a 4°C chamber. The rectal temperature of the mice was measured after 0, 1, 2, 3, 4, 5, 6, 12, and 18 hrs. For measuring diet-induced thermogenesis, 24-week-old mice were fasted overnight for 18 hrs. Then, we measured their rectal temperature at 0, 30, 60, 90, 120, 150, 180, and 240 min after HFHSD refeeding. The rectal and surface temperature were detected by thermal probes (Physitemp Therma TH-5, ThermoWorks, Alpine and MicroTherma 2T, ThermoWorks, Alpine, respectively).

### Positron emission tomography (PET) / CT (computer tomography) for assessment of insulin-stimulated glucose uptake

The measurement of glucose uptake in different tissues followed previous protocols (Cheng *et al*, 2011; Momcilovic *et al*, 2018). Briefly, mice were fasted for 4 hrs. A total of 0.5 MBq [^18^F]-FDG in a 0.1-ml volume was administered through the tail vein, followed immediately by injecting intraperitoneally 1.6 U/kg of insulin. A whole-body scan was performed for a total of three cycles (10 min per cycle) using a PET/CT scanner (eXplore Vista DR, GE). Images were analyzed using the Amide software (Loening & Gambhir, 2003). Region of interest (ROI) was determined using manual sagittal, horizontal, and vertical slices. Standardized uptake values of each ROI (for each tissue) were calculated to estimate glucose uptake.

### Immunohistochemistry staining

Mice perigonadal fat were paraffin-embedded and the 4-µm tissue sections were cut from the paraffin blocks on the slides by Pathology core of the Institute of Biomedical Sciences, Academia Sinica. The slides were conducted immunochemistry staining using primary antibodies for F4/80 (1:200; cat#H0005972-M01, Abnova) and digitized using an Olympus BX51 microscope combined with an Olympus DP72 camera and CellSens Standard 1.14 software (Olympus, Germany).

### Hepatic triglycerides content measurement

Approximately 80 mg of liver tissue was homogenized in 1800 μl of chloroform/methanol (2/1). Then, 360 μl of H_2_O was added. The homogenates were centrifuged at 2000 rpm for 10 min. The lower 200-μl layer was added with 100 μl of chloroform with 4% Triton X-100 and dried in a chemical hood. The dried pellet was redissolved with 200 μl of H_2_O and was determined the triglyceride concentrations with Wako TG LabAssay kit (cat. no. 290-63701, Wako).

### High-throughput compound screening (HTS)

High-throughput compound screening was conducted at the core service in the Genomics Research Center (GRC) of Academia Sinica, Taiwan. The GRC 120K ReSet comprises more than 125,000 compounds, selected as representatives by structural similarity clustered from the 2M GRC compound library. The ReSet was arrayed in 1,536-well plates as single compounds at 1 mM in 100% DMSO. The quality of all compounds was assured by the vendor (purity is greater than 90%) and was verified internally with 5% random sampling. Recombinant human PTGR2 protein was purified, and the screening was conducted using the NADPH-Glo Detection kit (Promega) to measure the amount of NADPH. The CV of HTS ranged from 4.8 % to 6.1 %, with a Z’ value of 0.7. The threshold was set as 1.5, resulting in ∼300 hits for further confirmation and determination of the half-maximal inhibitory concentration (IC_50_). Eight-point two-fold dilution of the compounds was prepared for IC_50_ determination and used in dose-dependent studies. The compounds showed dose-dependent increases of unused NADPH, indicating recombinant human PTGR2 enzymatic activity.

### Formulation for 15-keto-PGE2 for injection

For animal experiments, 15-keto-PGE2 was dissolved in liposome (40 mg/kg/day in 20 μL/g/day liposome, intraperitoneal injection twice daily) for the treatment compared to the vehicle groups (liposome vehicle 20 μL/g/day, intraperitoneal injection twice daily). We are grateful to Taiwan Liposome Company for designing and providing the liposome formulation.

### Computer modeling analysis

To evaluate the interaction among PTGR2 inhibitor BPRPT0245, 15-keto-PGE2, NADPH, and PTGR2, the X-ray structure of human PTGR2 (PDB ID: 2ZB4) was used. Ligand energy was minimized using PyRx program before docking (Dallakyan and Olson, 2015). Three-dimensional models were visualized using the PyMOL program (Rigsby et al., 2016).

### Nuclear Magnetic Resonance (NMR) spectroscopy

^15^N-labeled hPPARγ LBD protein was purified as described in Brust et al. (Brust R, *et al*. 2018) with some modifications. Briefly, hPPARγ LBD (residues 203–477 in isoform 1 numbering) cloned in a pET15b was expressed in *Escherichia coli* BL21(DE3) cells using minimal media (M9 supplemented with ^15^NH_4_Cl). After 48-hour incubation with 1mM IPTG at 18°C, cells were harvested and lysed by NanoLyzer N2 in lysis buffer (20 mM Tris pH 8.0, 0.5 M NaCl, 10% glycerol and 1 mM tris(2-carboxyethyl) phosphine (TCEP)) supplemented 5 mM imidazole, 2.5 mM MgCl_2_, 1X protease inhibitor cocktail (Roche), 1 IU/ml DNAse and 0.1 mg/ml Lysozyme. Lysates were cleared by centrifugation (10,000 × g, 1 h) and loaded onto 5 mL Histrap FF columns (GE Healthcare). After washing column with lysis buffer supplemented 5 mM imidazole, protein was eluted using lysis buffer with a gradient concentration of imidazole (from 5 mM to 300 mM in 30 minutes). For thrombin cleavage, protein was incubated at a 1:100 ratio with thrombin (Cas No. 27084601, Cytiva, USA) overnight at 4 °C in PBS buffer. Tag-cleaved protein was collected from the flow through of second His-trap resin purification and concentrated by 10 K-spin columns. Condensed protein was loaded onto Superdex 200 prep grade 10/30 column (GE Healthcare). The LBD samples following size exclusion chromatography was stored in 50 mM potassium chloride (pH 7.4), 20 mM potassium phosphate, 5 mM TCEP, and 0.5 mM EDTA. For ligand binding NMR experiments, ^15^N-labeled hPPARγ LBD at 150 µM was pre-incubated with a 2X molar excess of 15-keto Prostaglandin E2 (15- keto-PGE2) overnight at 4 °C. Before performance of NMR experiments, samples were supplemented with 10% D_2_O.

The NMR experiments were conducted at 298K on Bruker NEO 850MHz NMR spectrometer equipped with 5 mm triple resonance cryoprobe with Z-axis gradient. NMR data were then processed and analyzed using software Topspin4.3 (Bruker, Germany). Standard ^15^N-TROSY-HSQC experiments were used to identify binding of 15-keto PGE2 to hPPARγ LBD. Amide ^1^H/^15^N resonance assignments for apo-form were extracted from BMRB entry number 15518 (Lu *et al*, 2008). The ligand binding study was performed with 150 μM ^15^N-labeled hPPARγ LBD with 15-keto-PGE2 at molar ratio 1:1 or 1:2.

### Statistical analysis

All values were expressed as mean ± S.E.M. All reported sample sizes were biologically independent but not technically repeatedly measured. Skewed data was logarithmized to approximate normal distribution. Comparisons between two separate groups were performed using Student t-tests. Comparisons among multiple groups were conducted using a one-way analysis of variance with post hoc analyses. Statistical analyses were conducted using GraphPad Prism 8.0 and SAS 9. Energy expenditure between genotypes was analyzed using the generalized linear model according to international guidance (Speakman *et al*, 2013; Tschöp *et al*, 2011). Energy expenditure (expressed as kcal/hr) was regressed on body weight using the command “glm” implemented in STATA 14.0 (Speakman *et al*, 2013; Tschöp *et al*, 2011). Two-sided *p*-values < 0.05 were considered statistically significant.

### The paper explained

Problem: PPARγ is a master transcriptional regulator of systemic metabolism and energy balance. Synthetic agonists of PPARγ have been used to treat diabetes mellitus for decades. However, these synthetic agonists are associated with adverse effects, including weight gain, osteoporosis, and water retention. Moreover, the identity of endogenous physiological PPARγ ligands remains unclear.

Result: In this study, we provide comprehensive evidence that 15-keto-PGE2 is an endogenous physiological PPARγ ligand. Direct administration of 15-keto-PGE2, as well as genetic or pharmacological inhibition of PTGR2 (the enzyme responsible for degrading 15-keto-PGE2), prevented diet-induced obesity, improved glucose abnormalities, and reduced fatty liver without causing fluid retention or osteoporosis.

Impact: Our findings highlight the importance of endogenous bioactive lipids as a promising avenue for treating diabetes, obesity, and fatty liver.

## Supporting information

Appedix data

## Data availability

All data needed to evaluate the conclusions in the paper are presented in the paper and/or the Appendix data. The data can be provided by owner of data pending scientific review and a completed material transfer agreement. Requests for the data should be submitted to

## Acknowledgments

We thank the technical services provided by the Transgenic Mouse Model Core Facility of the National Core Facility for Biopharmaceuticals, Ministry of Science and Technology, Taiwan and the Animal Resources Laboratory of National Taiwan University Centers of Genomic and Precision Medicine for generation of *Ptgr2* knockout mice. LTQ-Orbitrap data and additional technical assistance were performed by the Metabolomics Core Facility in the Scientific Instrument Center. The ultrahigh performance liquid chromatography and the ion trap-orbitrap mass spectrometer for quantification were supported by the Metabolomics Core Laboratory of the Agricultural Biotechnology Research Center, Academia Sinica, Taiwan. We thank the Taiwan Animal Consortium (MOST 107-2319-B-001-002) and the Taiwan Mouse Clinic for technical support in indirect calorimetry, body composition, tissue fixation and slide sectioning, and H&E stain experiment. We thank the Common Mass Spectrometry Facilities for Proteomics and Protein Modification Analysis of the Institute of Biological Chemistry, Academia Sinica for native mass spectrometry, which is supported by the Academia Sinica Core Facility and Innovative Instrument Project (AS-CFII-111-209). We thank the Department of Nuclear Medicine of National Taiwan University Hospital for small animal PET CT. The CRISPR/Cas9 and lentiviral system was provided by the RNA Technique and Gene Manipulation Core at the Genome Research Center, Academia Sinica, Taiwan. NMR data were collected in High-Field NMR Center (HFNMRC) in Academia Sinica which is funded by Academia Sinica Core Facility and Innovative Instrument Project (AS-CFII-111-214). The drug screening was conducted by the ultra-high throughput screening core service of the Genome Research Center, Academia Sinica, Taiwan. We also thank the pathological core service of the National Laboratory Animal Center. This work is supported by the Ministry of Science and Technology, Taiwan (104-2314-B-002-219-MY3, 106- 2314-B-002-137-MY3, 106-2321-B-002-040, 107-2321-B-002-067) (L.M.C.) National Taiwan University and National Taiwan University Hospital, Taiwan (UN105-0072, UN109-008). (L.M.C.)

## Author contributions

**Yi-Cheng Chang**: Conceptualization; methodology; investigation; visualization; project administration; writing – original draft. **Meng-Lun Hsieh**: Conceptualization; methodology; performing experiments; investigation; writing – review & editing. **Hsiao-Lin Lee**: Methodology; investigation; performing experiments; project administration; writing – review & editing. **Siow-Wey Hee**: Methodology; investigation; project administration. **Chi-Fon Chang:** NMR experiments. **Hsin-Yung Yen**: Methodology; investigation. **Yi-An Chen**: Methodology; investigation. **Yet-Ran Chen**: Methodology; investigation, writing – original draft. **Ya-Wen Chou**: Methodology; investigation; writing – original draft. **Fu-An Li**: Methodology; investigation. **Yi-Yu Ke**: Methodology; investigation. **Shih-Yi Chen**: Methodology; investigation; project administration. **Ming-Shiu Hung**: Methodology; investigation. **Alfur Fu-Hsin Hung**: Methodology; investigation. **Jing-Yong Huang**: Methodology; investigation; visualization. **Chu- Hsuan Chiu**: Writing – original draft; writing – review & editing. **Shih-Yao Lin**: Methodology; investigation. **Sheue-Fang Shih**: Methodology; investigation. **Chih-Neng Hsu**: Investigation. **Juey-Jen Hwang**: Investigation. **Teng-Kuang Yeh**: Methodology; investigation. **Ting-Jen Rachel Cheng**: Methodology; investigation. **Karen Chia-Wen Liao**: Methodology; investigation. **Daniel Laio**: Visualization **Chun-Mei Hu:** Methodology; investigation. **Shu-Wha Lin**: Methodology; investigation. **Tzu-Yu Chen**: Methodology; investigation. **Ulla Vogel**, **Daniel Saar**, **Birthe B. Kragelund:** providing NMR raw data. **Lun Kelvin Tsou**: Methodology; investigation; project administration. **Yu-Hua Tseng**: Conceptualization; supervision; writing – review & editing. **Lee-Ming Chuang**: Conceptualization; funding acquisition; supervision; writing – review & editing.

## Disclosure and competing interest statement

All authors declare that they have no conflict of interest.

## Expanded View Figures

**Figure EV1.**
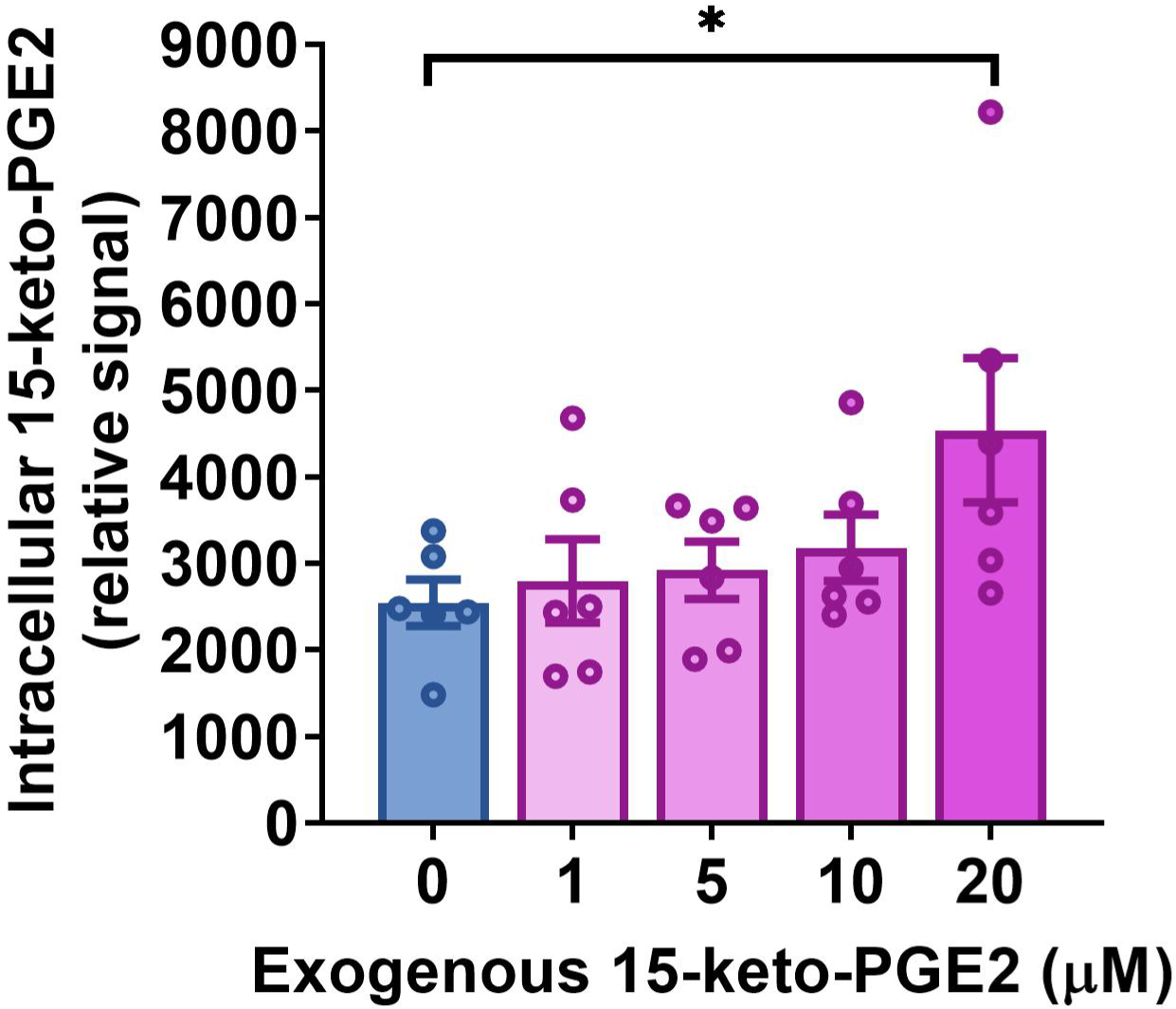
Relative intracellular 15-keto-PGE2 levels in cultured 3T3-L1 cells treated with exogenous 15-keto-PGE2 of different concentrations. Data information: All data are presented as mean and standard error (S.E.M.). Statistical significance was calculated by one-way analyses of variance (ANOVA) with Tukey’s post hoc test. \**p* < 0.05. Source data are available online for this figure.

**Figure EV2.**
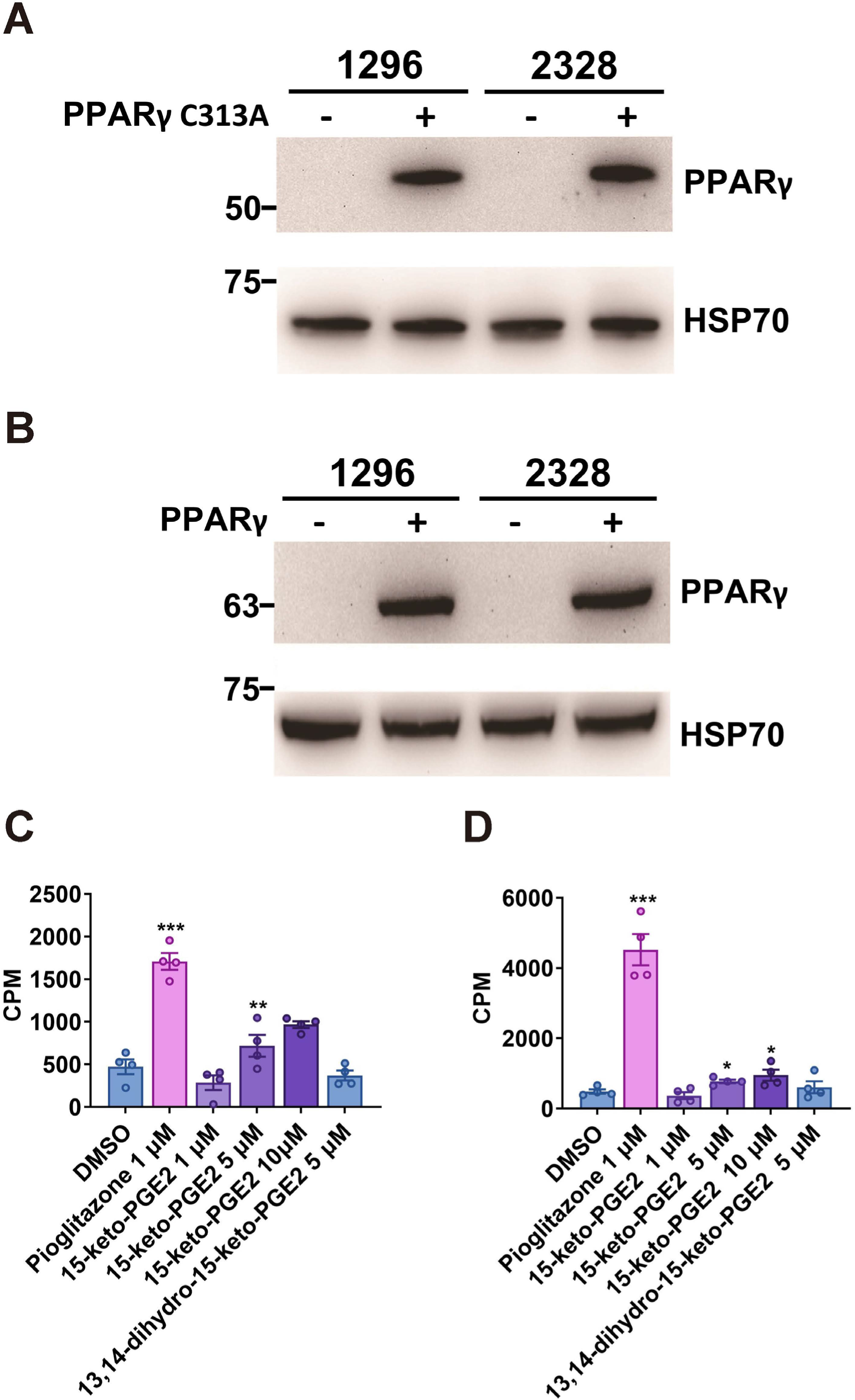
Immunoblots showing PPARγ expression in PPARγ-null 3T3-L1 clones #1296 and #2328 using the CRISPR techniques and then overexpress (A) mutant PPARγ2 (C313A) or (B) wild-type PPARγ2. 15-keto-PGE2 enhanced insulin-stimulated glucose uptake in clone #2328 rescue with (C) wild-type or (D) mutant PPARγ2 (C313A). Data information: All data are presented as mean and standard error (S.E.M.). Statistical significance was calculated by one-way analyses of variance (ANOVA) with Tukey’s post hoc test and two-sample independent *t*-test (C, D). \**p* < 0.05, ** *p* < 0.01, *** *p* < 0.001. Source data are available online for this figure.

**Figure EV3.**
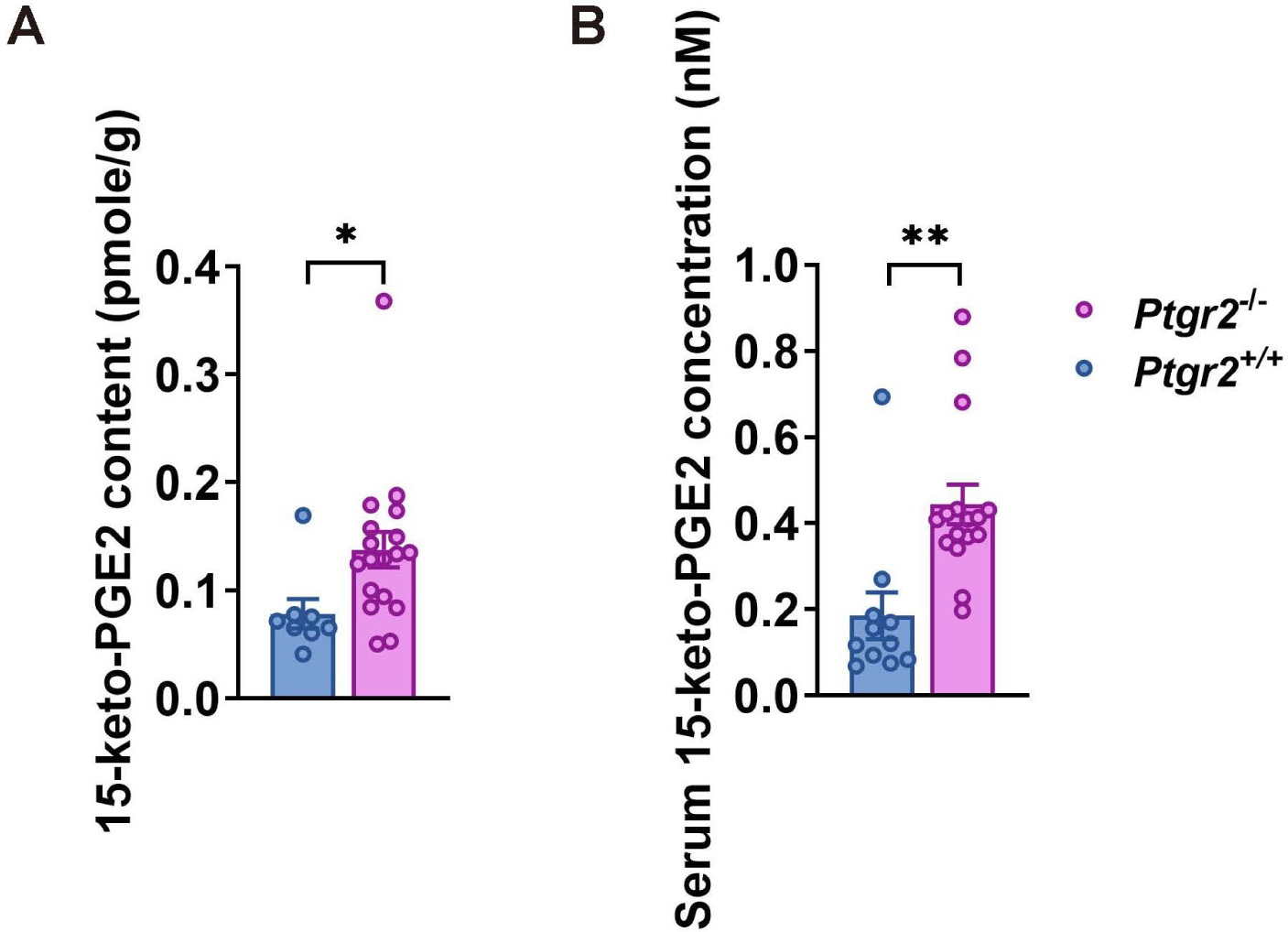
(A) Relative serum 15-keto-PGE2 level and (B) relative15-keto-PGE2 content in perigonadal fat (n=8:18) of *Ptgr2*^-/-^ and *Ptgr2^+/+^* mice on high-fat high-sucrose diet (HFHSD). Data information: All data are presented as mean and standard error (S.E.M.). Statistical significance was calculated by two-sample independent *t*-test in (A, B). \**p* < 0.05, ** *p* < 0.01. Source data are available online for this figure.

**Figure EV4.**
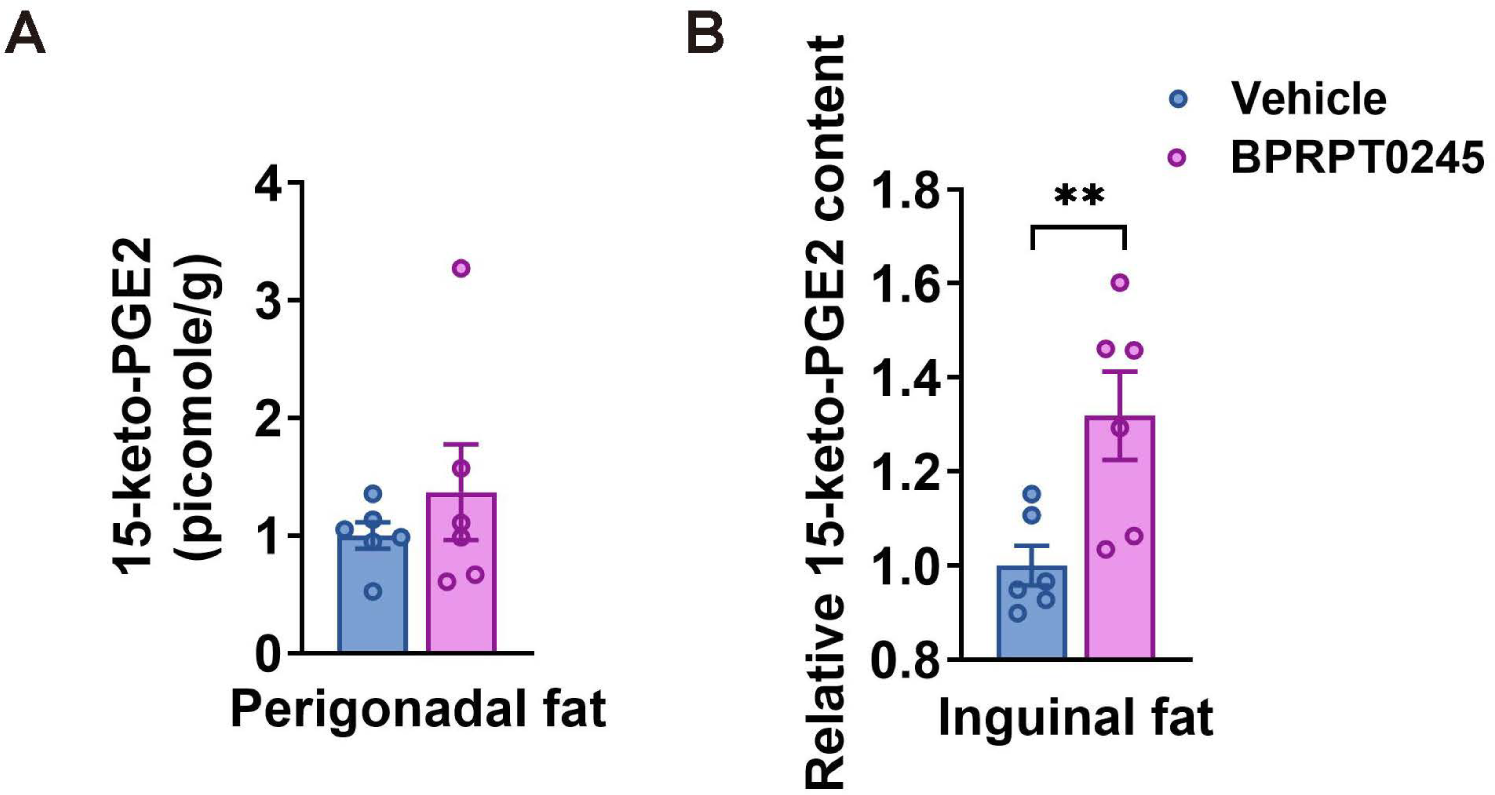
(A) Relative 15-keto-PGE2 contents in perigonadal fat (n=6:6) and (B) inguinal fat (n=6:6) after oral gavage of BPRPT0245 (100 mg/kg/day) for 4 days. Samples are harvested 2 hours after oral gavage of the latest dose. Data information: All data are presented as mean and standard error (S.E.M.). Statistical significance was calculated by two-sample independent *t*-test (A, B). \*\**p* < 0.01. Source data are available online for this figure.

**Figure EV5.**
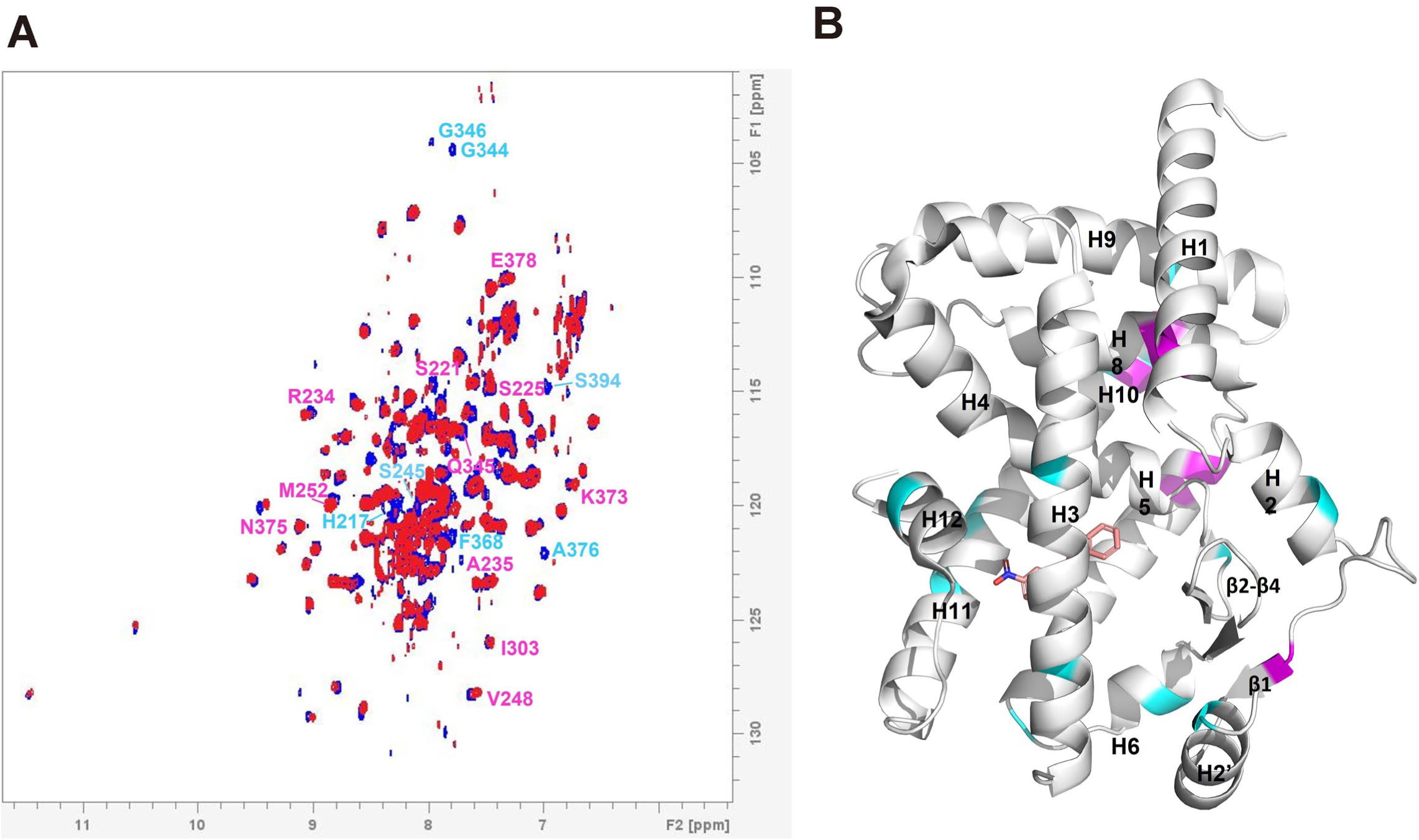
Comparison between 2D [^1^ H,^15^ N]-TROSY-HSQC NMR (nuclear magnetic resonance) spectra of apo-form and 15-keto-PGE2 bound PPARγ LBD (ligand binding domain). (A) Comparison between 2D [^1^ H,^15^ N]-TROSY-HSQC NMR spectra of apo-form and 15-keto-PGE2 bound PPARγ LBD (ligand binding domain). Cyanide color indicates missing peak. magenta color indicates chemical shift with Δδ > 0.05. (B) NMR missing peak (cyanide color) and chemical shift (magenta color) mapped onto PPARγ LBD structure.

## Notes

### Competing Interest Statement

The authors have declared no competing interest.

### Summary of Updates

We had added additioanl coauthors and associated information.

## References

Ahmadian M, Suh JM, Hah N, Liddle C, Atkins AR, Downes M, Evans RM (2013) PPARγ signaling and metabolism: the good, the bad and the future. Nat Med 19(5):557–66

Ardenkjær-Skinnerup, J., Saar, D., Petersen, P. S. S., Pedersen, M., Svingen, T., Kragelund, B. B., Hadrup, N., Ravn-Haren, G., Emanuelli, B., Brown, K. A., & Vogel, U. (2024). PPARγ antagonists induce aromatase transcription in adipose tissue cultures. Biochemical pharmacology, 222, 116095.

Brust R, Shang J, Fuhrmann J, Mosure SA, Bass J, Cano A, Heidari Z, Chrisman IM, Nemetchek MD, Blayo AL, Griffin PR, Kamenecka TM, Hughes TS, Kojetin DJ (2018). A structural mechanism for directing corepressor-selective inverse agonism of PPARγ. Nat Commun. 9(1):4687.

Bruning JB, Chalmers MJ, Prasad S, Busby SA, Kamenecka TM, He Y, Nettles KW, Griffin PR (2007) Partial agonists activate PPARgamma using a helix 12 independent mechanism. Structure 15(10):1258–71

Capelli D, Cerchia C, Montanari R, Loiodice F, Tortorella P, Laghezza A, Cervoni L, Pochetti G, Lavecchia A (2016) Structural basis for PPAR partial or full activation revealed by a novel ligand binding mode. Sci Rep. 6:34792

Cassard-Doulcier AM, Gelly C, Fox N, Schrementi J, Raimbault S, Klaus S, Forest C, Bouillaud F, Ricquier D (1993) Tissue-specific and beta-adrenergic regulation of the mitochondrial uncoupling protein gene: control by cis-acting elements in the 5’-flanking region. Mol Endocrinol 7(4):497–506

Chen IJ, Hee SW, Liao CH, Lin SY, Su L, Shun CT, Chuang LM (2018) Targeting the 15-keto-PGE_2_-PTGR2 axis modulates systemic inflammation and survival in experimental sepsis. Free Radic Biol Med 115:113–126

Cheng C, Nakamura A, Minamimoto R, Shinoda K, Tateishi U, Goto A, Kadowaki T, Terauchi Y, Inoue T (2011) Evaluation of organ-specific glucose metabolism by ¹⁸F-FDG in insulin receptor substrate-1 (IRS-1) knockout mice as a model of insulin resistance. Ann Nucl Med 25(10):755–61

Chhonker YS, Kanvinde S, Ahmad R, Singh AB, Oupický D, Murry DJ (2021) Simultaneous Quantitation of Lipid Biomarkers for Inflammatory Bowel Disease Using LC-MS/MS. Metabolites 11(2):106

Choi JH, Banks AS, Estall JL, Kajimura S, Boström P, Laznik D, Ruas JL, Chalmers MJ, Kamenecka TM, Blüher M, Griffin PR, Spiegelman BM (2010). Anti-diabetic drugs inhibit obesity-linked phosphorylation of PPARgamma by Cdk5. Nature 466(7305):451–6.

Choi JH, Banks AS, Kamenecka TM, Busby SA, Chalmers MJ, Kumar N, Kuruvilla DS, Shin Y, He Y, Bruning JB et al (2011) Antidiabetic actions of a non-agonist PPARγ ligand blocking Cdk5- mediated phosphorylation. Nature 477(7365):477–81

Chou WL, Chuang LM, Chou CC, Wang AH, Lawson JA, FitzGerald GA, Chang ZF (2007) Identification of a novel prostaglandin reductase reveals the involvement of prostaglandin E2 catabolism in regulation of peroxisome proliferator-activated receptor gamma activation. J Biol Chem 282(25):18162–18172

Chrisman IM, Nemetchek MD, de Vera IMS, et al. Defining a conformational ensemble that directs activation of PPARγ. Nat Commun. 2018;9(1):1794.

Dias MMG, Batista FAH, Tittanegro TH, de Oliveira AG, Le Maire A, Torres FR, Filho HVR, Silveira LR, Figueira ACM (2020) PPARγ S273 Phosphorylation Modifies the Dynamics of Coregulator Proteins Recruitment. Front Endocrinol (Lausanne). 27(11):561256.

Dallakyan S, Olson AJ (2015) Small-molecule library screening by docking with PyRx. Methods Mol Biol 1263:243–50

DePaoli AM, Higgins LS, Henry RR, Mantzoros C, Dunn FL, INT131-007 Study Group (2014) Can a selective PPARγ modulator improve glycemic control in patients with type 2 diabetes with fewer side effects compared with pioglitazone? Diabetes Care 37(7):1918–23

Forman BM, Tontonoz P, Chen J, Brun RP, Spiegelman BM, Evans RM (1995) 15-Deoxy-delta 12, 14-prostaglandin J2 is a ligand for the adipocyte determination factor PPAR gamma. Cell 83(5):803–12

Ding L, Zhou J, Ye L, Sun Y, Jiang Z, Gan D, Xu L, Luo Q, Wang G (2020). PPAR-γ Is Critical for HDAC3-Mediated Control of Oligodendrocyte Progenitor Cell Proliferation and Differentiation after Focal Demyelination. Mol Neurobiol. 57(11):4810–4824.

Harmon GS, Dumlao DS, Ng DT, Barrett KE, Dennis EA, Dong H, Glass CK (2010) Pharmacological correction of a defect in PPAR-gamma signaling ameliorates disease severity in Cftr-deficient mice. Nat Med 16(3):313–8

Harmon GS, Lam MT, Glass CK (2011) PPARs and lipid ligands in inflammation and metabolism. Chem Rev 111(10):6321–40

Hughes TS, Chalmers MJ, Novick S, Kuruvilla DS, Chang MR, Kamenecka TM, Rance M, Johnson BA, Burris TP, Griffin PR (2012) Ligand and receptor dynamics contribute to the mechanism of graded PPARγ agonism. Structure 20(1):139–50

Hughes TS, Giri PK, de Vera IM, Marciano DP, Kuruvilla DS, Shin Y, Blayo AL, Kamenecka TM, Burris TP, Griffin PR (2014) An alternate binding site for PPARγ ligands. Nat Commun 5:3571

Itoh T, Fairall L, Amin K, Inaba Y, Szanto A, Balint BL, Nagy L, Yamamoto K, Schwabe JW (2008) Structural basis for the activation of PPARgamma by oxidized fatty acids. Nat Struct Mol Biol 15(9):924–31

Irwin S, Karr C, Furman C, Tsai J, Gee P, Banka D, Wibowo AS, Dementiev AA, O’Shea M, Yang J, Lowe J, Mitchell L, Ruppel S, Fekkes P, Zhu P, Korpal M, Larsen NA. (2022). Biochemical and structural basis for the pharmacological inhibition of nuclear hormone receptor PPARγ by inverse agonists. J Biol Chem 298(11), 102539.

Inazumi T, Yamada K, Shirata N, Sato H, Taketomi Y, Morita K, Hohjoh H, Tsuchiya S, Oniki K, Watanabe T, Sasaki Y, Oike Y, Ogata Y, Saruwatari J, Murakami M, Sugimoto Y (2020). Prostaglandin E2-EP4 axis promotes lipolysis and fibrosis in adipose tissue leading to ectopic fat deposition and insulin resistance. Cell Rep. 33(2):108265.

Jiang X, Ye X, Guo W, Lu H, Gao Z (2014) Inhibition of HDAC3 promotes ligand-independent PPARgamma activation by protein acetylation. J Mol Endocrinol. 53(2):191–200.

Kliewer SA, Sundseth SS, Jones SA, Brown PJ, Wisely GB, Koble CS, Devchand P, Wahli W, Willson TM, Lenhard JM et al (1997) Fatty acids and eicosanoids regulate gene expression through direct interactions with peroxisome proliferator-activated receptors alpha and gamma. Proc Natl Acad Sci U S A 94(9):4318–23

Konda VR, Desai A, Darland G, Grayson N, Bland JS (2014) KDT501, a derivative from hops, normalizes glucose metabolism and body weight in rodent models of diabetes. PLoS One 9(1):e87848

Kozak UC, Kopecky J, Teisinger J, Enerbäck S, Boyer B, Kozak LP (1994) An upstream enhancer regulating brown-fat-specific expression of the mitochondrial uncoupling protein gene. Mol Cell Biol 14(1):59–67

Krey G, Braissant O, L’Horset F, Kalkhoven E, Perroud M, Parker MG, Wahli W (1997) Fatty acids, eicosanoids, and hypolipidemic agents identified as ligands of peroxisome proliferator-activated receptors by coactivator-dependent receptor ligand assay. Mol Endocrinol 11(6):779–91

Loening AM, Gambhir SS (2003) AMIDE: a free software tool for multimodality medical image analysis. Mol Imaging 2(3):131–7

Marso SP, Daniels GH, Brown-Frandsen K, Kristensen P, Mann JF, Nauck MA, Nissen SE, Pocock S, Poulter NR, Ravn LS et al (2016) Liraglutide and Cardiovascular Outcomes in Type 2 Diabetes. N Engl J Med 28;375(4):311-22

Miyamae Y (2021) Insights into Dynamic Mechanism of Ligand Binding to Peroxisome Proliferator-Activated Receptor γ toward Potential Pharmacological Applications. Biol Pharm Bull 44(9):1185–1195

Momcilovic M, Bailey ST, Lee JT, Zamilpa C, Jones A, Abdelhady G, Mansfield J, Francis KP, Shackelford DB (2018) Utilizing 18F-FDG PET/CT Imaging and Quantitative Histology to Measure Dynamic Changes in the Glucose Metabolism in Mouse Models of Lung Cancer. J Vis Exp 137:57167

Montanari R, Capelli D, Yamamoto K, Awaishima H, Nishikata K, Barendregt A, Heck AJR, Loiodice F, Altieri F, Paiardini A, Grottesi A, Pirone L, Pedone E, Peiretti F, Brunel JM, Itoh T, Pochetti G (2020). Insights into PPARγ Phosphorylation and Its Inhibition Mechanism. J Med Chem. 63(9):4811–4823.

Mottin M, Souza PC, Skaf MS (2015). Molecular Recognition of PPARγ by Kinase Cdk5/p25: Insights from a Combination of Protein-Protein Docking and Adaptive Biasing Force Simulations. J Phys Chem B. 119(26):8330–9.

Nagy L, Tontonoz P, Alvarez JG, Chen H, Evans RM (1998) Oxidized LDL regulates macrophage gene expression through ligand activation of PPARgamma. Cell 93(2):229–40

Niphakis MJ, Lum KM, Cognetta AB 3rd, Correia BE, Ichu TA, Olucha J, Brown SJ, Kundu S, Piscitelli F, Rosen H et al (2015) A Global Map of Lipid-Binding Proteins and Their Ligandability in Cells. Cell 161(7):1668–80

Nolte RT, Wisely GB, Westin S, Cobb JE, Lambert MH, Kurokawa R, Rosenfeld MG, Willson TM, Glass CK, Milburn MV (1998) Ligand binding and co-activator assembly of the peroxisome proliferator-activated receptor-gamma. Nature 395(6698):137–43

Ohno H, Shinoda K, Spiegelman BM, Kajimura S (2012) PPARγ agonists induce a white-to-brown fat conversion through stabilization of PRDM16 protein. Cell Metab 15(3):395–404

Parker CG, Galmozzi A, Wang Y, Correia BE, Sasaki K, Joslyn CM, Kim AS, Cavallaro CL, Lawrence RM, Johnson SR et al (2017) Ligand and Target Discovery by Fragment-Based Screening in Human Cells. Cell 168(3):527–541.e29

Rigsby RE, Parker AB (2016). Using the PyMOL application to reinforce visual understanding of protein structure. Biochem Mol Biol Educ.10;44(5):433–7.

Schopfer FJ, Lin Y, Baker PR, Cui T, Garcia-Barrio M, Zhang J, Chen K, Chen YE, Freeman BA (2005) Nitrolinoleic acid: an endogenous peroxisome proliferator-activated receptor gamma ligand. Proc Natl Acad Sci U S A 102(7):2340–5

Shang J, Mosure SA, Zheng J, Brust R, Bass J, Nichols A, Solt LA, Griffin PR, Kojetin DJ. (2020). A molecular switch regulating transcriptional repression and activation of PPARγ. Nature Commun, 11(1), 956.

Shiraki T, Kamiya N, Shiki S, Kodama TS, Kakizuka A, Jingami H (2005) Alpha,beta-unsaturated ketone is a core moiety of natural ligands for covalent binding to peroxisome proliferator-activated receptor gamma. J Biol Chem 280(14):14145–53

Soccio RE, Chen ER, Lazar MA (2014) Thiazolidinediones and the promise of insulin sensitization in type 2 diabetes. Cell Metab 20(4):573–91

Song YS, Lee DH, Yu JH, Oh DK, Hong JT, Yoon DY (2016) Promotion of adipogenesis by 15- (S)-hydroxyeicosatetraenoic acid. Prostaglandins Other Lipid Mediat. 123:1–8

Speakman JR, Fletcher Q, Vaanholt L (2013) The ‘39 steps’: an algorithm for performing statistical analysis of data on energy intake and expenditure. Dis Model Mech 6(2):293–301

Thangavel N, Al Bratty M, Akhtar Javed S, Ahsan W, Alhazmi HA (2017) Targeting peroxisome proliferator-activated receptors using thiazolidinediones: strategy for design of novel antidiabetic drugs. Int J Med Chem 2017:1069718

Toroitich EK, Ciancone AM, Hahm HS, Brodowski SM, Libby AH, Hsu KL (2021) Discovery of a cell-active SuTEx ligand of prostaglandin reductase 2. Chembiochem 22(12):2134–2139

Tschöp MH, Speakman JR, Arch JR, Auwerx J, Brüning JC, Chan L, Eckel RH, Farese RV Jr, Galgani JE, Hambly C et al (2011) A guide to analysis of mouse energy metabolism. Nat Methods 9(1):57–63

Waki H, Park KW, Mitro N, Pei L, Damoiseaux R, Wilpitz DC, Reue K, Saez E, Tontonoz P (2007) The small molecule harmine is an antidiabetic cell-type-specific regulator of PPARgamma expression. Cell Metab 5 (5):357–70

Waku T, Shiraki T, Oyama T, Fujimoto Y, Maebara K, Kamiya N, Jingami H, Morikawa K (2009) Structural insight into PPARgamma activation through covalent modification with endogenous fatty acids. J Mol Biol 385(1):188–99

Waku T, Shiraki T, Oyama T, Maebara K, Nakamori R, Morikawa K (2010) The nuclear receptor PPARγ individually responds to serotonin- and fatty acid-metabolites. EMBO J 29(19):3395–407

Wiviott SD, Raz I, Bonaca MP, Mosenzon O, Kato ET, Cahn A, Silverman MG, Zelniker TA, Kuder JF, Murphy SA et al (2019) Dapagliflozin and cardiovascular outcomes in Type 2 Diabetes. N Engl J Med 24;380(4):347-357

Wu YH, Ko TP, Guo RT, Hu SM, Chuang LM, Wang AH (2008) Structural basis for catalytic and inhibitory mechanisms of human prostaglandin reductase PTGR2. Structure 16(11):1714–23

Yamada M, Kita Y, Kohira T, Yoshida K, Hamano F, Tokuoka SM, Shimizu T (2015) A comprehensive quantification method for eicosanoids and related compounds by using liquid chromatography/mass spectrometry with high speed continuous ionization polarity switching. J Chromatogr B Analyt Technol Biomed Life Sci 995–996:74-84

Yu YH, Chang YC, Su TH, Nong JY, Li CC, Chuang LM (2013) Prostaglandin reductase-3 negatively modulates adipogenesis through regulation of PPARγ activity. J Lipid Res 54(9):2391–9

Zoete V, Grosdidier A, Michielin O (2007) Peroxisome proliferator-activated receptor structures: ligand specificity, molecular switch and interactions with regulators. Biochim Biophys Acta 1771(8):915–25

