## Supplementary material for "Identification of PTGR2 inhibitors as a new therapy for diabetes and obesity": Appedix data

### Appendix

#### Materials and Methods

##### Co-Immunoprecipitation

Lysates from the perigonadal fat of *Ptgr2*$^{+/+}$ and *Ptgr2*$^{-/-}$ mice were immunoprecipitated with a mouse anti-15-keto-$PGE_2$-cysteine-BSA antibody and then immunoblotted with a rat anti-PPARγ antibody (cat. no.sc-7273, Santa Cruz Biotechnology).

##### Computer modeling analysis

To illustrate the binding mode of 15-keto-PGE2, a covalent docking analyses were performed with the CovalentDock program (Ouyang et al., 2013). The human X-ray structure of human PPARγ (PDB ID: 5Y2O) (Lee et al., 2017) was employed to evaluate covalent binding with the small molecule 15-keto-PGE2. To mimic experimental results observed in mouse species, four amino acids (S302N, V307I, L435V, and Q454H) were mutated via computer modeling. After docking, molecular dynamics simulations were conducted using the BIOVIA 2017/Standard Dynamics Cascade program (BIOVIA, Inc., San Diego, CA) to observe how the 15-keto-PGE2 and PPARγ LBD complex behaves over time, eventually reaching equilibrium so that the interactions become stable and representative of a real biological environment. Three-dimensional models were visualized using the PyMOL program (Rigsby et al., 2016). This docking method was originally used to simulate the binding of 15-keto-PGE2 to the PPARγ LBD.

##### PPARγ phosphorylation assay in 3T3-L1 cell and adipose tissue

For cell-based assay, 3T3-L1 cells were induced differentiation one day after achieving 100% confluence with 0.5 mM IBMX, 1 μm dexamethasone and 10 μg/mL insulin in DMEM containing 10% FBS for 2 days and maintained differentiation in DMEM supplemented with 10 μg/mL insulin and 10% FBS for 4 days. The medium was changed every 2 days. On day 6, cells were starved in low-glucose DMEM with 0.25% non-fatty acid for 24 hrs. and then treated with indicated concentrations of 15-keto-PGE2 for 1 hr. Then TNFα was added to induce PPARγ phosphorylation in a concentration of 50 ng/ml for 15 minutes. Cells were harvested and lysed in RIPA buffer supplemented with 1X cOmplete™, EDTA-free Protease Inhibitor Cocktail (cat. no. 04693132001, Roche) and 1X PhosSTOP™ (cat. no. 4906845001, Roche).

For adipose tissue samples, eight-week old B6 mice were treated with or without 40 mg/kg/day of 15-keto-PGE2 for 3 weeks under HFHSD. The epididymal adipose tissues were obtained and lysed in RIPA buffer supplemented with 1X cOmplete™, EDTA-free Protease Inhibitor Cocktail (cat no.04693132001, Roche) and 1X PhosSTOP™ (cat. no. 4906845001, Roche).

The protein concentrations of tissue lysates were measured by Bradford reagent (cat. no. BP500-0006, Bio-Rad). The protein samples were mixed with 6X sample buffer (0.3 M Tris-HCl, pH 6.8, 0.6 M DTT, 12% SDS, 0.6% Bromophenel blue and 60% glycerol) and heated at 70°C for 5 minutes. 30 μg of each sample were separated by 10% SDS-PAGE gel and were transferred to PVDF membrane. The membrane was blocked by 3% BSA in Tris-buffered saline containing 0.1% Tween-20 for 1 hour at room temperature and incubated with primary antibody at 4°C overnight. Primary antibodies include anti-Ser273 phospho-PPARγ (1:1000, cat. no. BS-4888R, Bioss), anti-PPARγ (1:1500, cat. no.16643-1-AP, Proteintech) and anti-GAPDH (1:5000, cat. no. GTX100118, GeneTex). On the next day, the membranes were incubated with the anti-rabbit IgG HRP-secondary antibody (1:10000, cat. no. GTX213110-01, GeneTex) for 1h at RT. Immunoblots were developed using the Immobilon Forte Western HRP Substrate (cat. no. WBLUF0500, Lot Number: 231445, Millipore) and exposed by MultiGel-21 (cat. no. MGIS-21-C2, EBL). Protein expression of phospho-PPARγ was analyzed using ImageJ software and normalized against PPARγ and GAPDH.

**Cell-based PPARγ acetylation assay**

One day after achieving 100% confluence, 3T3-L1 cells were induced differentiation with 0.5 mM IBMX, 1 μm dexamethasone and 10 μg/mL insulin in DMEM containing 10% FBS for 2 days and maintained differentiation in DMEM supplemented with 10 μg/mL insulin and 10% FBS for 6 days. The medium was changed every 2 days. On day 8, cells were starved in low-glucose DMEM with 0.25% non-fatty acid for 24 hrs and then treated with indicated concentrations of 15-keto-PGE2 for 2 hrs. Cells were harvested and lysed in RIPA buffer supplemented with 1X cOmplete™, EDTA-free Protease Inhibitor Cocktail (cat. no.04693132001, Roche) and 1X PhosSTOP™ (cat. no.4906845001, Roche). The protein concentrations of cell lysates were measured by Bradford reagent (cat. no. BP500-0006, Bio-Rad). The protein samples were performed immunoprecipitation with magnetic beads (Protein G Mag Sepharose™, Cytiva). Briefly, 200 μg of protein samples were incubated in 500 μl of PBS buffer containing 1 μg of anti-PPARγ antibody or 1 μg of anti-rabbit IgG antibody with gentle rotation at 4°C overnight. On the next day, PBS-washed beads were incubated with overnight mixture of protein sample and antibody at 4°C for 1 hr with gentle rotation. Then the beads were washed three times with 1000 μl of PBS and harvested with centrifugation at 4°C and 10000 g for 5 minutes. 50 μl of 1X sample buffer (0.05 M Tris-HCl, pH 6.8, 0.1 M DTT, 2% SDS, 0.1% Bromophenel blue and 10% glycerol) were mixed with the beads and heated at 90°C for 5 minutes. 10 μl of each sample were separated by 10% SDS-PAGE gel and were transferred to PVDF membrane. The membrane was blocked by 10% skim milk in PBS containing 0.05 % Tween-20 for 1 hr at room temperature and incubated with primary antibody at 4°C overnight. Primary antibodies include anti-acetyl-lysine (1:1000, cat. no. 9441, Cell Signaling) and anti-PPARγ (1:1500, cat. no.16643-1-AP, Proteintech). On the next day, the membranes were incubated with the rabbit HRP-secondary antibody (1:10000, cat. no. GTX213110-01, GeneTex) for 1 hr at RT. Immunoblots were developed using the Millipore Immobilon Forte Western HRP Substrate (WBLUF0500, Lot Number: 231445) and exposed by MultiGel-21 (cat. no. MGIS-21-C2, EBL). Acetylation level of PPARγ was analyzed using Image J software and normalized against PPARγ.

**Pharmacokinetic studies**

Male ICR mice weighing 25-30 g each were obtained from BioLASCO, Taiwan Co., Ltd., Ilan, Taiwan for pharmacokinetic study. Briefly, a single dose of BRRPT245 was given intravenously at 2 mg/kg or oral gavage at 10 mg/kg to the mice. Blood samples were taken at 0 (immediately before dosing), 2, 5 (iv only), 15 and 30 min and at 1, 2, 4, 6, 8, 16 and 24 hr after administration. The $T_{1/2}$ (half-life), Cl (clearance), Vss (steady-state volume of distribution), AUC (area under curve), $C_{max}$ (maximum plasma concentration), $T_{max}$ (time taken to reach $C_{max}$), and F(bioavailability%) were calculated according to the published formula (Urso *et al*, 2002).

The steady levels of 15-keto-PGE2 and BPRPT0245 in plasma and fat tissue were performed by giving C57BL6/J mice 4 doses of BPRPT0245 (100mg/kg/day) once daily for4 days. Blood and perfused fat tissues were harvested2 hours after the last dose.

The levels of 15-keto-PGE2 and BPRPT0245 were determined by high-performance liquid chromatography and tandem mass spectrometry. The HPLC system consisted of an Agilent 1200 series LC System with

BinPump (Waldbronn, Germany) and a Agilent Eclipse XDB C8 (3.0×150 mm, 5 μm) interfaced to a Sciex API 4000 tandem mass spectrometer. Mobile phase consisted of 10 mM ammonium acetate containing 0.1% of formic acid (Solvent A) and acetonitrile (Solvent B). For determination of BPTPT0245, the following stepwise gradient system was used: 85% A (0-0.5 min), 85% A to 15% A (0.6-3.0 min), 5% A (3.1-5.0 min). Total running time was 5 min. The retention times of BPRPT0245 and BPR0L187 (internal standard) (IS) were 2.21 and 2.42 min, respectively. Data acquisition was via selected ion monitoring (SIM). The collision energy was +37V for the analyte and +38.5V for IS, respectively. The ions monitored for BPRPT0245 were m/z 447/189 and m/z 357/195, respectively. For determination of 15-keto-$PGE_2$, the gradient system was used: 90% A (0-0.5 min), 90% A to 10% A (0.5-4.5 min), 90% A (4.5-6 min). Total running time was 6 min. The retention times of 15-keto-PGE2 and 13,14-dihydro-15-keto PGE2-D4 were3.27 and 3.33 min, respectively. Data acquisition was via SIM. The collision energy was -37V for 15-keto-PGE2 and -20V for 13,14-dihydro-15-keto PGE2-D4, respectively. The ions monitored for 15-keto-PGE2 and 13,14-dihydro-15-keto-PGE2-D4 were m/z 349.0/331.2 and m/z 355.1/337, respectively.

**Sample preparation**

To prepare 15-keto-PGE2 serum samples, aliquot (60 μL) of serum was mixed with 120 μL of acetonitrile, 0.1% formic acid, and 0.005% butylated hydroxytoluene solution containing 20 ng/mL of 13,14-dihydro-15-keto PGE2-D4 as the IS. The mixture was vortexed for 30 seconds and then centrifuged at 15,000 rpm for 20 min in an Eppendorf Model 5417c centrifuge at room temperature. An aliquot (25 μL) of the mixture was then injected onto to LC-MS/MS.

To prepare the BPRPT0245 plasma samples, 50 μL of the plasma sample mixed with 100 μL acetonitrile containing 250 μL of BPR0L187. The mixture was vortexed for 30 seconds and then centrifuged at 15,000 rpm for 20 min in an Eppendorf Model 5417c centrifuge at room temperature. An aliquot (5 μL) of the mixture was then injected onto to LC-MS/MS.

**Necropsy, gross Examinations, and Histopathological examination**:

The animals were sacrificed by exsanguination under anesthesia with pure carbon dioxide.

At necropsy, tissue samples of the submitted mice were collected and preserved in 10% neutral buffered formalin. The required tissues/organs of animals were trimmed, processed, and embedded in paraffin. Sections 3-5 μm in thickness were cut and put on slides for hematoxylin and eosin (H&E) staining. The histopathological evaluation was performed on the submitted tissues. Severity of lesions was graded according to the methods described (Shackelford *et al*, 2002). Degrees of lesions were graded histopathologically from one to five depending on severity (1 = minimal; 2 = slight; 3 =moderate; 4 = moderately severe; 5 = severe/high).

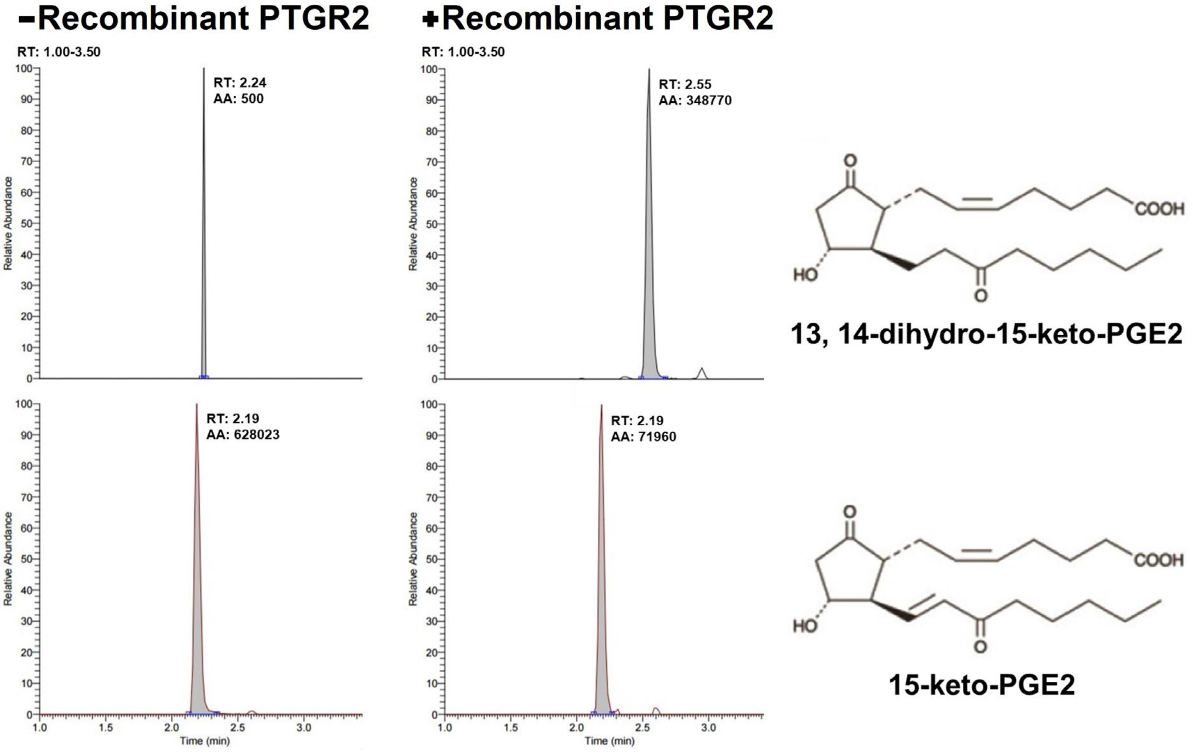

**Appendix Figure S1.** Orbitrap LC-MS/MS showing 15-keto-PGE2 is converted to 13,14-dihydro-15-keto-PGE2 by recombinant human PTGR2 protein.

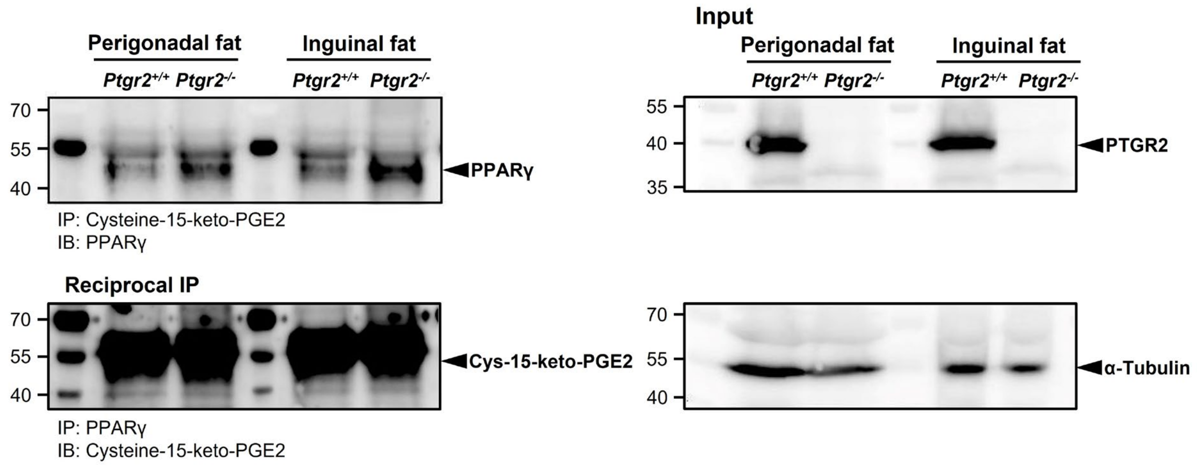

**Appendix Figure S2.** Reciprocal co-immunoprecipitation between 15-keto-PGE2 and PPARγ in mouse perigonadal white adipose tissues of *Ptgr2*$^{+/+}$ and *Ptgr2*$^{-/-}$ mice. A single band corresponding to murine PPARγ (57 kDa) was detected (upper panel), confirming the covalent binding of PPARγ to 15-keto-PGE2. When protein lysates were reciprocally immunoprecipitated with the anti-PPARγ antibody and immunoblotted with the anti-15-keto-PGE2-cysteine-BSA antibody. However, when protein lysates were reciprocally immunoprecipitated with the anti-PPARγ antibody and immunoblotted with the anti-15-keto-PGE2-cysteine-BSA antibody, the mPPARγ band was obscured by the IgG heavy chain, which is enriched in tissue (lower panel).

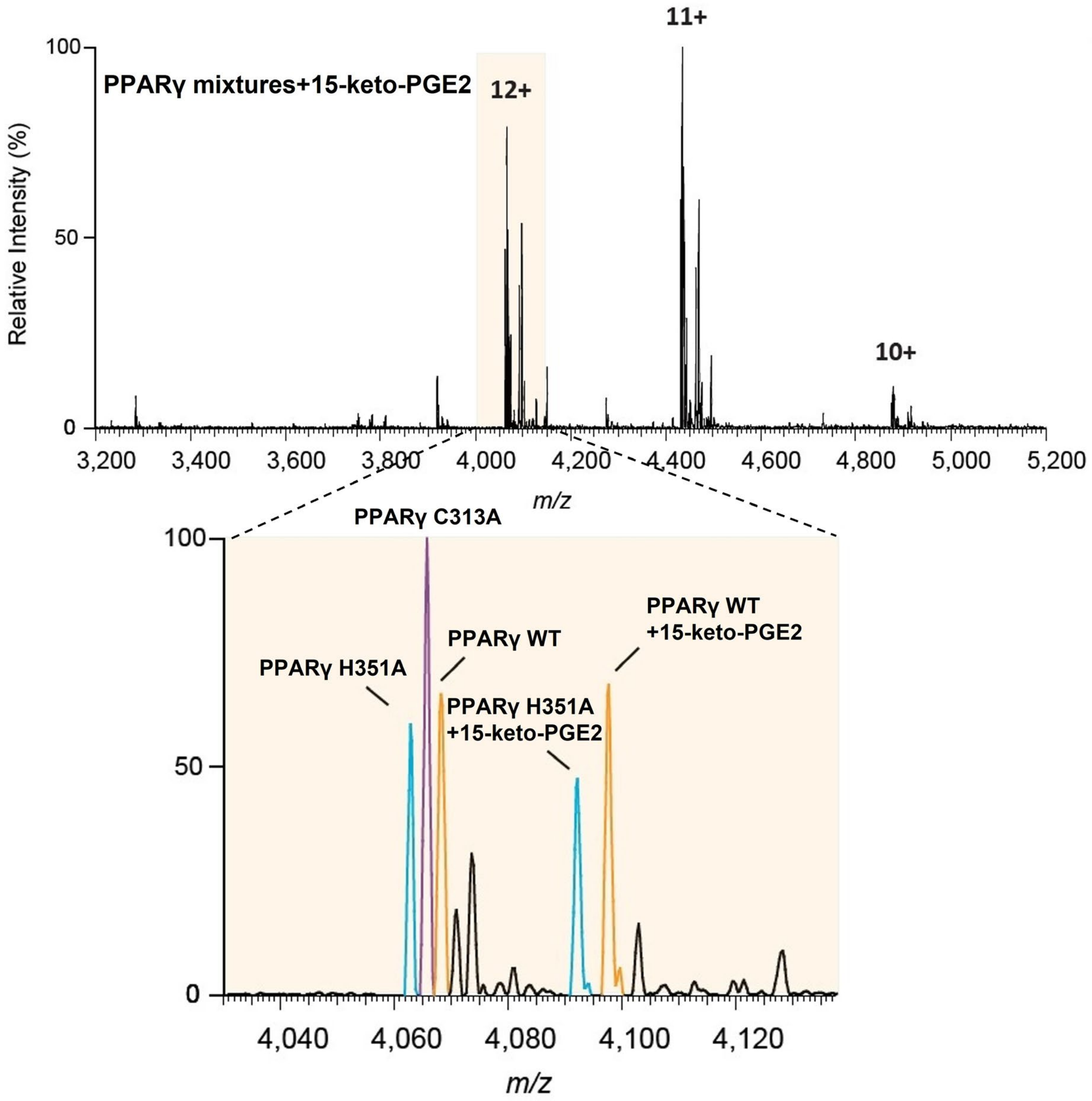

**Appendix Figure S3.** Examination of 15-keto-PGE2 to wild-type murine PPARγ (mPPARγ) and mutants by native mass spectrometry. The H351A mutants, C313A mutants and wild-type mPPARγ are denoted in blue, magenta and orange respectively. The 15-keto-PGE2 was analyzed by measuring the increased mass in individual proteins. The binding of 15-keto-PGE2 was analyzed at the indicated charge state (12+), highlighted in the vanilla background.

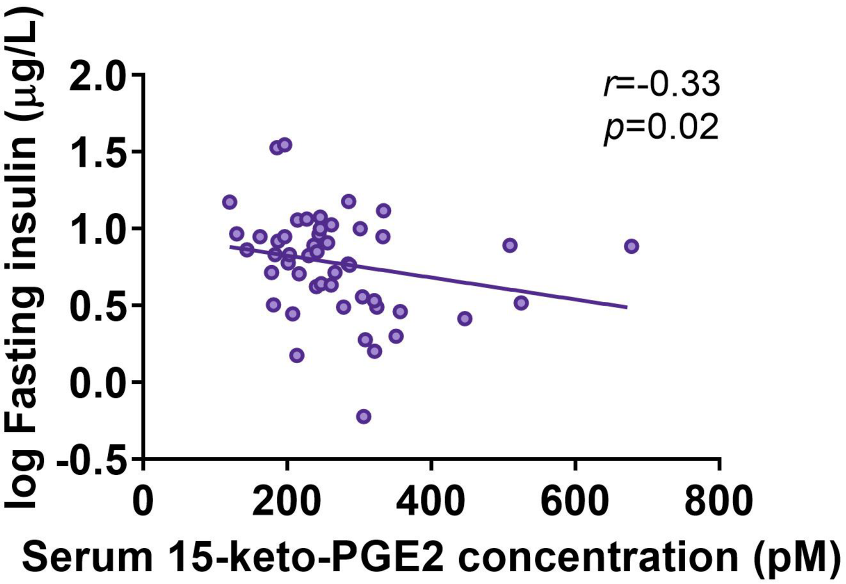

**Appendix Figure S4.** The inverse association between serum 15-keto-PGE2 levels and log fasting insulin levels in 50 non-diabetic human subjects. Data information: Statistical significance was calculated by Spearman's correlation. Source data are available online for this figure.

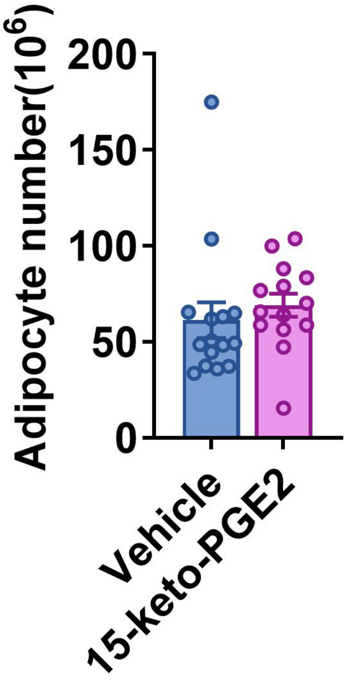

**Appendix Figure S5.** The number of adipocytes of perigonadal fat in 15-keto-PGE2-treated mice compared with control mice. Data information: All data are presented as mean and standard error (S.E.M.). Statistical significance was calculated by two-sample independent *t*-test. Source data are available online for this figure.

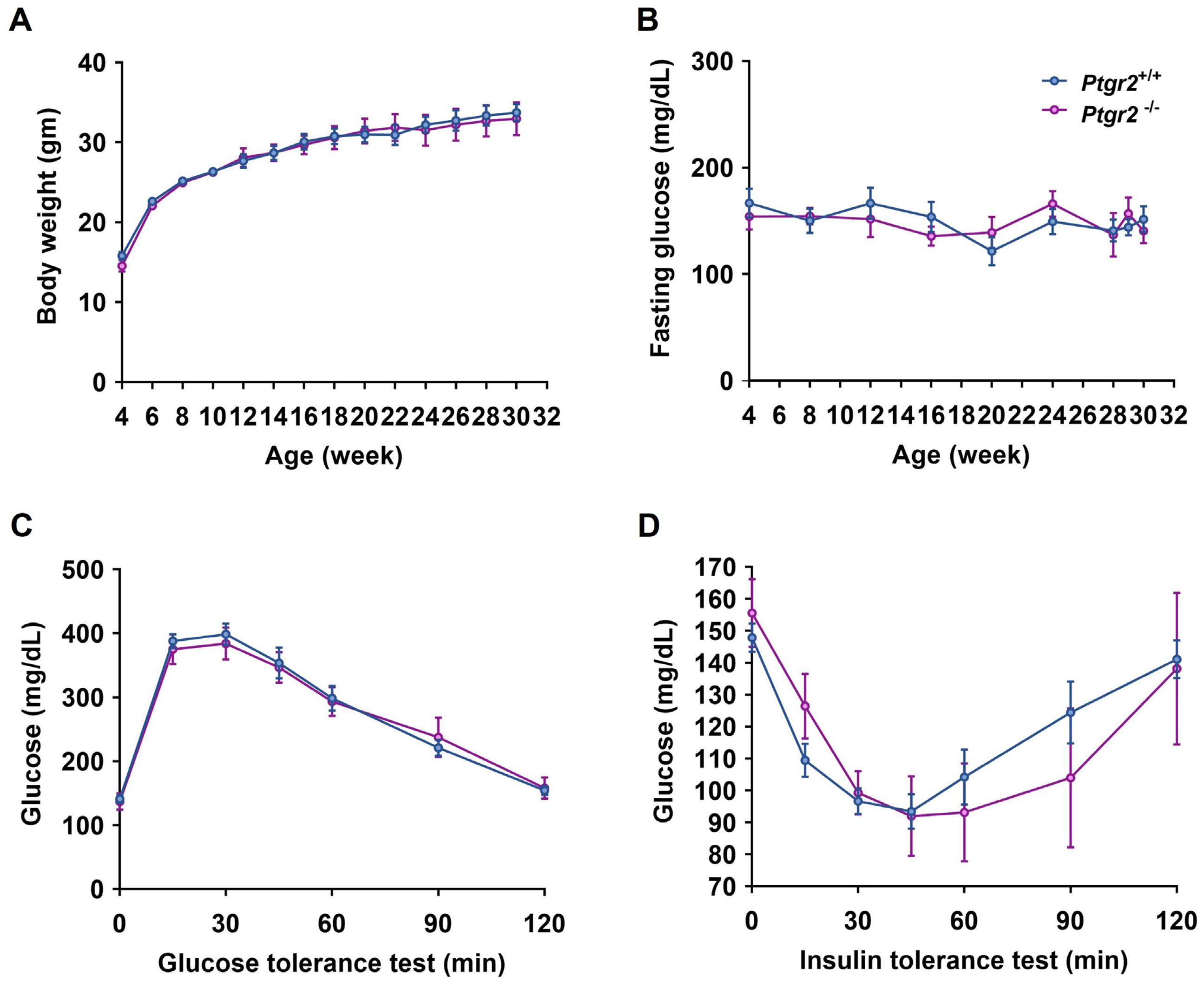

**Appendix Figure S6.** Body weight (n=44:42) (A), fasting glucose (n=25:26) (B), glycemic levels during intraperitoneal glucose tolerance test a (n=25:26) (C) and insulin tolerance test of *Prgr2*$^{-/-}$ and *Prgr2*$^{+/+}$ mice on chow diet (n=25:26) (D) at the age of 30 weeks.

Data information: All data are presented as mean and standard error (S.E.M.). Statistical significance was calculated by two-sample independent *t*-test. Source data are available online for this figure.

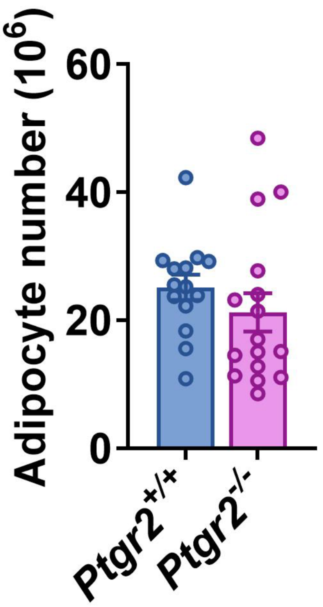

**Appendix Figure S7.** The number of adipocytes of in the perigonadal fat of *Ptgr2*$^{-/-}$mice compared with control mice. Data information: All data are presented as mean and standard error (S.E.M.). Statistical significance was calculated by two-sample independent *t*-test. Source data are available online for this figure.

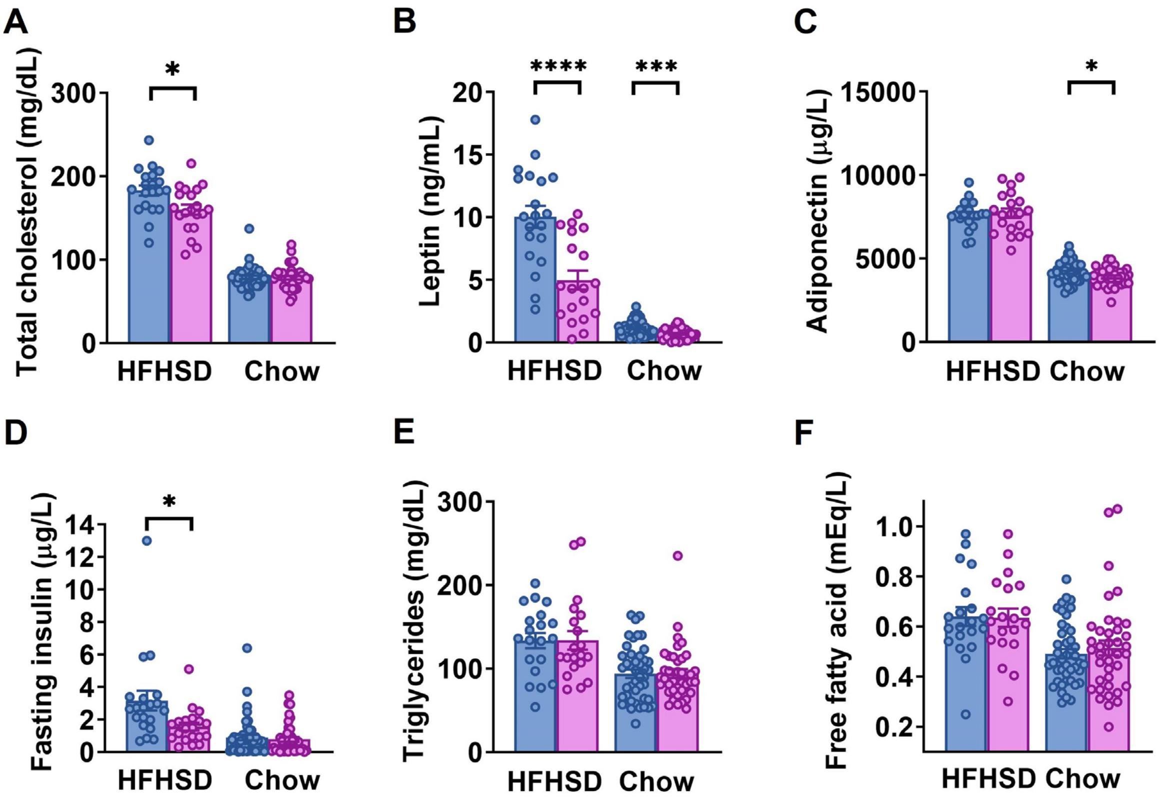

**Appendix Figure S8.** (A) Total cholesterol, (B) leptin, (C) adiponectin, (D) fasting insulin, (E) triglycerides and (F) free fatty acid of *Ptgr2*$^{-/-}$ and *Ptgr2*$^{+/+}$ mice on chow (n=44:38) or high-fat high-sucrose diet (HFHSD) (n=20:20) at the age of 30 weeks. The blue dot represents *Ptgr2*$^{+/+}$ mice and the purple dot represents *Ptgr2*$^{-/-}$ mice. Data information: All data are presented as mean and standard error (S.E.M.). Statistical significance was calculated by two-sample independent t-test. * $p < 0.05$, *** $p < 0.001$, **** $p < 0.0001$. Source data are available online for this figure.

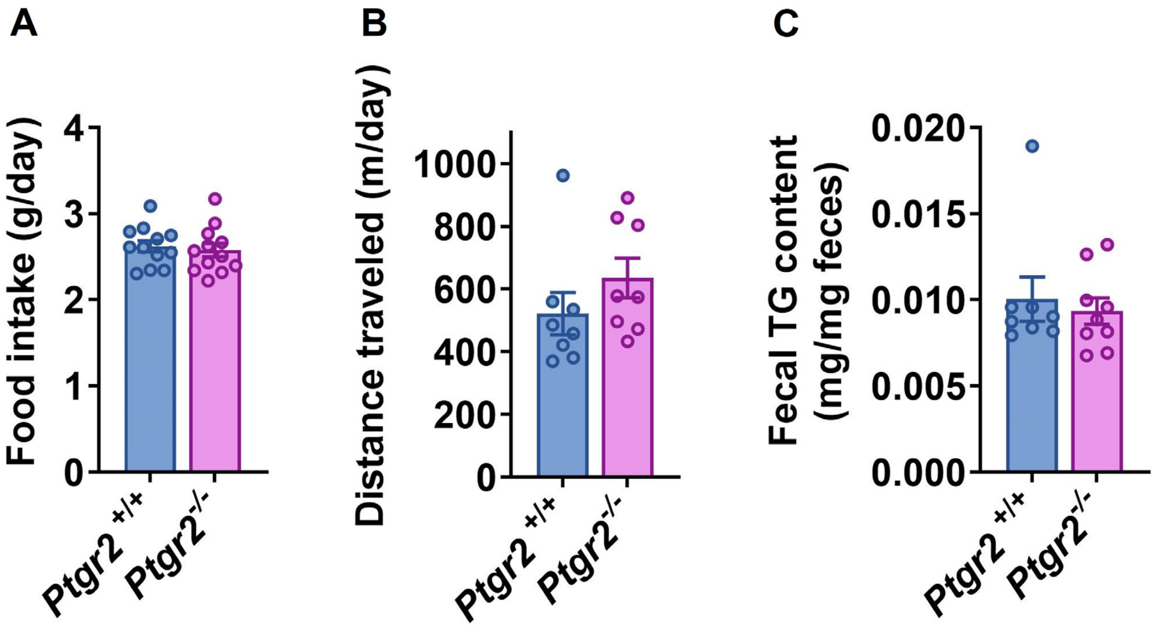

**Appendix Figure S9.** (A) Food intake (n=12:12), (B) distance traveled (n=8:8), (C) fecal triglyceride content (n=8:9) of *Ptgr2*$^{-/-}$ and *Ptgr2* $^{+/+}$ mice on HFHSD. Data information: All data are presented as mean and standard error (S.E.M.). Statistical significance was calculated by two-sample independent *t*-test. Source data are available online for this figure.

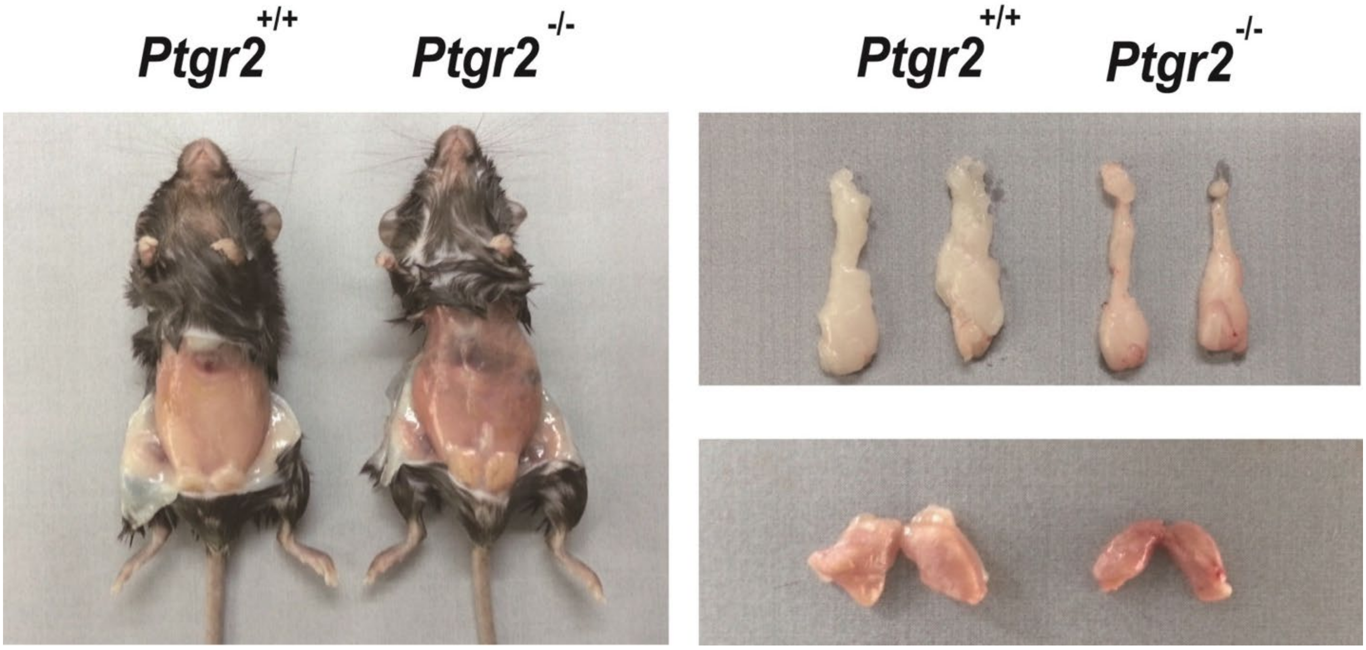

**Appendix Figure S10.** Gross appearance (left panel), perigonadal fat (right upper panel), and brown fat (right lower panel) of *Ptgr2*$^{-/-}$ mice and *Ptgr2*$^{+/+}$ mice on high-fat high-sucrose diet.

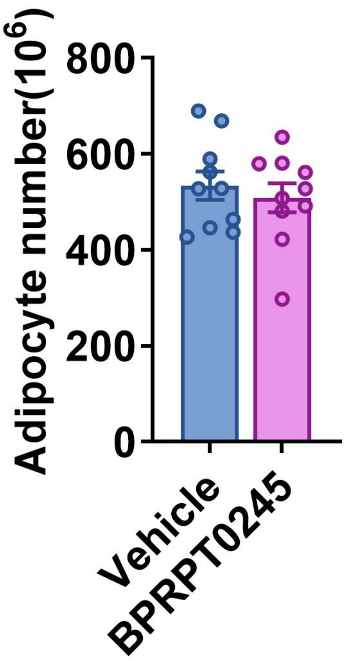

**Appendix Figure S11.** The adipocyte number of perigonadal fat in mice receiving BPRPT0245 compared to vehicle**.** Data information: All data are presented as mean and standard error (S.E.M.). Statistical significance was calculated by two-sample independent *t*-test. Source data are available online for this figure.

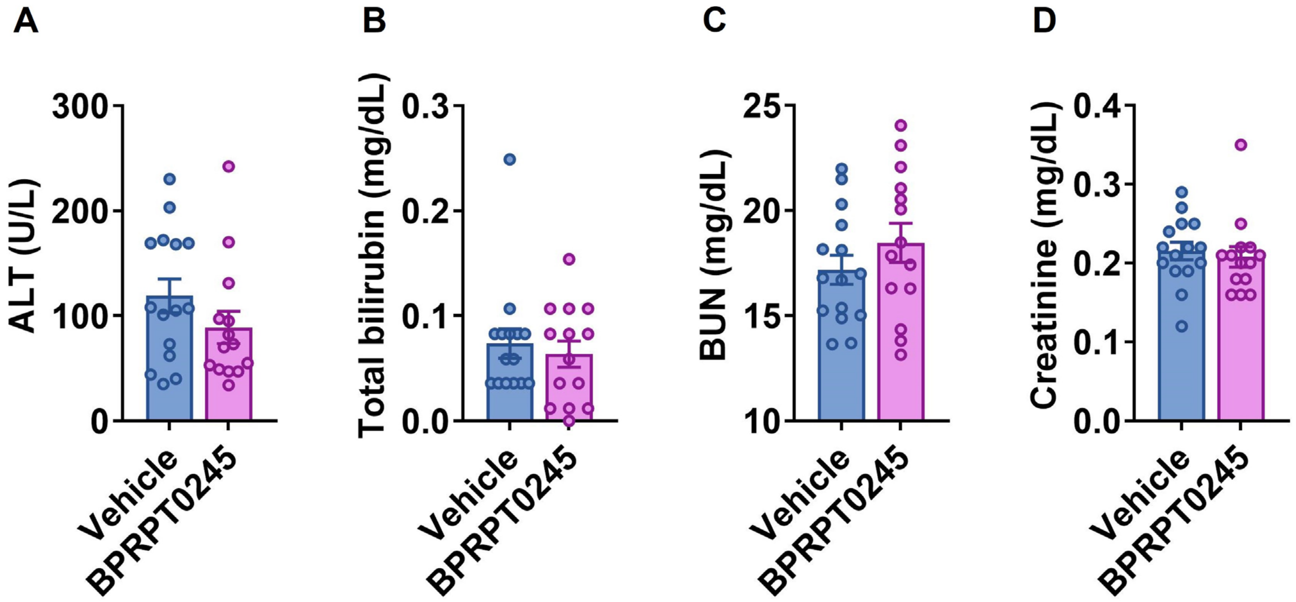

**Appendix Figure S12.** Serum (A) alanine aminotransferase (ALT) (B) total bilirubin (C) blood urea nitrogen (BUN) and (D) creatinine levels of C57BL6/J receiving vehicle and BPRPT0245 (100mg/kg/day) (n=15:14). Data information: All data are presented as mean and standard error (S.E.M.). Statistical significance was calculated by two-sample independent *t*-test. Source data are available online for this figure.

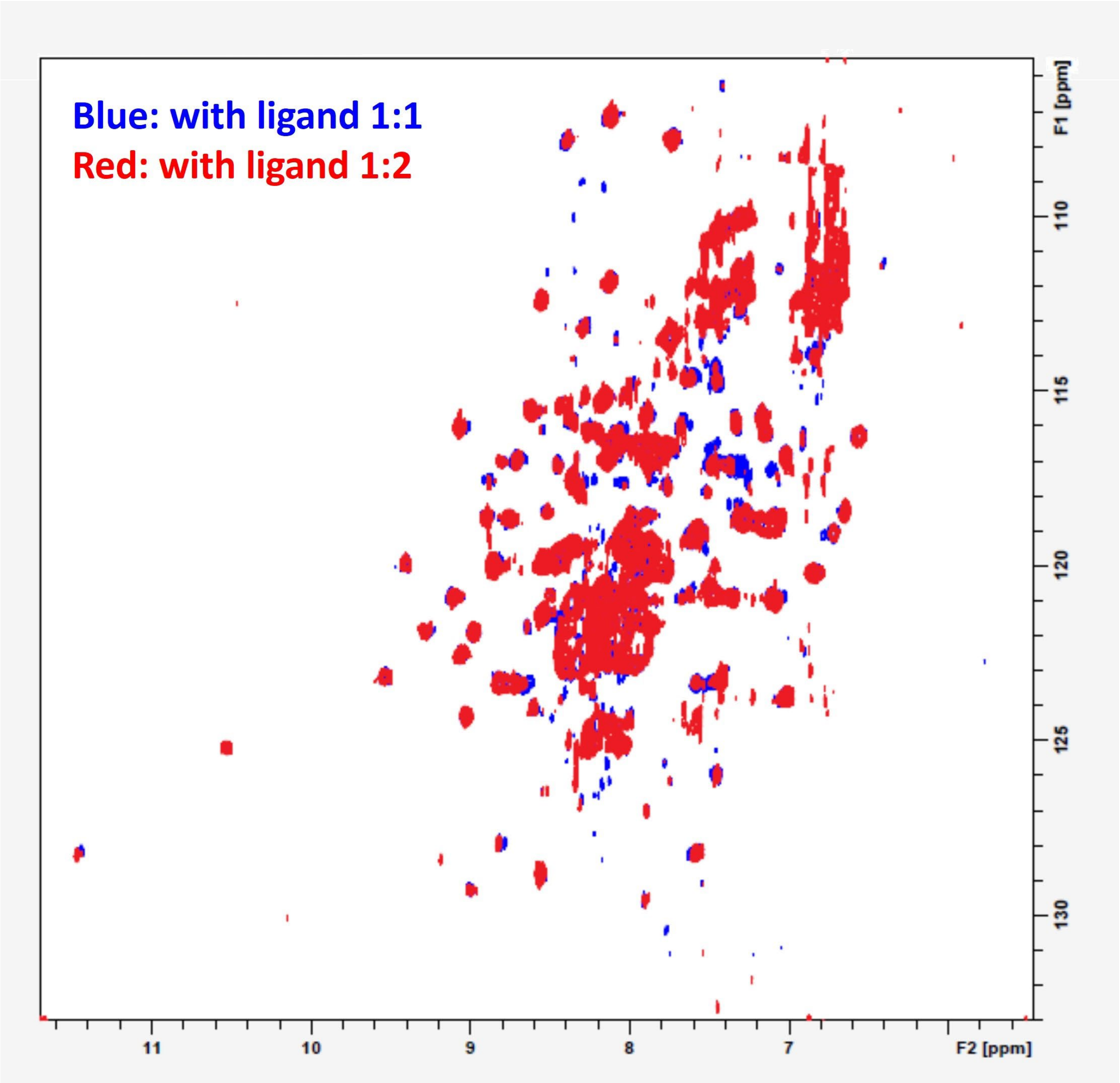

**Appendix Figure S13.** Comparison of the 2D $^{1}H$-$^{15}N$-TROSY-HSQC NMR spectra NMR spectra of 15-keto-PGE2-bound human PPARγ ligand binding domain at two different molar ratios of 15-keto-PGE2 (1:1 vs. 2:1). Source data are available online for this figure.

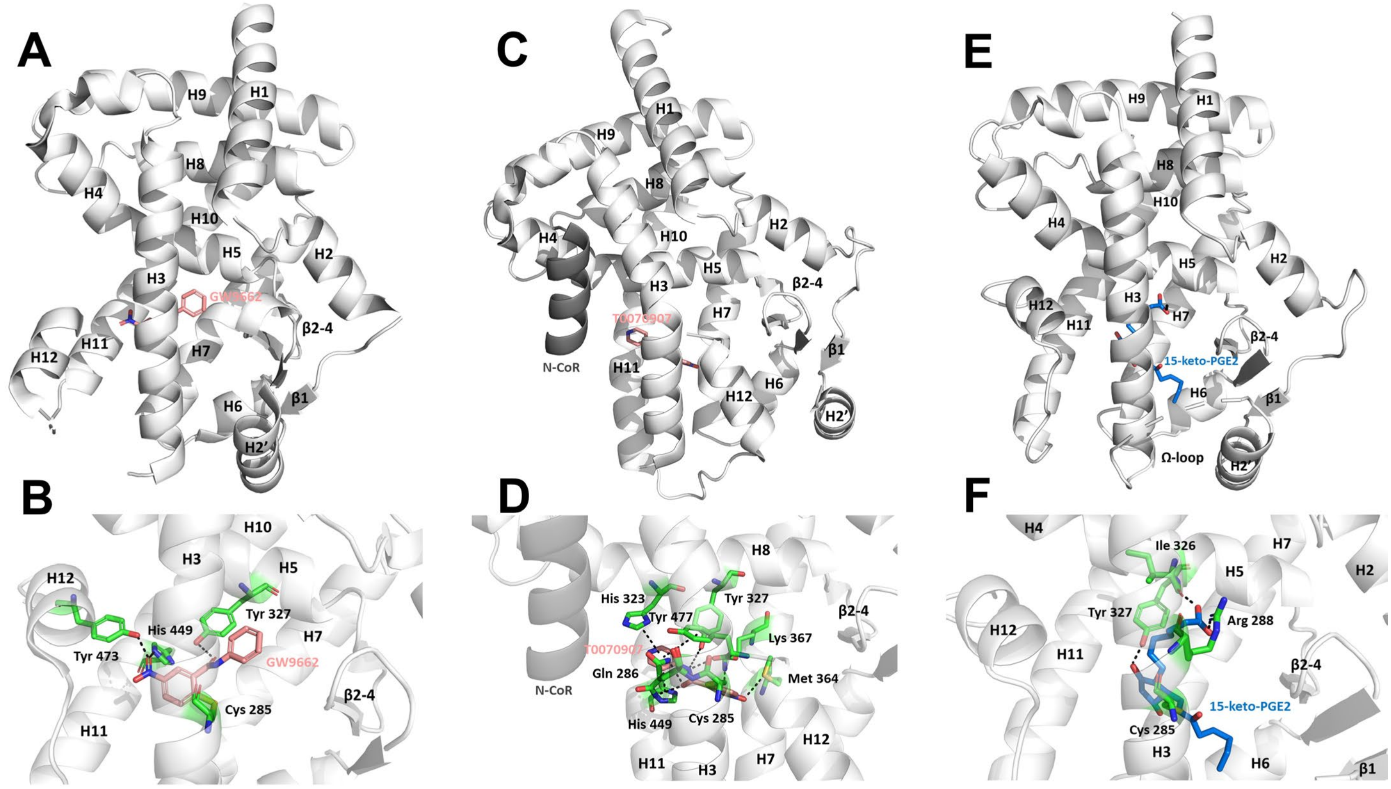

**Appendix Figure S14.** (A)(B) X-ray crystallography showed covalent PPARγ antagonist GW9662 forms a covalent bond with Cys285 (helix 3) and hydrogen bonds with Tyr327 (helix 5), His449 (helix 10) and Tyr473 at helix 12 of human PPARγ (hPPARγ) ligand binding domain (LBD) (PDB: 3B0R chain B). (C)(D) The recruitment of co-repressors such as NCoR1 by T0090907 leads to a unique transcriptionally repressive conformation of the hPPARγ LBD. In this unique conformation, helix 12 was turned into a pocket flanked by helix 3, helix 2', and β-sheets. This helical turn of helix 12 leaves the remaining AF-2 space exposed for more co-repressors binding. In this repressive conformation, an extensive network of interactions between T0070907 and nearby residues to lock helix 12 within the orthosteric ligand-binding pocket (PDB: 6ONI). (E)(F) Molecular docking showed that 15-keto-PGE2 forms a covalent bond with Cys285 and hydrogen bonds with Tyr327 (helix 5), Arg288 (helix 3) and Ile326 (helix 5) of hPPARγ LBD.

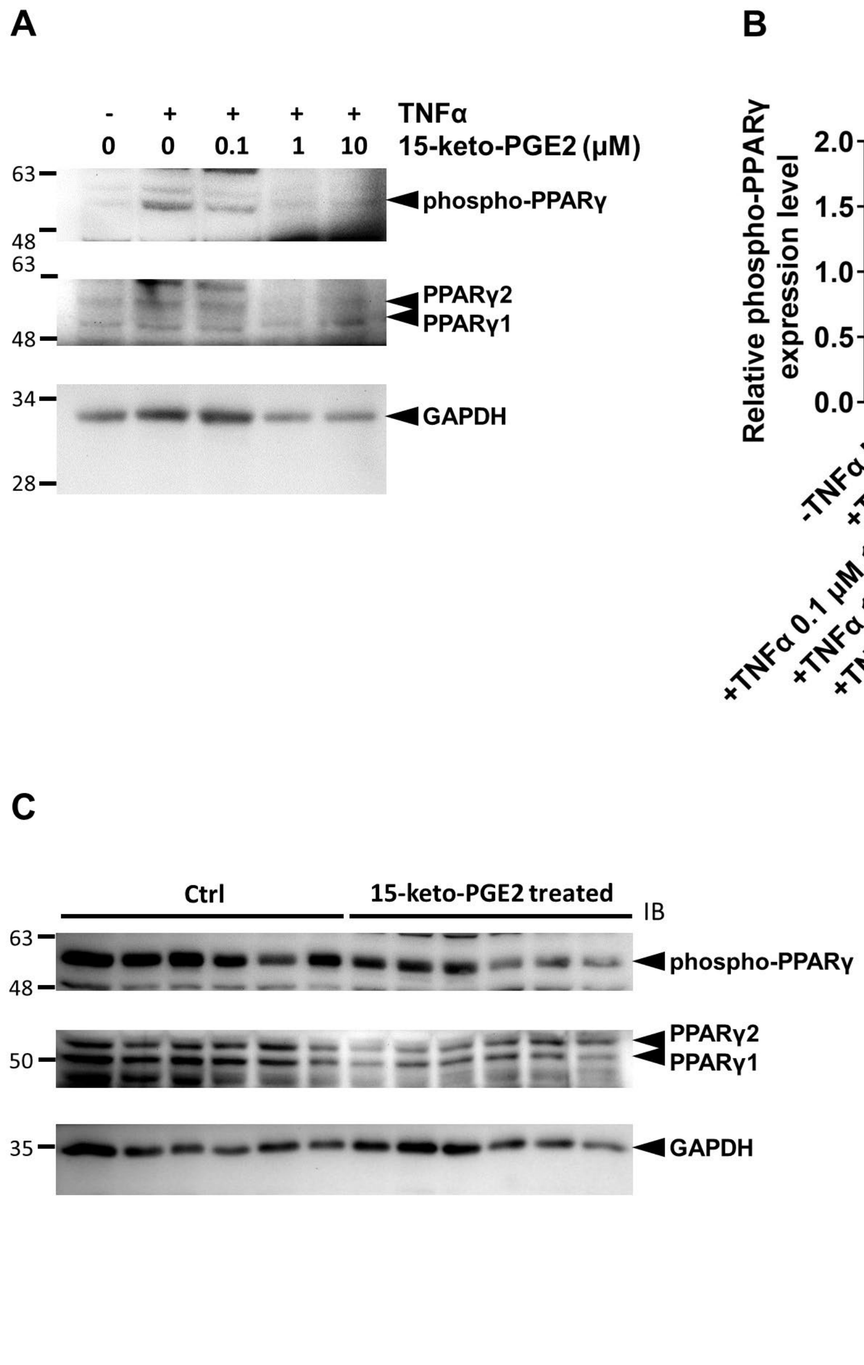

**Appendix Figure S15.** The effect of 15-keto-PGE2 on TNFα-induced phosphorylation of murine PPARγ Ser273 in both (A, B) cultured adipocytes and (C, D) perigonadal adipose tissue of mice.

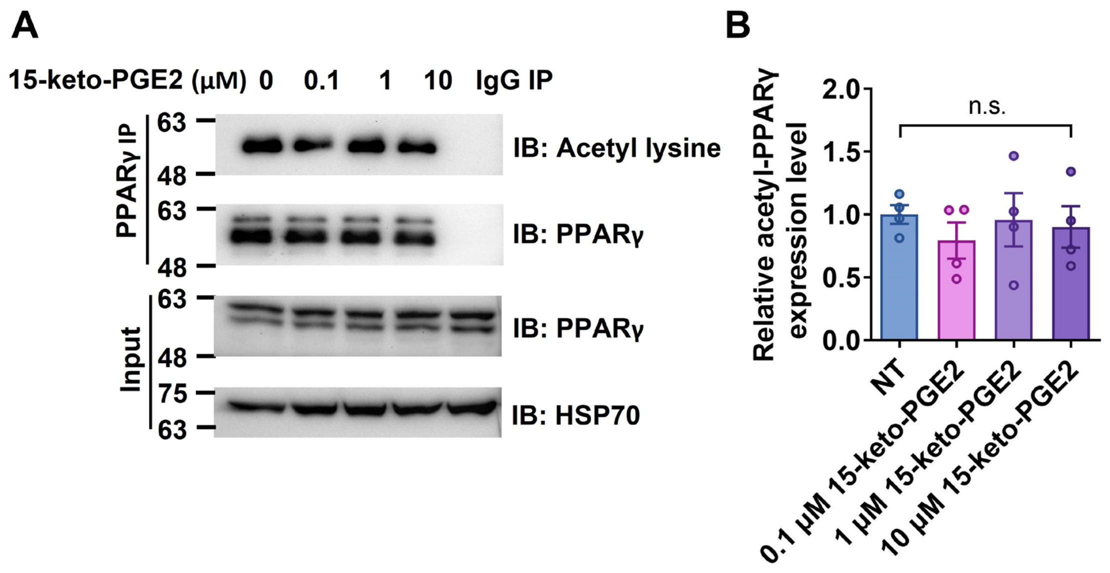

**Appendix Figure S16.** (A, B) The effect 15-keto-PGE2 on lysine acetylation of murine PPARγ in cultured adipocytes. NT: non-treatment; n.s: not significant

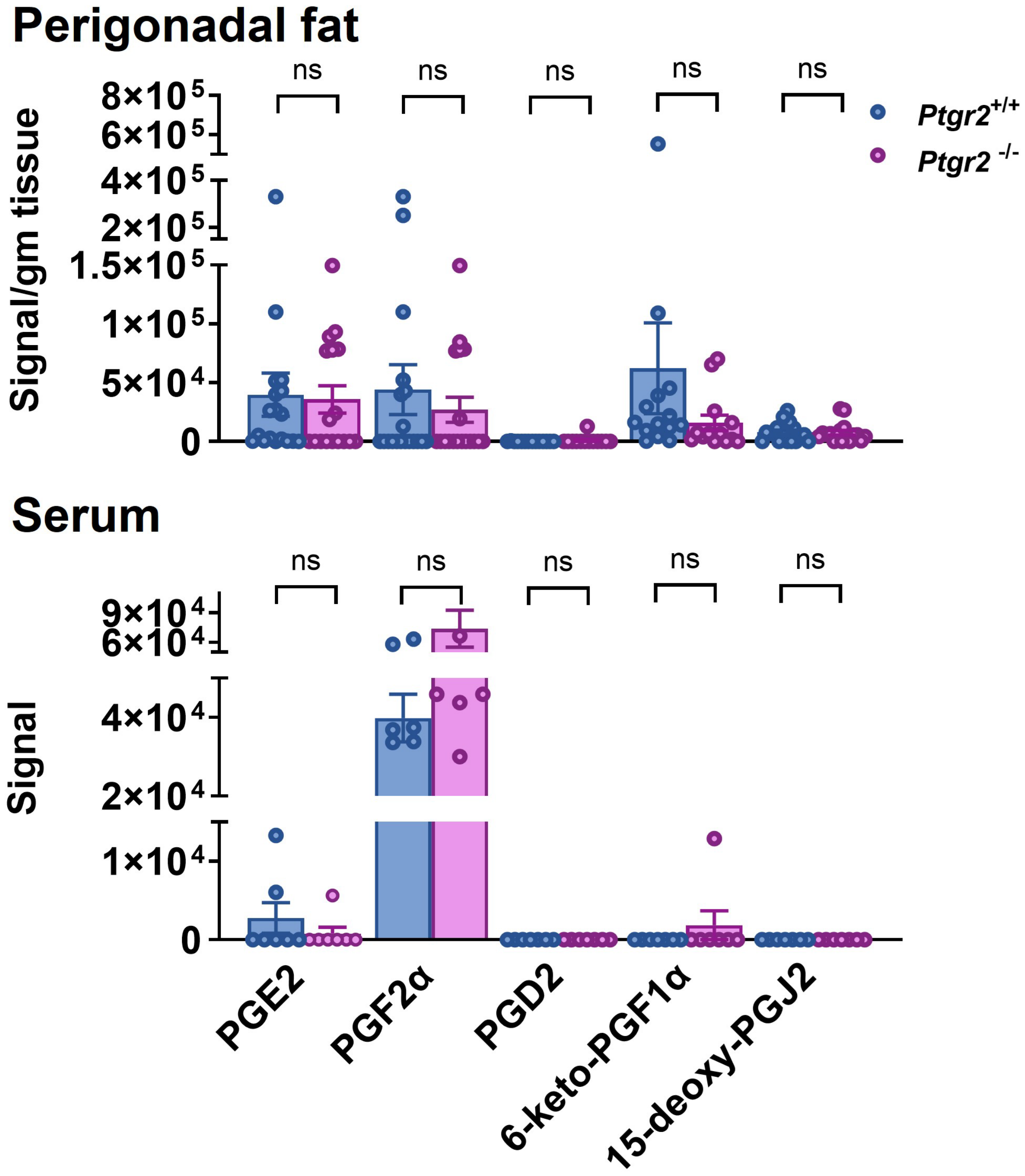

**Appendix Figure S17** Levels of PGE2, PGF2α, PGD2, 15-deoxy-PGJ2, and 6-keto-PGF1α in the perigonadal fat (n=14:13) and serum (n=7:7) of *Ptgr2*$^{+/+}$ and *Ptgr2*$^{-/-}$ mice.

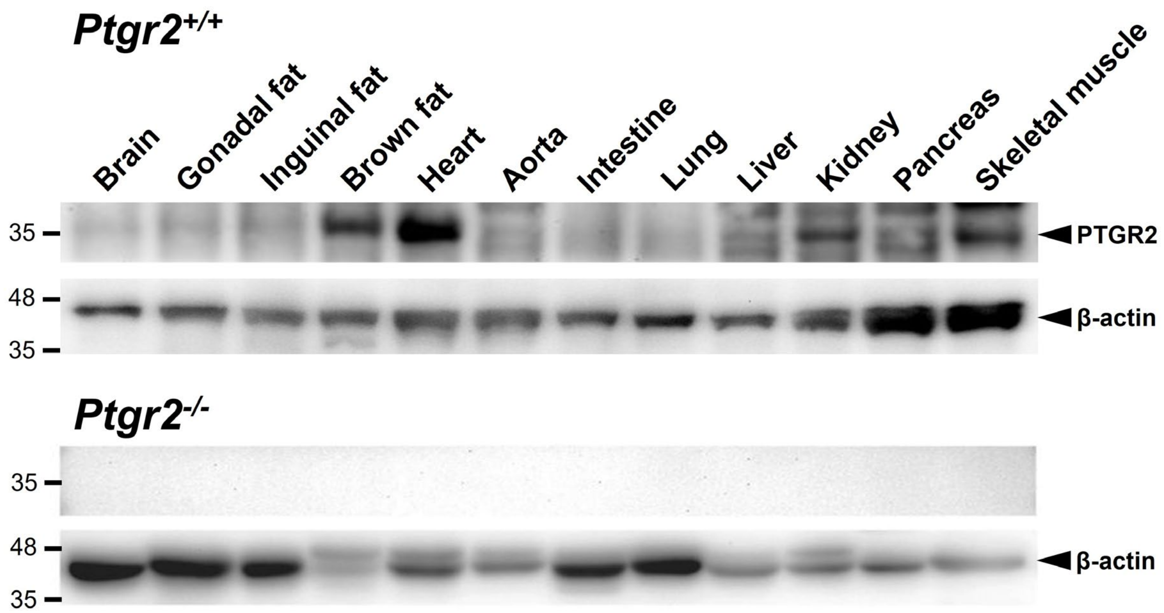

**Appendix Figure S18.** PTGR2 expression in various tissues of *Ptgr2*$^{+/+}$ (upper panel) and *Ptgr2*$^{-/-}$ (lower panel) mice.

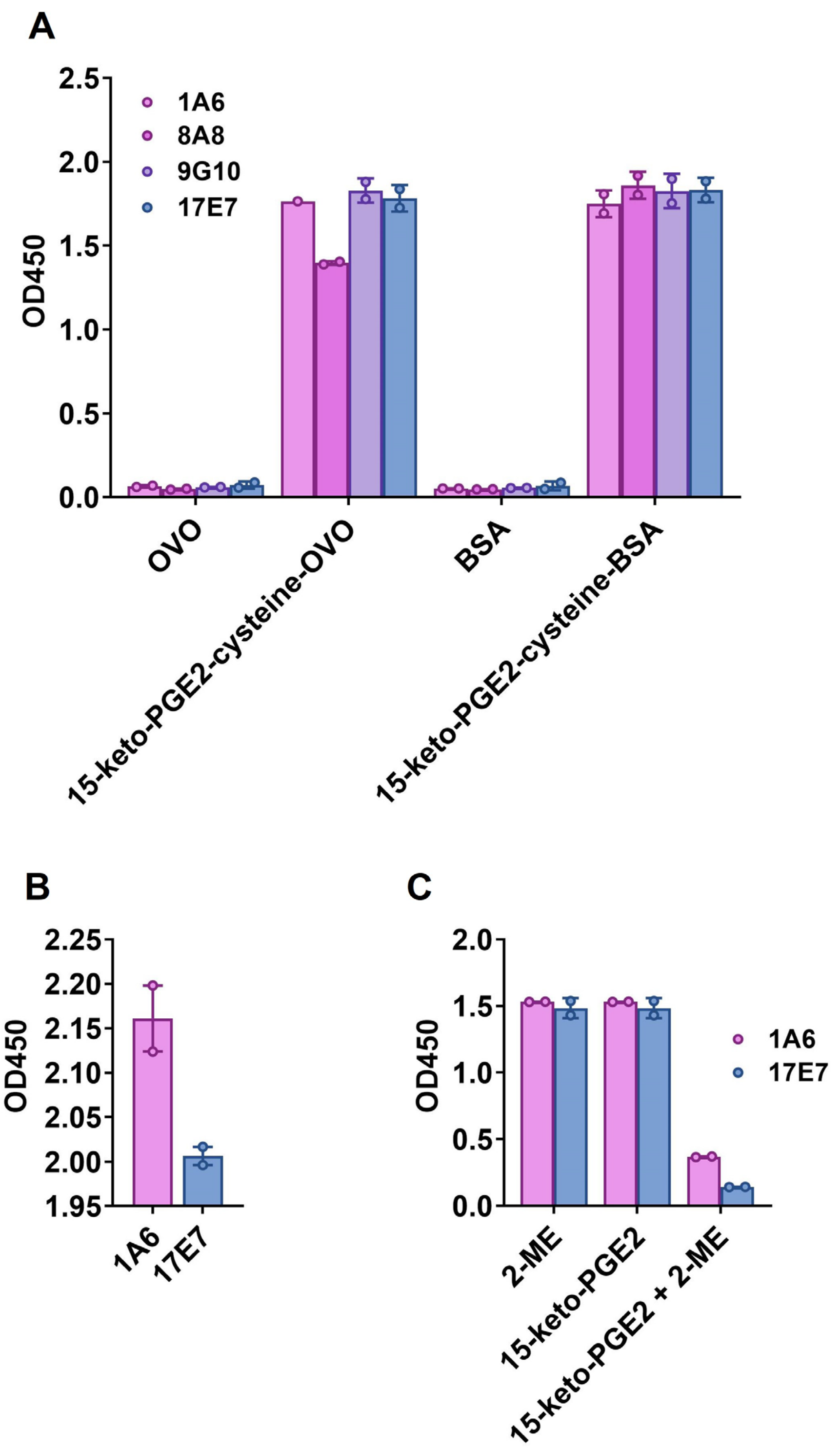

**Appendix Figure S19** Four anti-cysteine-coupled 15-keto-PGE2 hybridoma clones were identified. (A) Clone 1A6 and 17E7 have superior specificity and (B) react only to cysteine-coupled 15-keto-PGE2. (C) The specificity of monoclonal antibodies was examined using ELISA assays. Hybridoma culture supernatants were tested for the reactivity to different antigens coated onto micro-wells. To further test specificity, 10 nM 15-keto-PGE2 was reduced with 2-mercaptoethanol (2-ME). Reduced 15-keto-PGE2 (10 nM), 15-keto-PGE2 (10 nM), and 2-ME (20 nM) were mixed with 1 nM purified antibodies in PBS containing 1% BSA, respectively, and incubated at room temperature for 30 min. The antibody mixtures were then added into wells coated with 15-keto-PGE2-cysteine-BSA for assays.

**Appendix Table S1. Pharmacokinetic study of BPRPT0245 in mice.**

| Mouse IV (dose: 2mg/kg) | | | | Mouse PO (dose: 10 mg/kg) | | | | |
|---|---|---|---|---|---|---|---|---|
| $T_{1/2}$ (hr) | Cl (mL/min/kg) | Vss (L/kg) | AUC (0-inf) (ng/mL*hr) | $T_{1/2}$ (hr) | $C_{max}$ (ng/ml) | $T_{max}$ (hr) | AUC (0-inf) (ng/mL*hr) | F(%) |
| 2 | 79.5 | 5 | 480 | 2.5 | 398 | 0.4 | 725 | 30.2 |

$T_{1/2}$: half-life, Cl: clearance, Vss: steady-state volume of distribution, AUC: area under curve, $C_{max}$: maximum serum concentration, $T_{max}$: time take to reach $C_{max}$, F(%): bioavailability. (Unpublished data)

**Appendix Table S2. Pathological examination of major organs between mice receiving BPRPT0245 and vehicles**

| **Group** | **Control** | | | | |
|---|---|---|---|---|---|
| **Animal ID** | **C1** | **C2** | **C3** | **C4** | **C5** |
| **Brain, Forebrain** | X | X | X | X | X |
| **Brain, Midbrain** | X | X | X | X | X |
| **Brain, Cerebellum and Pons** | X | X | X | X | X |
| **Heart** | | | | | |
| Inflammatory cell infiltration/Cardiomyopathy | 1 | 1 | 1 | 1 | 1 |
| Vacuolation, cardiomyocyte | 1 | 1 | 1 | - | 1 |
| **Lung** | | | | | |
| Inflammatory cell infiltration, perivascular | 2 | 2 | 2 | 1 | X |
| Alveolar epithelium hyperplasia | - | - | 2 | - | X |
| Chronic inflammation, focal | - | 1 | - | 1 | X |
| **Liver** | | | | | |
| Steatosis | 3 | 3 | 3 | 3 | 4 |
| Inflammation | 2 | 1 | 2 | 2 | 2 |
| Lymphocytic cell infiltration, perivascular | - | 2 | 2 | - | 1 |
| Hepatocellular adenoma, multiple | - | - | + | - | - |
| **Kidney** | | | | | |
| Tubular degeneration/regeneration with inflammatory cell infiltration | 2 | 2 | 2 | 2 | 2 |
| Inflammatory cell infiltration, peripelvic /perivacscular | 2 | 2 | 2 | 2 | 2 |
| Hyaline cast, renal tubule | 1 | - | 1 | 1 | 1 |
| Cytoplasmic vacuolation, renal tubule | 2 | 2 | 2 | 2 | 2 |
| Mineralization, renal tubule | 1 | - | 1 | - | - |
| **Pancreas** | | | | | |
| Atrophy and accumulation adipocyte, acinar cell | - | - | 4 | - | - |
| Pancreatic islets hyperplasia | - | - | - | 2 | - |
| Lymphocytic infiltration, perivascular | 1 | 1 | 1 | 2 | - |
| Fat infiltration, exocrine | 2 | - | - | 2 | 1 |
| **Brown fat (largely in dorsal intrascapular region)** | | | | | |
| Increased adipocyte size | + | - | + | + | + |
| **White fat (Perigonadal fat)** | | | | | |
| Inflammatory cell infiltration | 1 | - | 1 | - | 1 |
| **Bone marrow, Sternum/ Femur** | X | X | | | |
| Mineralization, focal | | | 2 | - | - |

| | | | | | |
|---|---|---|---|---|---|
| Cellularity, decreased, bone marrow | | | - | 2 | 1 |

| **Group** | **Test** | | | |
|---|---|---|---|---|
| **Animal ID** | **T1** | **T2** | **T2** | **T4** |
| **Brain, Forebrain** | X | X | X | X |
| **Brain, Midbrain** | X | X | X | X |
| **Brain, Cerebellum and Pons** | X | X | X | X |
| **Heart** | | | | |
| Inflammatory cell infiltration/Cardiomyopathy | 2 | 1 | 2 | 2 |
| Vacuolation, cardiomyocyte | - | - | - | 2 |
| **Lung** | | | | |
| Inflammatory cell infiltration, perivascular | 1 | 1 | 2 | 2 |
| Alveolar epithelium hyperplasia | 2 | - | - | - |
| Chronic inflammation, focal | 1 | - | 1 | - |
| **Liver** | | | | |
| Steatosis | 2 | 4 | 2 | 4 |
| Inflammation | 1 | 2 | 1 | 2 |
| Lymphocytic cell infiltration, perivascular | 1 | 1 | 2 | 2 |
| Hepatocellular adenoma, multiple | - | - | - | - |
| **Kidney** | | | | |
| Tubular degeneration/regeneration with inflammatory cell infiltration | 2 | 2 | 2 | 2 |
| Inflammatory cell infiltration, peripelvic /perivacscular | 2 | 1 | 2 | 2 |
| Hyaline cast, renal tubule | 1 | - | - | - |
| Cytoplasmic vacuolation, renal tubule | 2 | - | 2 | 2 |
| Mineralization, renal tubule | 1 | - | - | - |
| **Pancreas** | | | | |
| Atrophy and accumulation adipocyte, acinar cell | - | - | - | - |
| Pancreatic islets hyperplasia | 1 | 1 | - | - |
| Lymphocytic infiltration, perivascular | - | - | 2 | 2 |
| Fat infiltration, exocrine | 2 | 1 | 2 | 1 |
| **Brown fat (largely in dorsal intrascapular region)** | | | | |
| Increased adipocyte size | + | + | + | + |
| **White fat (Perigonadal fat)** | X | X | | |
| Inflammatory cell infiltration | | | 1 | 2 |
| **Bone marrow, Sternum/ Femur** | X | | X | X |
| Mineralization, focal | | - | | |

| | | |
|---|---|---|
| Cellularity, decreased, bone marrow | | - |
| Infiltration, lymphocyte | | 1 |

+ = Present; - =Not present; X=Not remarkable lesion. Degrees of lesions were graded histopathologically from one to five depending on severity (1 = minimal (< 1%); 2 = mild (1–25%); 3 = moderate (26–50%); 4 = moderately severe (51–75%); 5 = severe/high (76–100%).

Gross examinations were summarized as follows: (1) smaller than usual size of the pancreas - The lesion was observed in control mouse (ID: C3). Histopathologically, the pancreas showed exocrine atrophy and fat infiltration. (2) pale discoloration of the liver - The lesion was observed in 2 mice of test group and 5 mice of control group. Histopathologically, the liver showed steatosis and inflammation. (3) small hepatic nodules (<3 mm) of the liver - Histopathologically, the liver showed hepatocellular adenoma. The microscopic alterations were presented in Supplementary Table 5 as follows: 1. Histopathology changes related to the experimental outcomes (non-spontaneous lesions)(1). steatosis with inflammation of the liver (fatty change of liver). (2). cytoplasmic vacuolation of the renal tubule. (3). cardiomyocyte vacuolation. (4). fatty infiltration of exocrine pancreas (adipocyte accumulation). (5). diffuse atrophy of exocrine acini and replacement by adipose tissue.(6). Increase adipocyte size of the brown fat (largely in dorsal intrascapular region). 2. Lesions related to the spontaneous lesions. Several microscopic lesions were observed in organs of the submitted mice, but they were considered to be spontaneous developmental and aging lesions or congenital abnormality. It consisted of: cardiomyopathy/inflammatory cell infiltration, inflammatory cell infiltration at perivascular area in the lung, alveolar epithelium hyperplasia, focal chronic inflammation of the lung, lymphocytic cell infiltration at perivascular area in the liver, hepatocellular adenoma, tubular degeneration/regeneration with inflammatory cell infiltration of the kidney, inflammatory cell infiltration in peripelvic area, renal tubular focal mineralization of medulla, hyaline cast formation of the renal tubule, pancreatic islets hyperplasia, lymphocytic infiltration at perivascular area in the pancreas, focal mineralization of the bone marrow (femur) and decreased cellularity of the bone marrow (femur).

In conclusion, several microscopic changes in various tissues of treated and control animals were considered incidental spontaneous and ageing changes, and these were not related to the test article administration.

**Appendix Table S3. RT-qPCR primer sequences**

| Gene | Gene ID | Forward primer (5'-3') | Reverse primer (5'-3') |
|---|---|---|---|
| *Glut4 (Slc2a4)* | 20528 | ACATACCTGACAGGGCAAGG | CGCCCTTAGTTGGTCAGAAG |
| *Irs2* | 384783 | CACAATTCCAAGCGCCACAA | CATCACCTCCTCCCAGGGTA |
| *Sorbs1* | 20411 | TGGACCAGACAAAGACATGGA | TTCCCCTGTAGGAAGACGAGG |
| *Cd36* | 12491 | ATGGGCTGTGATCGGAACTG | GTCTTCCCAATAAGCATGTCTCC |
| *Acs* | 60525 | GCTGCCGACGGGATCAG | TCCAGACACATTGAGCATGTCAT |
| *Cebpα* | 12606 | GAACAGCAACGAGTACCGGGTA | GCCATGGCCTTGACCAAGGAG |
| *Adipq* | 11450 | TCCTGGAGAGAAGGGAGAGAAAG | CCCTTCAGCTCCTGTCATTCC |
| *Ppia* | 268373 | GCATACGGGTCCTGGCATCTTGTCC | ATGGTGATCTTCTTGCTGGTCTTGC |

**Appendix Table S4. Primers for site-directed mutagenesis**

| | **Sequence (5’-3’)** |
|---|---|
| Forward | CATCCGAATTTTTCAAGGGGCCCAGTTTCGATCCGTAGAA |
| Reverse | TTCTACGGATCGAAACTGGGCCCCTTGAAAAATTCGGATG |

**Appendix Table S5. Sequence of sgRNAs**

| | **Sequence (5'-3')** |
|---|---|
| sgRNA1 | CGGCTTCTACGGATCGAAAC |
| sgRNA2 | GCACCCTTGAAAAATTCGGA |

### References

Lee, M.A., Tan, L., Yang, H., Im, Y.G., Im, Y.J. (2017). Structures of PPARγ complexed with lobeglitazone and pioglitazone reveal key determinants for the recognition of antidiabetic drugs. Sci Rep. 7, 16837.

Lu J, Chen M, Stanley SE, Li E. (2008) Effect of heterodimer partner RXRalpha on PPAR gamma activation function-2 helix in solution. *Biochem Biophys Res Commun*. 365(1):42-6

Ouyang X, Zhou S, Su CT, Ge Z, Li R, Kwoh CK (2013). CovalentDock: automated covalent docking with parameterized covalent linkage energy estimation and molecular geometry constraints. J Comput Chem. 34(4):326-36.

Rigsby RE, Parker AB (2016). Using the PyMOL application to reinforce visual understanding of protein structure. *Biochem Mol Biol Educ.*10;44(5):433-7.

Shackelford C, Long G, Wolf J, Okerberg C, Herbert R (2002) Qualitative and quantitative analysis of nonneoplastic lesions in toxicology studies. *ToxicolPathol* 30(1):93-6.

Urso R, Blardi P, Giorgi G (2002) A short introduction to pharmacokinetics. *Eur Rev Med Pharmacol Sci* 6(2-3):33-44.
